# Nanoclustering of a plant transcription factor enables strong yet specific DNA binding

**DOI:** 10.1101/2025.11.05.686732

**Authors:** Kasper Arfman, Bas P.J. Janssen, Robin Romein, Sophie van den Boom, Maarten van der Woude, Lucas Jansen, Maartje Rademaker, Jorge Hernández-García, João Jacob Ramalho, Melissa Dipp-Álvarez, Jan Willem Borst, Dolf Weijers, Carlo P.M. van Mierlo, Joris Sprakel

## Abstract

Transcription factors (TFs) are traditionally depicted as monomeric, or oligomeric, units that recognize and bind specific DNA sequences to regulate transcription. Emerging evidence suggests that many TFs can undergo liquid phase separation, resulting in condensates that bind DNA through a different mechanism. For the Auxin Response Factors (ARFs), canonical plant TFs, evidence for both scenarios exists. But which of these scenarios is operational in the plant nucleus is unclear. Here, we demonstrate using MpARF2 of *Marchantia polymorpha* that a third scenario is operational: MpARF2 forms nanoscopic clusters under physiological conditions. Nanoclusters combine DNA-binding features that cannot be accessed by the other two scenarios: high-affinity, switch-like, and sequence-specific. Our results suggest nanoclustering as a mechanism that equips TFs with the DNA-binding properties necessary for transcriptional regulation.

**Teaser:** Transcription factor nanoclusters balance the DNA-binding trade-off between weakly binding oligomers and nonspecific condensates.

## Introduction

Transcription factors (TFs) are proteins that bind to specific DNA sequences to regulate gene expression (*1*–*3*). The classical picture of TF binding is that of a monomeric or small oligomeric structure that binds to a specific DNA sequence through structural recognition of a DNA-binding domain (DBD) (*4*–*6*) to activate or repress the transcription of a particular gene or set of genes (*7*). The other domains in a TF are typically not associated with DNA binding but act as modules for homo- or hetero-interactions, for example to recruit accessory proteins needed for transcriptional regulation (*8*).

A contrasting picture has recently begun to emerge; many transcription factors are found to form clusters (*i*.*e*. high-order assemblies) that exhibit enormous multivalency in their binding to DNA (*9*–*19*). These clusters are often described as biomolecular condensates formed through liquid-liquid phase separation (LLPS) (*20, 21*). The formation of multimeric, and in some cases even microscopic, structures is associated with the presence of intrinsically disordered regions (IDR) (*22*–*25*) and can have a drastic impact on the DNA binding mechanism (*26*–*28*). While the monomeric picture relies on structural recognition between the DBD and a regulatory element in the DNA, liquid condensates bind DNA through a physical wetting process (*26*). Although functions for the formation of large TF structures, such as condensates, have been proposed (*29*), their physiological relevance and precise biological function remain elusive.

These recent discoveries have led to a dichotomy. Studies that emphasize the classical picture, often from the viewpoint of the precise targeting of a DBD, do not suggest a need for the formation of higher-order structures. Yet, studies focusing on higher-order TF structures are beginning to highlight their prevalence but leave their function and relevance unanswered. Specifically, the *in vivo* occurrence of TF condensates is open to debate, as many observations rely on non-physiological conditions, such as strong overexpression of the TF, to report on a process that is by nature strongly concentration dependent. To reconcile these perspectives, two key questions must be addressed: (i) what structural form do TFs take under native conditions in the cell nucleus, and (ii) by what mechanism do these structures bind genomic DNA?

In this paper, we explore these questions in the context of the Auxin Response Factors (ARFs), a canonical family of plant TFs (*30*–*33*). Transcription factors in the ARF family, which are deeply conserved across all land plants and even their algal sister clades (*34*), regulate gene expression in response to the plant hormone auxin. In addition to a DBD, which encodes sequence-specific binding to Auxin Responsive Elements (AuxREs) (*5*), many ARF proteins contain an intrinsically disordered middle region (MR), and a C-terminal Phox and Bem1 (PB1) multimerization domain. The DBD is reported to form dimers in solution and on DNA (*35*) and disrupting DBD-DBD interactions between ARFs results in strong defects (*5*), indicating that these interactions are essential for ARF function. The MR and PB1 domains both appear to play a role in the formation of higher-order structures (*14*).

For ARFs, evidence exists to support both pictures outlined above. In line with the classical picture, binding of ARFs to AuxREs is specific (*36*) and appears in previous work to occur in a dimeric configuration (*5, 35, 37*). However, these conclusions are based on measurements performed on short dsDNA oligomers with two AuxREs in tandem (*5, 37*–*39*). These oligomers are designed to accommodate dimers, but their short length precludes the binding of higher-valency structures. Moreover, in a biological context, ARFs may interact with DNA regions flanking the recognition motif; however, the use of oligomeric DNA artificially eliminates the possibility of such nonspecific interactions. Therefore, oligomers cannot be used to determine whether the DNA-binding structure is larger than a dimer.

In contrast, ARFs can also form microscopic condensates *in vivo* (*14, 40*), with each condensate containing many protein molecules. By analogy to other condensate-forming TFs, this could result in completely different DNA binding mechanisms and affinities (*26*). However, ARF condensates have only been observed in plants that either overexpress the ARF protein or contain a transgene that expresses a fluorescently tagged copy of the ARF gene from a non-native genomic location (*14, 40*). Consistent with this concern, various studies report that higher-order assemblies of other TFs also depend strongly on the level of overexpression, with droplets appearing only when overexpression is sufficiently strong (*41*–*43*). Moreover, ARF condensates have been observed in the cytoplasm where they cannot interact with genomic DNA (*14, 40*). Given these shortcomings in the reported observations of ARF condensates, their physiological relevance is questionable.

We use the repressor MpARF2 from the liverwort *Marchantia polymorpha* as a model (*44*–*47*) to explore the formation of higher-order assemblies of this TF and how this impacts DNA binding. We find that, at native nuclear concentrations, MpARF2 forms nanoscopic clusters, both *in vivo* and *in vitro*. These nanoclusters are significantly larger than a protein dimer but much smaller than the previously observed condensates and form a thermodynamic state distinct from phase-separated condensates. Although MpARF2 does form condensates *in vitro*, these only appear at protein concentrations well above the physiological range. And while condensates can bind DNA, their binding is sequence-independent when a sufficiently long DNA template is used, which is incompatible with the biological function of TFs. Instead, we show how nanoclusters, that feature a size-limiting mechanism similar to micelles, turn the gradual and low-affinity binding of monomeric ARFs into a high-affinity switch-like binding transition in the physiological range, while retaining strong sequence specificity. We conclude that, at physiological concentrations, MpARF2 monomers and dimers may not bind DNA, and condensates do not exist, but that micelle-like nanoclusters are the relevant regulatory entity that equip ARFs with DNA binding properties beneficial for transcriptional regulation.

## Results

### MpARF2 forms nanoclusters *in vivo* at native expression levels

To explore the assembly state of our TF, MpARF2 (Fig. S1), we first realize that the association of proteins is strongly concentration dependent. Hence, to arrive at an understanding of ARF assembly and DNA binding that is physiologically meaningful, we first need to define the native biological context by studying the appearance of MpARF2 in its native cellular context (Fig. 1A).

**Fig. 1.**
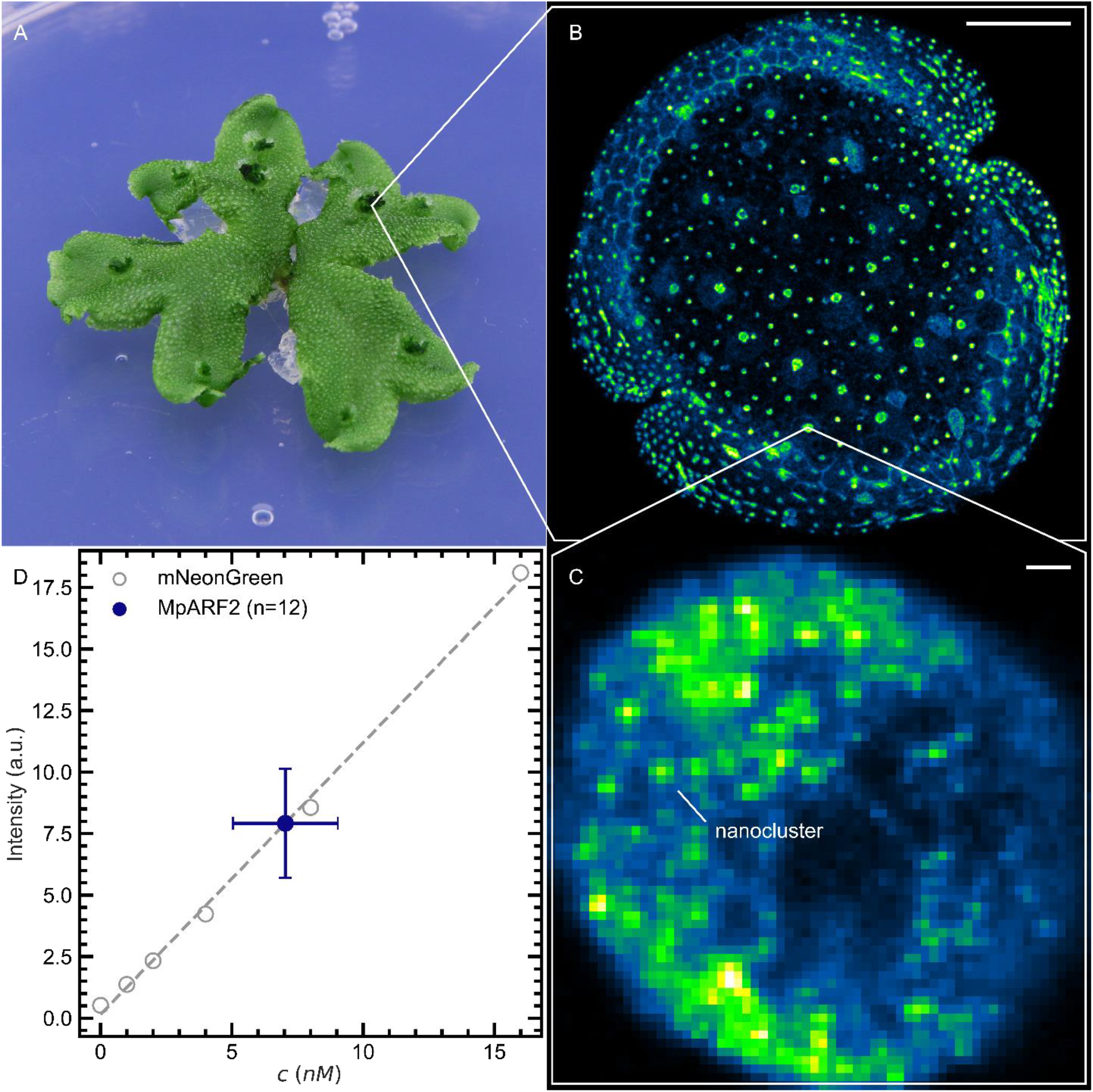
MpARF2 forms nanoclusters at native expression levels *in vivo*. (**A**) *M. polymorpha* knock-in plant expressing MpARF2-mNeonGreen at native levels. (**B**) Confocal fluorescence image of an entire gemma showing MpARF2 localization in nuclei. Scale bar: 100 µm. (**C**) High-resolution confocal image of a single nucleus showing nanoclusters at native expression. Scale bar: 1 µm. (**D**) The MpARF2 concentration in nuclei (n=12) was estimated at 7 ± 2 nM (mean ± SD, filled symbol) by fluorescence intensity calibration with purified mNeonGreen solutions of known concentration (open symbols).

We make use of the recently developed genomic MpARF2-mNeonGreen knock-in lines (*47*). In these lines, a fluorescent mNeonGreen protein is fused to the C-terminus of MpARF2 in its native genomic locus. Since increased or decreased MpARF2 levels show dramatic developmental defects (*47*), and MpARF2-mNeonGreen knock-in plants are normal, we can infer (i) that the fusion protein is biologically active and (ii) that it accumulates at (or nearly at) native concentrations. We imaged gemmae, the asexual propagules of *M. polymorpha*, by confocal microscopy (Fig. 1B) and find that MpARF2 is present in nuclei as small punctae in a background of low fluorescence intensity (Fig. 1C). The punctae appear as diffraction-limited objects, implying they are of nanoscopic dimensions (*i*.*e*. below 200 nm in diameter) and we observe that their fluorescence intensity is weak. We introduce the term *nanocluster* to refer to a nanoscopic protein assembly without implying any specific molecular identity. The small size and low brightness of the nanoclusters suggests that they contain a rather limited number of MpARF2 molecules. These assemblies are larger than monomers or dimers yet much smaller than the liquid-like droplets that are observed in overexpression lines of ARFs (*40*) or other TFs (*48*–*50*). The nanoclusters show limited mobility in the nucleus (Movie S1), suggesting that they are anchored to immobile matrix components, likely genomic DNA.

To enable performing *in vitro* experiments at physiologically relevant concentrations, we determined the concentration at which MpARF2 is present in nuclei. We first calibrated the optical response of our confocal fluorescence microscope by imaging solutions of purified mNeonGreen at known concentrations, resulting in a linear calibration curve (Fig.1D, open symbols). Next, using identical experimental settings, we imaged nuclei in rhizoid precursor cells in *M. polymorpha* gemmae. From the fluorescence intensity averaged over entire nuclei, we estimate the native nuclear MpARF2 concentration to be 7 ± 2 nM (Fig. 1D, closed symbol). Although expression levels of this protein likely vary across tissue types and developmental stage, this provides an estimate of the biologically relevant concentration range for this protein.

### MpARF2 forms nanoclusters *in vitro* at physiological concentrations

To explore the phase behavior of MpARF2 we purified recombinant MpARF2. We observed that large protein assemblies formed during purification, which were difficult to separate from monomeric protein (Fig. 2A). These observations indicate that MpARF2 has a strong intrinsic tendency to form larger structures. This tendency became more apparent when we performed Dynamic Light Scattering (DLS) on the protein immediately after its elution from a Size-Exclusion Chromatography (SEC) column. The protein fraction that eluted as a monomer was diluted to 50 nM in the same buffer and, within minutes, the solution reached a state consisting of a mixture of monomers and distinct nanoclusters of ∼150 nm in diameter (Fig. 2B, solid line).

**Fig. 2.**
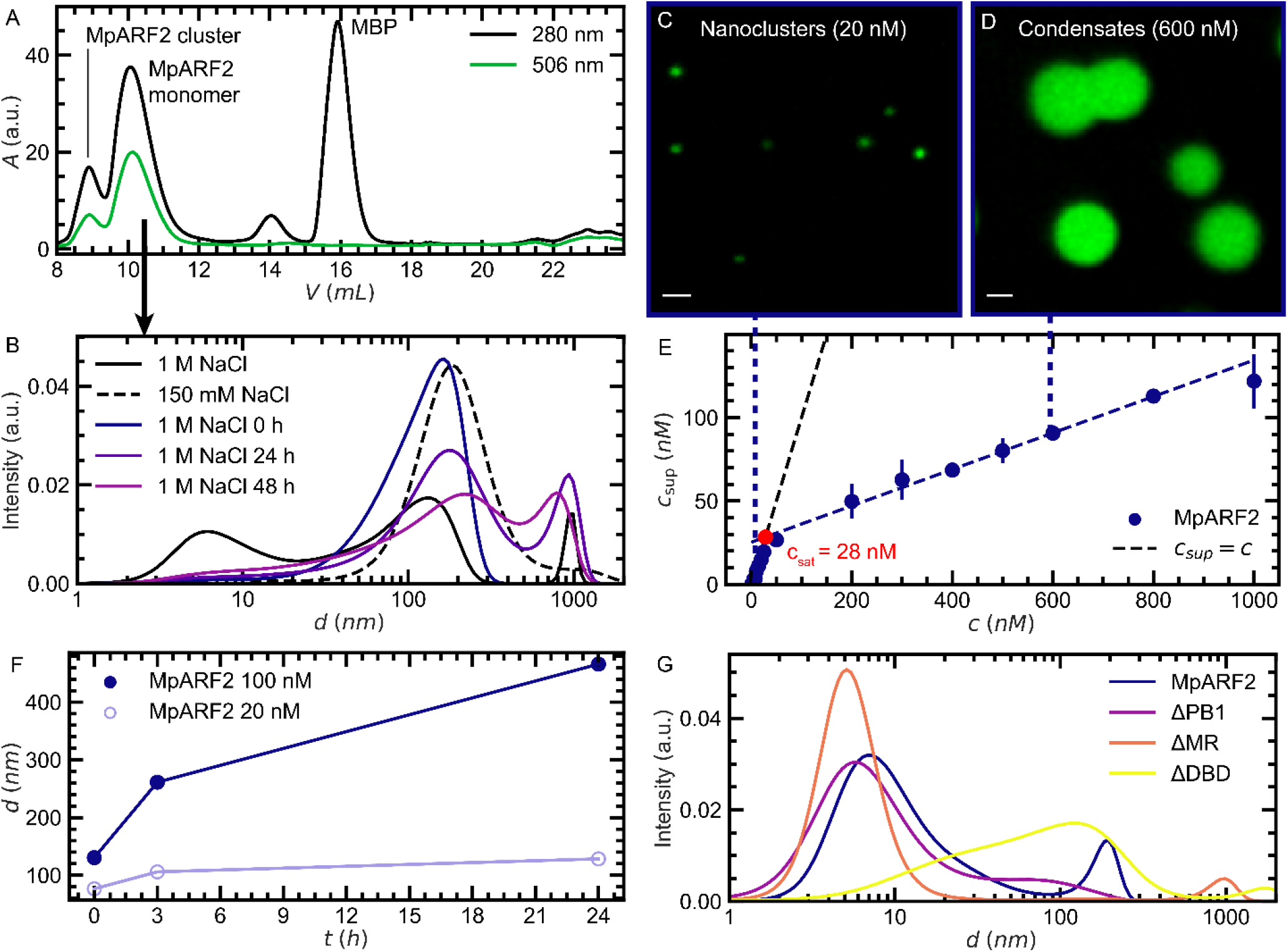
MpARF2 forms nanoclusters and condensates *in vitro*. (**A**) SEC chromatogram as the final step in the purification of MpARF2, after removal of the affinity/solubility tag MBP; mNeonGreen absorbs at 506 nm. (**B**) Dynamic Light Scattering (DLS) size distributions of the MpARF2 monomer fraction after sequential salt treatments: starting with 1 M NaCl (solid black line) down to 150 mM (dashed line) and readjusting to 1 M (colored-line time series). (**C-D**) Confocal fluorescence images of MpARF2 nanoclusters (C) and condensates (D; scale bar: 1 µm). (**E**) Centrifugation assay in which the protein concentration in the supernatant after centrifugation, *c*_sup_, is measured as a function of the total protein concentration. The onset of condensate formation is estimated at *c*_sat_ ≈ 28 nM. (**F**) Assembly sizes, determined by DLS, show that condensates (100 nM) grow substantially whereas nanoclusters at 20 nM remain nanoscopic in size. (**G**) DLS size-distributions for domain-deletion variants of MpARF2. The MR is the main driver of clustering, the PB1 enhances clustering, and the DBD confers a defined cluster size.

Of note is that the size distribution is distinctly bimodal, which indicates that nanoclusters have a well-defined size, while intermediate-sized assemblies (*i*.*e*. between monomers and nanoclusters) are disfavored. The size distribution at high salt shows a small peak for monomers and a larger peak for nanoclusters. However, the DLS results are presented as intensity-weighted rather than number-weighted distributions. Since in light scattering the scattered intensity is proportional to the diameter of the object to the sixth power, these distributions are heavily biased towards larger objects. Based on our data, we estimate that at high salt, the number ratio between monomers and nanoclusters is 10^8^: 1. Hence, at high salt, the solution is primarily monomeric. However, lowering the salinity to a more physiological level (150 mM NaCl) at 50 nM MpARF2 induced nanocluster formation to such an extent that the monomer population could no longer be detected by DLS (Fig. 2B, dashed line). To assess whether nanocluster formation is reversible, we induced clustering by lowering the salinity (150 mM NaCl) and then reverted the sample back to high salt conditions (1 M NaCl, 50 nM MpARF2). This revealed a gradual and partial dissociation of nanoclusters, but the signal of monomers was not recovered (Fig. 2B).

Because nanoclusters form rapidly and reversibly, even at low concentrations, these data suggest that MpARF2 exists in a dynamic equilibrium between nanoclusters and low-order oligomers under physiologically relevant conditions. This implies that MpARF2 cannot be maintained as a pure monomer *in vitro*, but it also means that such a state is unlikely to occur naturally *in vivo*. Small oligomers and nanoclusters coexist almost unavoidably in solution, and this behavior mirrors our *in vivo* observations of small clusters superimposed on a diffuse background of weak fluorescence (Fig.1C).

Confocal microscopy on purified MpARF2 solutions confirms that at physiological protein concentrations, in the low nM range, nanoclusters appear as diffraction-limited spots (Fig. 2C), which is consistent with our *in vivo* observations at native protein levels (Fig. 1C). However, at much higher concentrations, MpARF2 forms microscopic condensates, recognizable by their spherical shape and their ability to coalesce (Fig. 2D; Movie S2), similar in appearance to what has been observed for AtARF19 in *Arabidopsis thaliana* (*14, 40*). Moreover, this is also consistent with the prediction by the FuzDrop model (*51*) that MpARF2 shows a 99.8% probability of spontaneous liquid-liquid phase separation (Fig. S2).

### MpARF2 nanoclusters are distinct from condensates

AtARF19 has been shown to form microscopic condensates in plant cells at high expression levels (*14*), and its domain architecture is homologous to that of MpARF2. This raises the question whether the nanoclusters we observe both *in vivo* and *in vitro* are small condensates or a different type of structure. To determine the protein concentration at which condensates form, we performed a centrifugation assay that allows condensates to be gravitationally separated from nanoclusters and small oligomers. After centrifugation, we measure the protein concentration remaining in the supernatant, *c*_sup_, as a function of the total protein concentration prior to centrifugation. In the absence of condensate formation, *c*_sup_ equals the total protein concentration. Thermodynamics dictates that condensates form as a first-order phase transition, the onset of which is marked by a threshold concentration, *c*_sat_, at which the solution has become saturated with dissolved protein (*52*). Indeed, we determine the formation of condensates as a distinct departure from the linear trend at *c*_sat_ ≈ 28 nM (Fig. 2E). Nanoclusters remain in the supernatant after centrifugation under subsaturated conditions, whereas they are depleted from solution once concentrations exceed *c*_sat_ (Fig. S3). This suggests that nanoclusters phase-transition into condensates at *c*_sat_ and thus represent a distinct type of assembly.

For condensates formed by LLPS, the thermodynamic equilibrium state would be one single condensate that minimizes the interfacial area between the two phases. Therefore, condensate droplets are expected to continuously grow by coalescence and Ostwald ripening (*53*). Indeed, using fluorescence imaging, we found that MpARF2 solutions above *c*_sat_ exhibit continuous condensate growth over time. However, below *c*_sat_, where we observe only nanoclusters, we find virtually no change in size or fluorescence intensity over 24 h (Fig. 2F; Fig. S4). Hence, we conclude that the nanoclusters we observe below *c*_sat_ are not condensates.

If these MpARF2 nanoclusters are not condensates, then what are they? To address this question, we investigated the nanoclustering mechanism using domain deletions. A complete loss of clustering was obtained for MpARF2 in which the intrinsically disordered middle region was removed (ΔMR); hence, the MR is essential for nanocluster formation (Fig. 2G, orange line). Moreover, we find that MR deletion increases the threshold concentration for condensation *c*_sat_ from 28 nM to 1.5 µM (Fig. S5). Purified MR forms condensates on its own with *c*_sat_ ≈ 200 nM (Fig. S6) and forms nanoclusters in subsaturated solutions as low as 15 nM (Fig. S7). Therefore, the MR domain is necessary and sufficient for nanoclusters to form at biologically meaningful protein concentrations. The Phox and Bem1 (PB1) domain mediates multimerization via self-complementary charged interactions (*54*). Deleting PB1 significantly reduces clustering, although some nanoclusters were still observed (Fig. 2G, purple line). Thus, PB1 appears to enhance the clustering that is driven by the MR. By providing additional multivalent interactions, PB1 likely creates local spots of increased MpARF2 concentration where cluster formation is nucleated. Based on these observations, one may conclude that MR and PB1 are responsible for nanocluster formation, with the DBD primarily effectuating DNA binding. However, our results suggest something different. Deleting the DBD domain leads to very strong clustering, exceeding that of the full-length protein, but these clusters lacked a defined size (Fig. 2G, yellow line). This suggests that the DBD acts as a growth-limiting module: a behavior strongly reminiscent of micellization, where the combination of a phase separation tendency with a growth-limiting mechanism leads to the formation of nanoscopic structures of well-defined size, typically containing tens to hundreds of monomers.

The notion of a micellar structure is consistent with our observations of a bimodal size distribution, as micelles coexist with monomers under thermodynamic equilibrium. The growth-limiting mechanism in micelles typically originates from the formation of an outward-facing corona, such as the charges on surfactant molecules. In this view, DBDs might form an outer shell that stabilizes the micelle-like nanoclusters. Supporting this idea, AlphaFold-Multimer predicts that MpARF2 pentamers (the largest supported multimer in this model) adopt a conical shape, with DBDs at the base and MRs at the tip (Fig. S8). An arrangement of outward-facing DBDs could also be beneficial for the DNA-binding function of TF nanoclusters.

Another defining feature of micelles is the presence of a Critical Micelle Concentration (CMC). The CMC marks a concentration threshold below which micelles are absent, and above which micelles form abruptly with their number increasing linearly with concentration. To test whether MpARF2 follows this description, we quantified cluster abundance as a function of protein concentration using single-particle tracking (SPT). Indeed, MpARF2 nanoclustering commences abruptly at a critical concentration of °0.5 nM, and the number of clusters scales linearly with concentration beyond this threshold (Fig. S9). Together, these observations demonstrate that MpARF2 nanoclusters in subsaturated solutions have many of the defining features of a micelle.

### Higher-order MpARF2 assemblies gain DNA-binding affinity but lose specificity

Above we have established that MpARF2 forms nanoclusters under physiological protein and salt concentrations, both *in vitro* and *in vivo*. However, it remains unclear how the formation of these structures impacts DNA binding. Through single-molecule studies using immobilized oligos, the dissociation constant, *K*_d_, of MpARF2-DBD to short oligomeric DNA with a tandem AuxRE was previously estimated at 61 nM (*55*). Since the nuclear concentration of MpARF2 that we measure is much lower, it appears that under physiological conditions in the nucleus, virtually no monomer or dimer would be bound to DNA.

However, by featuring many DBDs in one entity, nanoclusters have the potential to interact with DNA with a high valency. In multivalent interactions, weak binding affinities of monomers are exponentially enhanced. AuxREs are sparse in genomic DNA. Hence, the number of *specific* DNA interactions that may form is likely not limited by cluster size, but rather by the prevalence of recognition elements in the DNA that are within reach of a single cluster. Clustering should therefore not substantially increase the number of specific DNA contacts that can form but greatly enhances the likelihood of nonspecific interactions. The DNA-binding domains of ARFs are highly sequence specific (*38*). To achieve such specificity, nonspecific DNA interactions must be very weak, potentially negligible, compared with binding to an AuxRE. Hence, nonspecific interactions have little effect on monomeric DNA binding. However, for multimers, given that the binding affinity scales exponentially with valency, many nonspecific interactions may collectively contribute significantly to DNA binding.

To test this hypothesis, we conducted Electrophoretic Mobility Shift Assay (EMSA) experiments in which protein-DNA mixtures are separated by gel electrophoresis, allowing free and bound states to be spatially resolved and quantified using fluorescence imaging (Fig. 3A). The experimentally determined bound fraction of DNA, *θ*_DNA_, plotted as a function of protein concentration, produces a binding curve from which an effective dissociation constant, 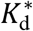, of the interaction may be estimated. The true *K*_d_ may be overestimated in EMSA because the system is not at equilibrium during electrophoresis: bound protein-DNA complexes can dissociate while the gel is running, but reassociation is limited as the unbound species become physically separated. EMSA was originally developed to study the lac repressor transcription factor (*56*), which does not contain an intrinsically disordered region (IDR). However, it is predicted that ∼90% of TFs contain IDRs (*57*), and the presence of an IDR is one of the hallmark features of phase-separating proteins (*58*). Protein clustering may complicate the EMSA experiment, as large protein assemblies bound to DNA may not be able to penetrate the nanoporous gel and thus become trapped in the loading well. Fluorescence signals coming from wells may therefore be unavoidable, but excluding them from analysis, as is common practice, would introduce a bias.

**Fig. 3.**
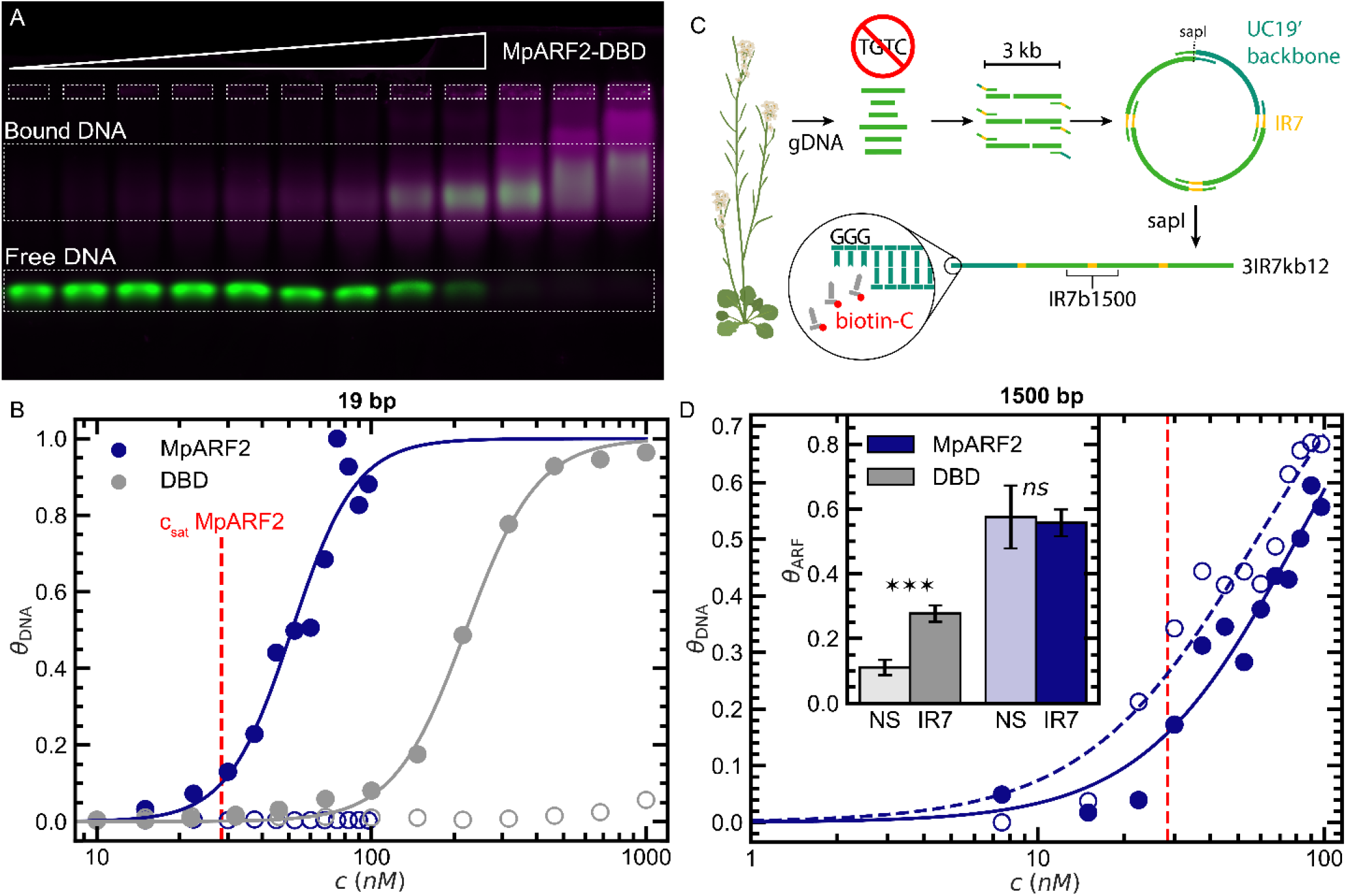
Clustering and DNA accessibility define DNA-binding. (**A**) Result of an EMSA assay to quantify binding of MpARF2-DBD to oligomeric dsDNA, showing both DNA (green) and protein fluorescence (magenta, log color scale). (**B**) DNA binding curves of full-length MpARF2 (blue) and its DBD (gray) to 19 bp DNA containing the IR7 recognition sequence (filled symbols) or lacking the TGTCNN motif (open symbols). (**C**) Approach for obtaining long DNA templates containing IR7 motifs surrounded by nonspecific DNA. *A. thaliana* gDNA fragments without TGTCNN motifs were inserted into a pUC19 backbone, with IR7 being introduced via primer overhangs. The resulting plasmid was linearized and biotinylated for single-molecule studies (3IR7kb12) or PCR amplified for EMSA (IR7b1500). (**D**) MpARF2 binding to 1500 bp DNA containing IR7 (filled) or a nonspecific variant lacking TGTCNN (NS, open symbols), showing that sequence preference is lost. Inset: concentration-averaged fractions of bound protein, *θ*_ARF_, showing that the DBD retains sequence-specificity for 1500 bp DNA, whereas MpARF2 does not. \*\*\**P*<0.001; *ns, P*>0.05; Welch’s t-test.

To assess DNA binding in the absence of extensive clustering, we first performed EMSA on MpARF2-DBD with 19 bp oligonucleotides containing two AuxREs in inverted repeat with 7 bp spacing (IR7b19; Fig. 3A). MpARF2-DBD exhibits a moderate affinity for IR7b19 with an effective 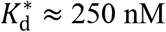. Moreover, we find that this interaction is sequence specific: MpARF2 shows virtually no binding to a non-specific DNA template of the same length (NSb19) that lacks the TGTCNN recognition sequence (Fig, 3B, gray; Fig. S10). Hence, classical TF binding behavior is observed when clustering is suppressed.

To study how TF clustering can affect DNA binding, we repeated the experiment using full-length MpARF2. As before, DNA binding to oligomeric DNA was highly sequence-specific. However, consistent with our earlier observations on AtARF2 (*37*), the full-length protein has a substantially higher affinity 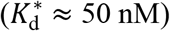 compared to MpARF2-DBD, already revealing how DNA-binding thermodynamics can be controlled by multimerization (Fig. 3B, blue; Fig. S11).

The use of short oligos is standard practice in EMSA and in the study of interactions between ARFs and DNA. However, we suspected that the use of rather short DNA oligos could impose an experimental bias, since TF clusters have the capacity to occupy many more binding sites than a short DNA template can provide. In fact, the oligonucleotides most commonly used in previous studies are just long enough to include a tandem AuxRE, such as IR7 (19 bp), and can accommodate at most two DBDs (*59*). In their native cellular context, ARF proteins interact with genomic DNA of which accessible regions vary in length but are often much longer than 19 bp (*60*). To explore the scenario of a longer stretch of accessible DNA, we engineered 1500 bp DNA fragments of two types: one version containing a single IR7 AuxRE in the center (IR7b1500), and a nonspecific version (NSb1500) free of the recognition motif (Fig. 3C; Table S1).

MpARF2-DBD binds 1500 bp DNA in the same way it binds 19 bp oligomers: visual inspection of the gels reveals that binding remains sequence specific and relatively weak (Fig. S12). However, this experiment could not be analyzed using conventional EMSA quantification, because the 1500 bp DNA is roughly twenty times heavier than the DBD molecule. Consequently, monomeric or dimeric binding does not cause a measurable DNA mobility shift. To quantify DNA-binding, we computed the bound fraction of protein (instead of DNA), *θ*_*ARF*_, averaged across protein concentration to improve statistics. This analysis confirms that MpARF2-DBD remains sequence-specific even for long DNA templates. For full-length MpARF2 however, extending the DNA template from 19 bp to 1500 bp resulted in a complete loss of sequence specificity (Fig. 3D, inset).

Like the DBD, full-length MpARF2 is much lighter than the 1500 bp DNA template, yet, in contrast to the DBD-only experiments, binding of MpARF2 caused the DNA signal to shift completely into the wells (Fig. S13). This pronounced shift indicates that, whereas MpARF2-DBD binds as low-order assemblies (likely dimers), full-length MpARF2 binds 1500 bp DNA exclusively in assemblies of at least tens to hundreds of protein molecules. The shift observed only for full-length MpARF2 allowed the binding curve to be constructed from *θ*_DNA_ as before. The transition of this curve coincides with *c*_sat_, suggesting that condensates, rather than nanoclusters, are responsible for the DNA binding observed here (Fig. 3D). In the condensate regime, the length of DNA thus imposes a tradeoff between the affinity and specificity of DNA binding. However, this is problematic: sequence specificity is a crucial trait of TFs, so our results seem to be at odds with biological reality.

Our EMSAs highlight the DNA-binding behavior of condensates formed at relatively high TF concentrations. But what about nanoclusters present at significantly lower and biologically relevant protein concentrations? We would like to emphasize that EMSA may fall short in detecting DNA binding of nanoclusters due to limitations related to stoichiometry and sensitivity. In an EMSA, the amount of DNA used determines the maximum observable binding fraction. For example, at a 100-fold excess of DNA, the proportion of bound DNA can at most be 0.01 if all protein molecules are bound to DNA in a monomeric state. Clustering amplifies this stoichiometry effect even further: because clusters contain many individual MpARF2 molecules, the concentration of clusters can only be a small fraction of the total MpARF2 concentration.

Nanoclusters are a low-abundance species (*c*_cluster_ ≪ *c*_protein_) present only at the low end of our protein concentration range (*c*_protein_ ≪ *c*_DNA_). Therefore, their binding to DNA can be expected to have a negligible effect on the total proportion of bound DNA. To restore this stoichiometry imbalance, the DNA content would have to be drastically reduced. However, doing so would compromise the detectability of the already faint DNA bands. In short, EMSA is not a suitable technique to probe the low nM binding of TF clusters due to intrinsic technical limitations. Hence, an alternative approach is required that combines high sensitivity with the ability to operate at low DNA concentrations.

### Nanoclusters offer strong, sequence-specific, switch-like DNA binding at physiological levels

To investigate DNA binding of MpARF2 nanoclusters, we employed a single-molecule imaging approach that integrates microfluidics with total internal reflection fluorescence (TIRF) microscopy. In this assay, custom-engineered DNA molecules of 12 kb in length, containing three equally spaced IR7 sites (3IR7kb12; Fig. 3C), are tethered to the surface of a passivated microfluidics chip through biotin-neutravidin interactions. The DNA was stained with SYTOX Orange (SxO), allowing individual DNA molecules to be visualized under mild flow. This flow stretches the DNA coils into linear strands, facilitating the observation of protein binding (Fig. 4A). For these experiments, MpARF2 proteins, either full-length or domain deletions, were covalently labeled with ATTO633 with a degree of labeling of approximately 1 dye molecule/protein. The proteins were introduced into the flow cell, already containing the surface-bound DNA; protein and DNA channels were imaged in an alternating fashion. We used protein concentrations between 1 – 10 nM, a physiological concentration range well below *c*_sat_, to prevent condensates from being present. This approach allows tens to hundreds of individual DNA molecules to be monitored simultaneously within a single field-of-view, providing a real-time visualization of MpARF2 binding (Fig. 4B).

**Fig. 4.**
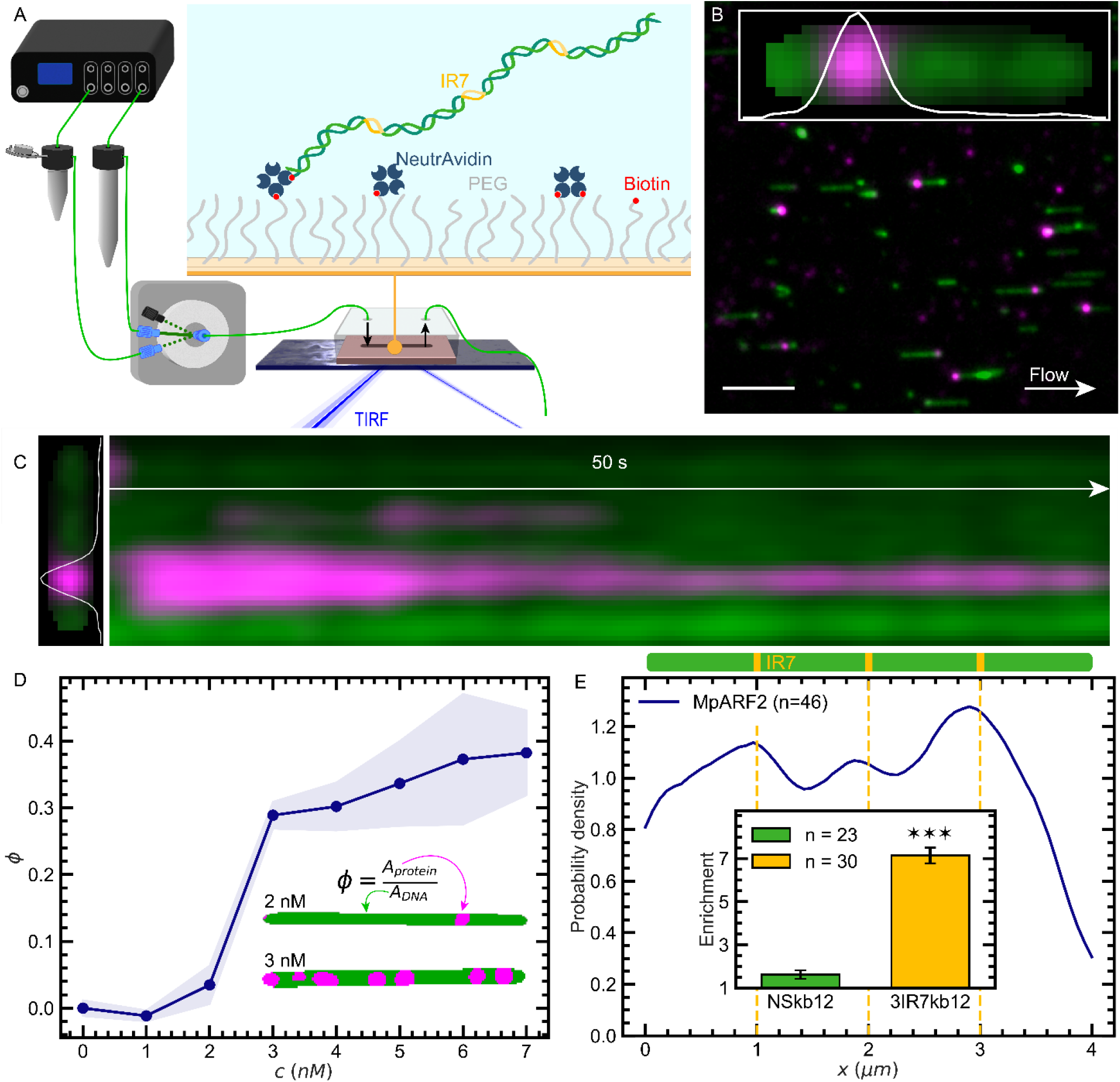
DNA binding of nanoclusters at physiological concentrations is strong, specific and switch-like. (**A**) Illustration of the single molecule imaging assay, which combines microfluidics, surface immobilization and TIRF microscopy. (**B**) Typical field-of-view showing individual flow-aligned DNA strands (green) with MpARF2 nanoclusters (magenta) bound to the DNA templates. [MpARF2] = 1 nM. Scale bar: 5 µm. Inset: filtered Region Of Interest (ROI). (**C**) Kymograph of a nanocluster binding to DNA, illustrating the high temporal stability of a nanocluster once bound, with virtually no lateral movement. (**D**) Binding isotherm of nanoclusters, determined from the area fraction of protein covering the DNA, *ϕ*. Inset: binary-filtered ROIs used in coverage analysis. (**E**) Spatial distribution of binding events along individual DNA strands (solid) compared to the predicted IR7 positions (dashed). Inset: enrichment of protein on nonspecific (green) and IR7-containing (orange) DNA templates, measured as the ratio of protein intensity on DNA compared to the surrounding background. \*\*\**P*<0.001; Welch’s t-test.

Upon introducing full-length MpARF2, we observe small protein clusters, significantly brighter than a single fluorophore, attaching to the DNA. This reveals that nanoclusters bind DNA at physiological protein concentrations (Fig. 4B). Although some nanoclusters also bind to the passivated glass surface, we discriminate these from DNA-bound nanoclusters by their positional fluctuations that follow the waving motion of the DNA strands to which they are bound (Movie S3). DNA binding is observed at MpARF2 concentrations as low as 1 nM, suggesting that nanoclusters have a strong affinity for DNA, well below reported *K*_d_ values for binding of monomeric/dimeric species to short oligomers (*55*). Due to the high affinity and multivalent interaction, nanoclusters binding events are stable and typically remain bound for minutes without noticeable motion of the nanoclusters along the DNA (Fig. 4C). While nanoclusters form high-affinity and time-stable bonds with DNA, monomeric or dimeric ARFs bind much weaker and exhibit fast dissociation dynamics (*59*).

Above we reported that in the absence of DNA, clusters of full-length MpARF2 form spontaneously in solution, but that this relies on the presence of a PB1 domain that acts as a concentrator. Here, our observations further support that a concentrator is needed. For full-length MpARF2, DNA binding occurred instantaneously (< 100 ms), suggesting that nanoclusters pre-form in solution rather than assembling on DNA. In contrast, the ΔPB1 protein variant typically first binds to DNA and then progressively grows into a cluster, rather than forming a cluster in bulk solution (Fig. S14). The EMSA data provide complementary evidence: at ∼10 nM, ΔPB1 can bind DNA as small oligomers without producing a mobility shift, whereas larger protein clusters dominate at elevated protein concentrations (Fig. S15). Together, this suggests that when the PB1 domain is present, clustering occurs primarily in bulk. In the absence of this domain, clustering in the bulk solution is suppressed, consistent with the hypothesized role of the PB1 domain as a concentrator. However, by acting as a scaffold, DNA can take over the role of the PB1 as a concentrator, thereby forcing cluster formation to occur after binding. This picture is fully consistent with the concept of templated assembly, where multiple binding transitions (MR-MR, PB1-PB1 and DNA-DBD), acting synchronously, orchestrate DNA binding (*61*).

At 1 nM MpARF2, binding events are rare and the number of nanoclusters present in solution is limited. By contrast, at 10 nM, DNA rapidly gets covered by MpARF2 nanoclusters, which are also more abundant in solution. This hints at a narrow binding transition between 1 and 10 nM. To better resolve this binding transition, we conducted our assay on a 48 kb λ-DNA template containing numerous AuxREs and systematically varied the MpARF2 concentration inside the same microfluidics chip. Between measurements, the flow cell was thoroughly washed with buffer to minimize any unwanted time-dependent effects. We quantified the coverage of the DNA with MpARF2, *ϕ*, and observe a surprisingly sharp, almost binary, switch-like binding transition only 2 nM wide (Fig. 4D). This is remarkable, as the binding curves for ARF monomers and dimers span orders of magnitude (*37*). We also note that the DNA coverage plateaus below full coverage, as would be expected for a sequence-specific DNA-binder to DNA that contains both specific and nonspecific regions.

In our EMSA assays, we observed that the formation of large protein assemblies (*i*.*e*. condensates) led to a loss of sequence specificity in DNA binding (Fig. 3D). We hypothesized that, by keeping their size limited, nanoclusters retain their sequence specificity, while still benefiting from the enhanced affinity provided by clustering. To test this hypothesis, we generated a nonspecific DNA variant (NSkb12; Table S1) that lacks the IR7 motif and contains minimal TGTCNN elements. Aside from the absence of the three IR7 motifs, the sequence of NSkb12 is identical to that of 3IR7kb12. We performed single-molecule experiments on both DNA variants using 10 nM MpARF2 and compared the extent of DNA binding. We first select regions of interest that each contain a single DNA strand. For each ROI, DNA fluorescence was used as a mask to distinguish protein signal on the DNA from that outside of it. DNA binding was quantified as the signal enrichment on the DNA (⟨*I*_in_⟩/⟨*I*_out_⟩).

Conducting this assay on NSkb12 DNA revealed that nanoclusters can occasionally interact with nonspecific DNA (Fig. S16); however, such interactions were rare, showing only a modest enrichment of ∼1.5-fold. By contrast, DNA containing three IR7 motifs binds MpARF2 much more strongly, with the enrichment increasing to ∼7. This confirms our hypothesis and reveals that nanoclusters, while binding cooperatively, retain strong sequence-specificity (Fig. 4E, inset).

In fact, sequence specificity was pronounced enough that we could resolve it within a single DNA molecule. When stretched by weak hydrodynamic flow, the three IR7 binding sites on a single 3IR7kb12 DNA strand are physically separated by approximately 1 mm, which is just enough to resolve them optically. We selected 46 ROIs, gathered across 1 – 10 nM protein concentrations, in which DNA binding is clearly visible, and computed the spatial distribution of MpARF2 along the contour of the DNA strand, averaged over time and across ROIs. Indeed, three preferred binding sites emerged, corresponding almost perfectly with the predicted IR7 motif positions within this DNA construct (Fig. 4E).

## Discussion

The results presented above provide a picture of how the plant transcription factor MpARF2 forms nanoscopic assemblies at biologically relevant protein concentrations. These nanoclusters, estimated to contain tens to several hundred protein molecules, exhibit many of the characteristics of a micelle, including a well-defined size corresponding to a state of thermodynamic equilibrium. This distinctly separates them from condensates, which are macroscopic droplets containing thousands to possibly millions of proteins and are thermodynamically unstable; they tend to grow until a single macroscopic condensate remains. MpARF2 also forms these phase-separated condensates *in vitro*, but not at concentrations we encountered in plant nuclei. The nanoclusters of this auxin response factor arise from a balanced interplay between the different protein domains. The middle region, high in intrinsic disorder, is the main driver of nanocluster formation, while the PB1 interaction domain appears to function as a concentrator that promotes protein clustering, and the DNA binding domain appears to function as a growth limiter. This is also exactly what is needed for micelle formation: a driving force for phase separation that is limited to a nanoscopic size by a growth limitation mechanism.

The nuclear MpARF2 concentration range we observed *in vivo* lies within a regime where neither monomers/dimers nor condensates are capable of efficiently binding DNA. The measured MpARF2 levels are below the saturation threshold required for condensate formation, ruling out condensates as the dominant DNA-binding species. Although monomers and dimers are likely present in the nucleus, their weak affinities would preclude stable DNA interactions at physiological protein concentrations. Instead, our data supports a model in which nanoclusters represent the functional DNA-binding form of MpARF2. These assemblies exhibit high-affinity, sequence-specific and stable interactions with DNA, and their binding displays a switch-like dependence on protein concentration, with the transition of this switch occurring withing the physiological MpARF2 range.

Monomers/dimers, nanoclusters and condensates are notably different in their DNA-binding. But which DNA-binding traits are actually favorable from a biological perspective? To address this, it is helpful to consider the conceptual requirements for TFs to accurately regulate gene expression. Their binding must be specific to their target genes, *i*.*e*. they must exhibit sequence specificity. At the same time, their binding is ideally switch-like, so distinct DNA-bound and -unbound states can be achieved. A high affinity is favorable for reasons of protein economy and such that the binding is temporally stable, so that a transcriptional state can be maintained without large fluctuations. And finally, there must be options to toggle this switch-like state. Therefore, the binding must be tunable via a signaling process.

From a physical chemistry perspective, it is evident that monomeric binding cannot meet all needs: while monomers can offer excellent sequence specificity, it is challenging to create interactions that are both high-affinity and tunable. Moreover, the binding isotherms of monomers will be gradual rather than resembling a switch. Macroscopic condensates formed by LLPS, on the other hand, can meet the constraints of switch-like and high-affinity binding, but have little sequence specificity because the background affinity for non-specific DNA is dramatically amplified by the enormous multivalency of a droplet containing thousands of proteins. Our data demonstrate that clustering, by amplifying weak, nonspecific interactions, inherently poses a trade-off between DNA-binding affinity and specificity. Without clustering, DNA binding is specific but weak. The formation of large assemblies, on the other hand, leads to strong but nonspecific binding. Nanoclusters, situated in the middle of these extremes, provide an optimal balance. By coupling phase separation with a growth-limiting process, nanoclusters achieve high-affinity switch-like binding at concentrations within the physiological range, while sequence specificity is retained. It is therefore not surprising that the formation of nanoscopic assemblies is beginning to emerge as a conserved and universal mechanism for DNA- and RNA-binding proteins (*62*–*67*).

However, our data do not fully address how nanoclusters can be toggled between bound and unbound states. Since we observe an almost binary switch-like transition from unbound to bound, changes in TF concentration could be used as a switching mechanism. Indeed, mechanisms exist for the targeted degradation of ARFs (*44, 47*). However, two additional regulatory mechanisms would be possible based on our data. Firstly, we identify the accessibility of DNA as a key factor in DNA binding. Through multivalent interactions, nanoclusters can bind strongly to DNA if many binding sites are available. However, when shorter stretches of DNA are available for binding, this collectivity is lost causing clusters to engage DNA in a way that resembles monomeric binding. In a biological context, the length of accessible DNA may vary from spanning just a few base pairs to possibly kilobases (*60*). From an experimental point of view, this means that a DNA template of fixed, limited length likely provides a meaningful, but incomplete description of DNA binding. Biologically, this suggests that cells can tune the interactions between nanoclusters and DNA not only by changing the concentration of TFs, but also by regulating the accessibility of DNA (*68*).

Secondly, DNA-binding may be regulated through PB1 interactions. Our results indicate that the PB1 domain enhances nanocluster formation by promoting specific interactions between ARFs. Thus, we speculate that PB1 domains on other proteins may, through competition, suppress the formation of ARF nanoclusters and their subsequent DNA binding. Notably, the key binding partner of ARFs in the nuclear auxin pathway, the auxin-regulated Aux/IAA, contains a PB1 domain. In the absence of auxin, Aux/IAA binds the ARF through the PB1 interaction, preventing it from binding to DNA. When auxin is introduced, the Aux/IAA protein is degraded, which may enable ARF to form nanoclusters and perhaps bind its targets through a different mode. Based on our results, this picture can be interpreted as follows: without auxin, the PB1 interaction of Aux/IAA competes with PB1 interactions between ARFs, suppressing nanocluster formation. In the presence of auxin, this PB1 competition between Aux/IAA and ARFs is suppressed, releasing ARFs for mutual PB1 association which then switches on nanoclustering and thus the activation of DNA binding. While this remains speculative at this stage, if true, it could provide a mechanistic understanding of the regulatory role of TF nanoclusters as transcriptional switches.

## Materials and Methods

### Confocal microscopy *in vivo*

The *Marchantia polymorpha* MpARF2-mNeonGreen knock-in line used for *in vivo* imaging of MpARF2 was previously described in (*47*). Plants were grown on ½ strength Gamborg B5 medium at 22 °C with 40 μmol photons m^−2^ s^−1^ of continuous white light. Dormant gemmae from six-week-old plants were mounted on liquid Gamborg B5 medium and imaged using a Leica Stellaris inverted confocal microscope. mNeonGreen fluorescence was detected in photon-counting mode with time-gated acquisition to suppress autofluorescence. A 100× objective was used to obtain high-resolution recordings of nuclei. To estimate native MpARF2 concentrations in gemmae, *z*-stacks were taken using a 20× objective. The mean fluorescence intensity of nuclei, corrected for background, was measured in the image plane where each nucleus appeared brightest and converted to absolute concentrations by calibration. Calibration was performed by measuring purified mNeonGreen solutions (0-16 nM)) in TE7.5 buffer (20 mM Tris, 1 mM EDTA, 150 mM NaCl, pH 7.5) under identical imaging conditions.

### Cloning expression plasmids of MpARF2 variants

To simplify cloning, we first generated pET_MBP-3C-MCS-TEV-mNG-His, a pET plasmid containing a cloning site (MCS), flanked by regions encoding a 3C-cleavable MBP (N-terminal), and a TEV-cleavable mNG and His-tag (C-terminal). This was achieved through HiFi assembly (NEB) of linearized pET28a(+) plasmid, MBP and mNG (Table S2, pET28, MBP, mNG) along with a MCS dsDNA oligo (Table S1; MCS). This plasmid was linearized using restriction digestion or PCR (Table S2, pET_MCS) and MpARF2 variants were inserted as described in Table S3. The sequence of MBP was amplified from pET_His-MBP-TEV-LIC (Addgene #29656) provided by Balwina Koopal, and MpARF2 and mNG coding sequences were amplified from pMON_35S::MpARF2-mNG provided by Jan Willem Borst.

### Protein purification

All media were supplemented with 50 μg/mL kanamycin and 10 μg/mL chloramphenicol. Unless mentioned otherwise, cells were grown at 37 °C, 180 rpm. All chromatography steps were performed at RT. Rosetta™ (DE3) *E. coli* (Novagen) harboring pET vectors expressing the MBP-3C-MpARF-TEV-mNG-His fusion proteins were first grown overnight in 50 mL Luria Broth (LB). This preculture was then diluted 1:100 into 3 L Terrific Broth (TB). The newly inoculated culture was then grown to OD 0.6-0.8, after which 0.3 mM IPTG was added, the temperature was lowered to 20 °C, shaking speed was lowered to 160 rpm, and growth was continued overnight (18-20 h). Cells were harvested by centrifugation at 5,000× g, weighed (typically 30 g), and resuspended in lysis buffer (20 mM Tris, 1 M NaCl, 10% glycerol, 0.1% NP-40, 2 mM MgCl_2_, 10 mg DNAse I and cOmplete Protease Inhibitor, pH 8) at a ratio of 3 mL buffer per 1 g cell pellet. Cells were lysed by sonicating the suspension in a Q500 sonicator (Qsonica) equipped with a ½” probe at 80% amplitude for 2.5 minutes (5 sec on/20 sec off). Cell Free Extract (CFE) was generated by centrifuging the lysate at 50,000× g for 1 hour at 4 °C, and collecting the supernatant by gently decanting.

To the CFE, 10 mL of Nuvia™ Ni-IMAC resin (Bio-Rad) was added, and the mixture was shaken at RT at 120 rpm for 1 h. The mixture was then transferred to a column and the resin was collected by pumping from the bottom using a peristaltic pump. After removing the CFE, the resin was washed with 10 column volumes (CVs) of Ni Wash Buffer (20 mM Tris, 1 M NaCl, 10% glycerol, 1 mM TCEP, 20 mM imidazole, 0.01% Tween-20, pH 8), after which the protein was eluted with 2.5 CVs of Ni Elution Buffer (Ni Wash Buffer + 480 mM imidazole). The eluted protein was then passed over a Superdex™ 200PG column with bed dimensions 2.6 cm × 100 cm, equilibrated in SEC buffer #1 (20 mM Tris pH 8, 1 M NaCl, 10% glycerol, 1 mM EDTA, 1 mM DTT, 0.01% Tween-20). Monomer fractions were pooled, 5 mM DTT was added and incubated for 30 minutes. The pooled fractions were then loaded onto a 5 mL MBPTrap™ (Cytiva) equilibrated with SEC buffer #1, after which the column was washed with 2 CVs of SEC buffer #1, and the protein was eluted with 2 CVs of Amylose elution buffer (SEC Buffer #1 + 20 mM Maltose). We collected fractions containing the highest concentrations as estimated by A_280_ (usually ∼3 mL), concentration was calculated by measuring A_506_, using the ε_506_ of mNG (116,000 M^-1^ cm^-1^). Protein purity was assessed through SDS-PAGE and Coomassie staining. Purified protein was snap-frozen using liquid nitrogen in 100 μL aliquots and stored at -80 °C until use for experiments.

For centrifugation assays, confocal microscopy and dynamic light scattering, a fourth column is run on the day of the experiment. Up to 500 μL of MBP-tagged protein was thawed, to which HRV 3C protease (Sigma-Aldrich) was added at a ratio of 1 μg 3C per 1 nmol protein. After incubating for 2 h at room temperature, the mixture was passed over a Superdex 200 Increase 10/300 GL (Cytiva) equilibrated with SEC buffer #2 (20 mM Tris, 500 mM NaCl, 1 mM EDTA, 1 mM DTT, 0.001% Tween-20, pH 7.5), and monomer fractions were collected (retention volume ∼10 mL). If the concentration of the protein was insufficient, monomer fractions were pooled and concentrated using Amicon Ultra-0.5 10k MWCO Centrifugal Filters (Millipore). Concentration was then calculated by measuring A_506_ on a NanoDrop One (Thermo Fisher) in triplicate. The protein was stored in a Protein Lo-Bind microcentrifuge tube at RT, for up to one hour before use. For DLS, SEC was run in 1 M instead of 500 mM NaCl.

### Centrifugation assay

40 μL assays were prepared in a Protein Lo-Bind 1.5 mL microcentrifuge tube using a Hamilton syringe. Proteins were pre-diluted in high-salt buffer (20 mM Tris, 500 mM NaCl, 1 mM EDTA, 1 mM DTT, 0.001% Tween-20, pH 7.5) and brought to desired concentrations by 4× dilution in no-salt buffer (20 mM Tris, 0.001% Tween-20, pH 7.5). Samples were incubated for 15 min at RT and centrifuged for 1 min at 21,000× g at RT. After centrifugation, 25 μL supernatant was collected without disturbing the pellet and diluted in 75 μL assay buffer (20 mM Tris, 0.25 mM EDTA, 125 mM NaCl, 0.25 mM DTT, 0.001% Tween-20, pH 7.5). The protein concentration of diluted supernatant was determined from fluorescence intensity using a Synergy H1 Plate reader (Bio-Tek), using MBP-tagged protein at known concentrations for calibration. *c*_sup_ of the original sample was then calculated by correcting for the 4× dilution.

### Confocal microscopy

Confocal microscopy samples were prepared as described for centrifugation assays, with a total volume of 20 μL. Immediately after preparation, 5 μL sample was transferred onto a microscopy slide fitted with a Secureseal 8-wells imaging spacer (Grace Bio-Labs). A coverslip was added, and the slide was inverted and incubated for 15 min to allow the condensates to settle. Coverslip surfaces were imaged on a Leica SP8 equipped with single molecule detection using a 63× water-immersion objective. mNG fluorescence was excited with a pulsed white light laser at 488 nm and emission was collected from 515-545 nm with hybrid detectors. Samples for coarsening were prepared with a total volume of 40 μL, and imaged in a μ-slide 18-well glass coverslip (Ibidi). Solutions were then imaged at 15 minutes and 24 hours after sample preparation. Between measurements, the coverslip was sealed with parafilm and incubated in the dark at RT.

### Dynamic Light Scattering

Monomeric protein obtained after SEC was diluted in high salt buffer (20 mM Tris, 1 mM EDTA, 1 M NaCl, pH 7.5, sterile filtered). Protein variants were studied at 1 µM to provide enough signal to fit broad distributions. 150 mM NaCl samples were prepared by pre-diluting the protein in high salt buffer such that it could be brought to desired concentrations by dilution in no-salt buffer (20 mM Tris, 1 mM EDTA, pH 7.5, sterile filtered). Reversibility of clustering was assessed by first inducing MpARF2 clustering at 100 nM in 150 mM NaCl, as described above, and then diluting the sample to 50 nM MpARF2 in 1 M NaCl. Measurements were performed using disposable microcuvettes (BRAND; ≥ 100 nM) or quartz cuvettes (Hellma; < 100 nM) on a Malvern Zetasizer Nano with an acquisition time of 15 min (3-5 cycles). All DLS data presented are shown as intensity-weighted size distributions.

### DNA engineering

3IR7kb12 and NSkb12 DNA were obtained by building plasmids, consisting of one pUC19-derived backbone edited to contain minimal TGTC (Table S1, UC19’), and three unique TGTC-free ‘blocks’ amplified from *A. thaliana* genomic DNA. All assemblies were done using HiFi. Blocks were made from pairs of ∼1500 bp TGTC-free fragments (Table S2, A1, A2, B1, B2, C1, C2) and assembled into block plasmids (pBlockA, pBlockB, pBlockC). UC19’ was purchased as a gBlock (Integrated DNA Technologies) and assembled into a backbone plasmid (Table S4, pUC19’) using genomic filler DNA (Table S2, filler) that lacks TGTC but harbours a SapI site capable of producing a GGG-overhang. The extended backbone (Table S2, pUC19’) and blocks were amplified and assembled into p3IR7kb12 (Table S2, A_IR7, B_IR7, C_IR7) and pASkb12 (Table S2, A_NS, B_NS, C_NS) plasmids. Sequences were verified by nanopore sequencing (PlasmidSaurus). Plasmids were linearized using SapI and biotinylated using DNA polymerase Klenow fragment. Standard protocols were used, except for the use of biotin-dCTP instead of dCTP in the Klenow buffer. Custom Python software was used to scan the *Arabidopsis* genome for regions to amplify. IR7b1500 and NSb1500 were PCR amplified from corresponding 12 kb plasmids (Table S2, b1500) and purified using gel electrophoresis and cleanup. IR7b19 and NSb19 (Table S1) were obtained by annealing Cy5-labeled oligos in a 95 °C heat bath which was gradually cooled overnight. λ-DNA (NEB) templates were biotinylated in batches of 0.2 pmol by annealing primers (Table S2; CosR-bio, 2 pmol; CosL, 20 pmol) at 65 °C followed by ligation (NEB, T4 Ligase) for 2 h at RT, as described in (*69*). Biotinylated λ-DNA was separated from unreacted oligos using centrifugal spin filters (Amicon Ultra-0.5 100K) according to the supplier’s protocol.

### Single-particle tracking

MpARF2 aliquots were thawed and centrifuged at 21,000× g for 30 min, then diluted to desired concentrations in tracking buffer (20 mM Tris, 1 mM EDTA, 150 mM NaCl, 10% w/w PEG1000, pH 7.5). Single-particle tracking was performed on a Nikon Ti-2E inverted microscope equipped with a Teledyne Kinetix sCMOS camera and a 100× silicon oil immersion objective. Recordings of 301 frames were acquired in duplicate at 50 fps. Tracking analysis was carried out in Python using the TrackPy package. Trajectories shorter than 10 frames were excluded from analysis. The number of clusters was quantified as the average number of particles detected per field of view (208 × 208 µm).

### Electrophoretic Mobility Shift Assay

Agarose gels (15×15 cm) were prepared in 0.5× TBE, using two 15-well combs: one at the top and the other in the middle of the gel. 2% agarose was used for 19 bp DNA and 0.5% for other DNA lengths. Frozen MBP-MpARF2-mNG-His aliquots (1 µM, 100 µL) were thawed and cleaved with 3C protease (1 µg/nmol ARF) for 2 h at RT to cleave the MBP tag. 13 binding reactions (40 µL) containing DNA (50 ng) and protein (10-1000 nM, logarithmic, for DBD or 7.5 -97.5 nM, linear, otherwise) were prepared in binding buffer (20 mM HEPES, 100 mM Tris, 2.5% glycerol, 150 mM NaCl, pH 7.5) along with negative controls for protein and DNA, and equilibrated for 30 min at RT. The protein concentration of the control equals that of the middle lane. The samples were loaded using 10× Loading dye and ran for 90 min at 3 V/cm in 0.5× TBE at 4 °C. 19 bp DNA templates contained a Cy5 fluorescent label. Gels using unlabelled 1500 bp DNA were post-stained using 50 mL SxO (50 nM in Milli-Q) for 30 min on a rotating platform. Gels were imaged with a Typhoon gel scanner (Cytiva), detecting protein (Cy2 filter set) and DNA (Cy3 or Cy5 filter set depending on the stain) fluorescence. Custom Python software was used to integrate band intensities and to perform subsequent analysis. Fluorescence signals coming from the wells, representing a clustered bound fraction, were included in the analysis; the fraction of bound DNA was calculated as

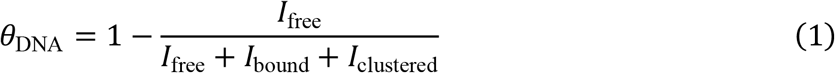

The fraction of bound protein was calculated similarly, instead using protein fluorescence. Band intensities were corrected for background fluorescence by subtracting signals coming from the appropriate negative controls.

### Pretreatment of coverslips for single molecule imaging

Microfluidic devices were assembled and surface-treated in batches of eight. A clean glass staining jar (BR472700, Sigma-Aldrich) was filled with 70 mL of spectroscopy-grade acetone. Wearing powder-free gloves, glass coverslips (24×60 mm #1.5) were rinsed with Milli-Q water and isopropanol to remove particulates and organic residues. Coverslips were then dried using compressed air and treated with plasma (Harrick PDC-32g) for 1 min. After plasma, the coverslips were immediately placed in acetone and transferred to a fume hood. Wearing gloves, 1.4 mL VECTABOND (VectorLabs) was added and mixed by pipetting up and down without introducing bubbles. After incubating for 15 min, VECTABOND solution was discarded appropriately. Coverslips were rinsed once with acetone and five times with Milli-Q. Slides were dried gently with compressed air and temporarily kept in falcon tubes.

### Microfluidic chip assembly

Microfluidic chips were assembled from a pretreated coverslip, a double-sided tape spacer, and a PDMS lid. Double-sided tape (Hi-Bond, HB397F-25), protected by foil on both sides, was laser-cut into 25×20 mm rectangular spacers containing a 10×1 mm channel. To obtain protective backing on both sides of the tape, two strips were stacked front-to-back, and the exposed layer of tape was peeled off. Rectangular blocks of 25×20×5 mm were cast from PDMS (10:1 PDMS:resin) and provided with an inlet and outlet using a biopsy punch. Punctured PDMS pieces were rinsed with water and ethanol, air dried, and treated with plasma for 15 seconds. Chips were assembled by attaching tape to the cover glass and sticking the PDMS lid on top. Chips were firmly pressed to ensure a tight seal.

### Passivation of microfluidic chips

4 mg Biotin-PEG-SC (MW 5000, Laysan Bio) and 32 mg mPEG-SVA (MW 5000, Laysan Bio) were dissolved in 160 µL MOPS buffer (50 mM, pH 7.5). The solution was vortexed for 1 min and centrifuged for 1 min at 21,000× g. 20 µL of the PEG solution was pipetted directly into the chips, and the inlet and outlet were covered with parafilm to prevent air from entering. The chips were incubated for 3 h at RT in a humidity box, which had been prepared by partially filling an empty pipette tip box with water. A secondary PEGylation was performed to further improve passivation. A solution of 25 mM MS4-PEG (Thermo Fisher) in sodium bicarbonate buffer (0.1 M, pH 8,5) was prepared and stored at -20 °C in 500 µL aliquots. After thawing, an aliquot was centrifuged for 1 min at 21,000× g before injecting 50 µL into each chip. The chips were incubated overnight at 4 °C in a humidity box.

### Protein labelling for single molecule imaging

Frozen MBP-MpARF2-mNG-His aliquots (100 µL, 1 µM) were thawed and brought to an amine-free labelling buffer (20 parts PBS, adjusted to 1 M NaCl; 1 part 0.2 M sodium bicarbonate, pH 9) using a desalting spin column (Zeba Spin, Cat. 89882) according to the supplier’s protocol. Protein concentration was measured using a Nanodrop, and a 10× molar excess of ATTO633-NHS was introduced and incubated protected from light for 1 hour at RT. Degree-of-labelling was measured according to the ATTO-NHS labelling protocol, which involves separating free and unreacted dye via a gravity-flow desalting column (BIO-RAD; 10DG). Aliquots of 10 pmol labelled protein were flash-frozen in 1.5 mL LoBind tubes (Eppendorf) and stored at -80 °C.

### Single-molecule imaging

Microfluidic chips were mounted on a custom built TIRF microscope (*70*) and connected to a distributor valve (MUX-D-12, Elveflow) using Tygon tubing. Three distributor valve positions were used to connect (i) a 1.5 mL tube for samples, (ii) a 15 mL tube for buffers, and (iii) a plug to interrupt flow. DNA tethering was achieved by sequentially flushing in various 1 mL solutions with incubating steps in between. Firstly, chips were washed with TE7.5 buffer (10 mM Tris, 1 mM EDTA, 150 mM NaCl, pH 7.5) to provide a clean starting point. Next, NeutrAvidin (0.05 g/L in TE7.5) was incubated for 5 min and flushed out using blocking buffer (1 g/L Pluronic F108, 1 g/L BSA in TE7.5 buffer). Finally, biotinylated DNA (1 pM in blocking buffer) was introduced and incubated for 1 h. Tethered DNA was visualized and excess DNA was removed by flushing with imaging buffer (50 nM SxO in blocking buffer). Frozen ATTO633-labelled protein aliquots were thawed, cleaved with 3C protease (2 h, RT, 1 µg/nmol sample), and centrifuged for 1 min at 21,000× g to remove large aggregates. Samples were adjusted to contain 50 nM SxO and 150 mM NaCl. Recordings (Andor Zyla 5.5 sCMOS) of alternating excitation (561 nm, 642 nm, 50 ms interval) were started after which flow was switched on to flush in protein using gravity at an approximate flow rate of 100 µL/min for 5-10 minutes.

### Single-molecule imaging – data analysis

Fields of view (FOVs) were manually marked to indicate regions of interest (ROIs) and flow direction. A convolutional median filter (101×101 pixels) was applied to correct for uneven illumination before extracting ROI data as individual files for further analysis. For each ROI, DNA was detected by applying a 3D Gaussian blur followed by binarization and a series of morphological operations (erosion, dilation, skeletonization, pruning) that yield an unbranched chain one pixel in width. This chain is expanded to create a DNA mask, which was applied to binarized ROIs (for coverage analysis) and to blurred ROIs (for specificity analysis). Filtered ROIs were inspected and excluded in case DNA detection failed. DNA coverage, *ϕ*, was quantified from binary ROI data as

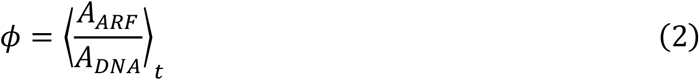

where *A* represents the area of the signal in pixels. Sequence specificity was quantified as the enrichment of protein on the DNA compared to the environment (⟨*I*_in_⟩ /⟨*I*_out_⟩), averaged over time and across ROIs. The DNA intensity profile along the *x*-axis was obtained by averaging the protein signal over the *y*-axis, time, and across ROIs.

## Acknowledgments

We dedicate this paper to the memory of our colleague, mentor and good friend Hanne van der Kooij, whose contributions to our team were immense. We gratefully acknowledge Dylan Jansen for technical contributions at the initiation of this work.

## Funding

European Research Council Consolidator grant Catch 101000981 (JS).

European Research Council Advanced grant DIRNDL 833867 (DW).

Human Frontiers Science Program grant RGP0015/2022 (DW).

Marie Skłodowska-Curie Individual Fellowship 2020 grant REOX 101026004 (JHG).

Leica Stellaris STED microscope is funded by NWO roadmap grant 184.036.012.

## Author contributions

Conceptualization: KA, BJ, JWB, DW, CvM, JS.

Cloning of MpARF2 constructs: BJ, JWB, CvM.

Purification of MpARF2 and variants: BJ, JWB, CvM.

Preparing DNA constructs: KA, JHG, JJR.

*In vivo* experiments: KA, MDA, JWB.

*In vitro* confocal microscopy: BJ, CvM.

Centrifugation assay: BJ, CvM.

Electrophoretic mobility shift assay: KA, RR.

Dynamic Light Scattering: KA, MvdW.

Single-Molecule Imaging: KA, LJ, SvdB.

Single-Particle Tracking: KA, MR.

Data Analysis Conceptualization: KA, JS.

Programmatic Data Analysis: KA.

Writing: KA, JS. All authors reviewed the manuscript.

## Competing interests

Authors declare that they have no competing interests

## Data and materials availability

All data analysis codes used in the work presented in this article have been made publicly available at https://git.wur.nl/bic/2025_mparf2-nanoclusters. All data will be made publicly available upon request.

## Notes

### Competing Interest Statement

The authors have declared no competing interest.

