## Supplementary information for "Nanoclustering of a plant transcription factor enables strong yet specific DNA binding"

Science Advances

AAAS

Kasper Arfman *et al.*

**This PDF file includes:**

Fig. S1. Structure and domains of MpARF2
Fig. S2. FuzDrop prediction of MpARF2
Fig. S3. MpARF2 nanoclusters/condensates before and after centrifugation
Fig. S4. Growth kinetics of MpARF2 assemblies
Fig. S5. Centrifugation assay of the ΔMR protein variant
Fig. S6. Centrifugation assay of the MpARF2-MR protein variant
Fig. S7. Confocal image of MpARF2-MR nanoclusters
Fig. S8. AlphaFold-Multimer prediction of MpARF2 pentamers
Fig. S9. MpARF2 nanocluster abundance as a function of protein concentration
Fig. S10. EMSA gel of MpARF2-DBD and 19 bp DNA
Fig. S11. EMSA gel of MpARF2 and 19 bp DNA
Fig. S12. EMSA gel of MpARF2-DBD and 1500 bp DNA
Fig. S13. EMSA gel of MpARF2 and 1500 bp DNA
Fig. S14. Binding kinetics in single-molecule imaging
Fig. S15. EMSA of ΔPB1 on IR7b1500 DNA
Fig. S16. Rare example of an MpARF2 nanocluster binding to nonspecific DNA
Table S1. dsDNA sequences
Table S2. ssDNA sequences
Table S3. Strategies used to generate MpRF2 variant plasmids
Table S4. Overview of plasmids used for generating protein variants and DNA templates

**Other Supplementary Materials for this manuscript include the following:**

Movie S1 (.mp4 format). MpARF2 nanoclusters inside the nucleus at native expression
Movie S2 (.mp4 format). Coalescence of MpARF2 condensates
Movie S3 (.mp4 format). Single-molecule assay of a DNA-bound nanocluster

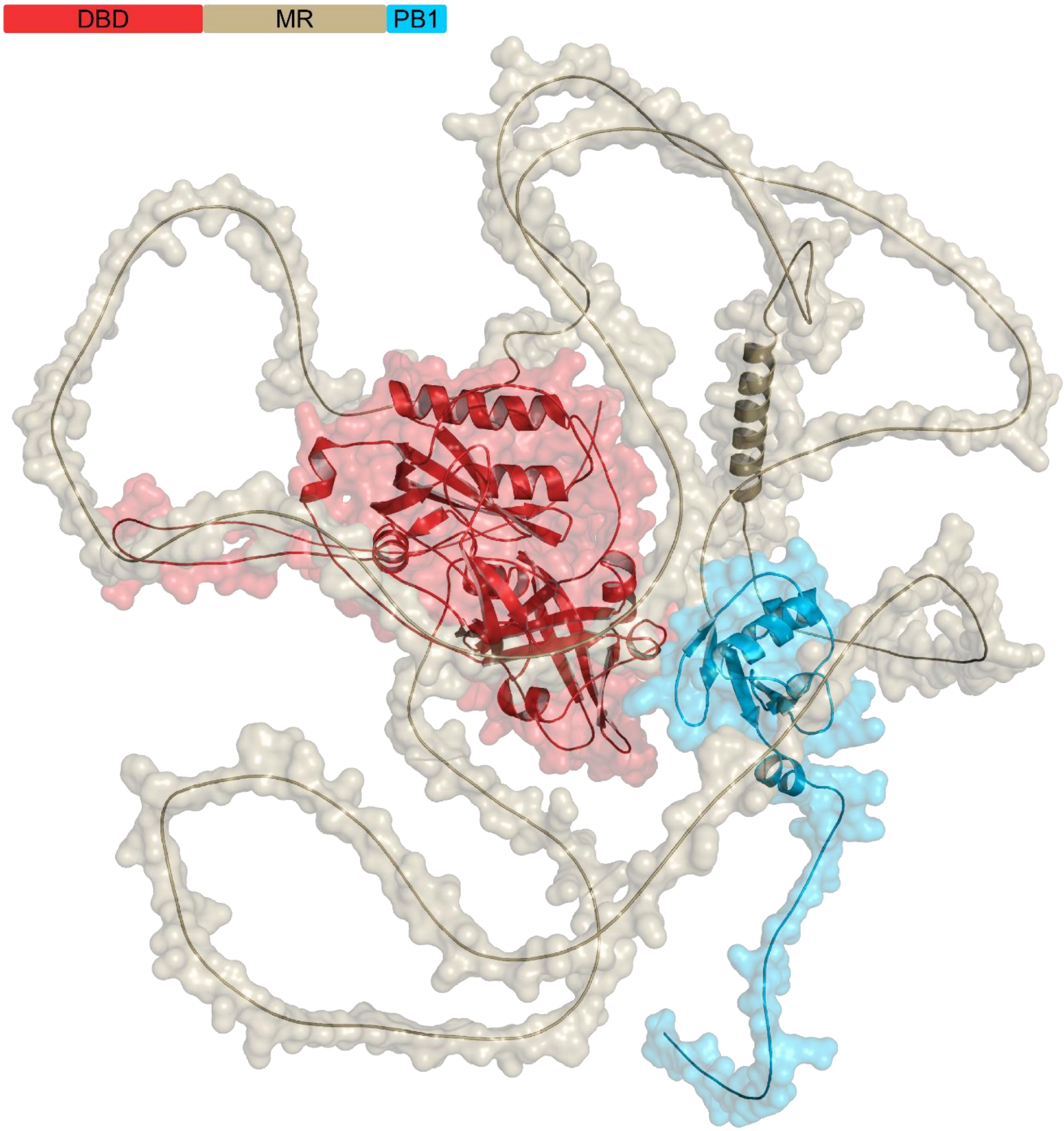


**Fig. S1. Structure and domains of MpARF2**

AlphaFold structure prediction of MpARF2, consisting of a DNA-binding domain (DBD, red), an intrinsically disordered middle region (MR, sand), and a Phox and Bem1 multimerization domain (PB1, cyan).

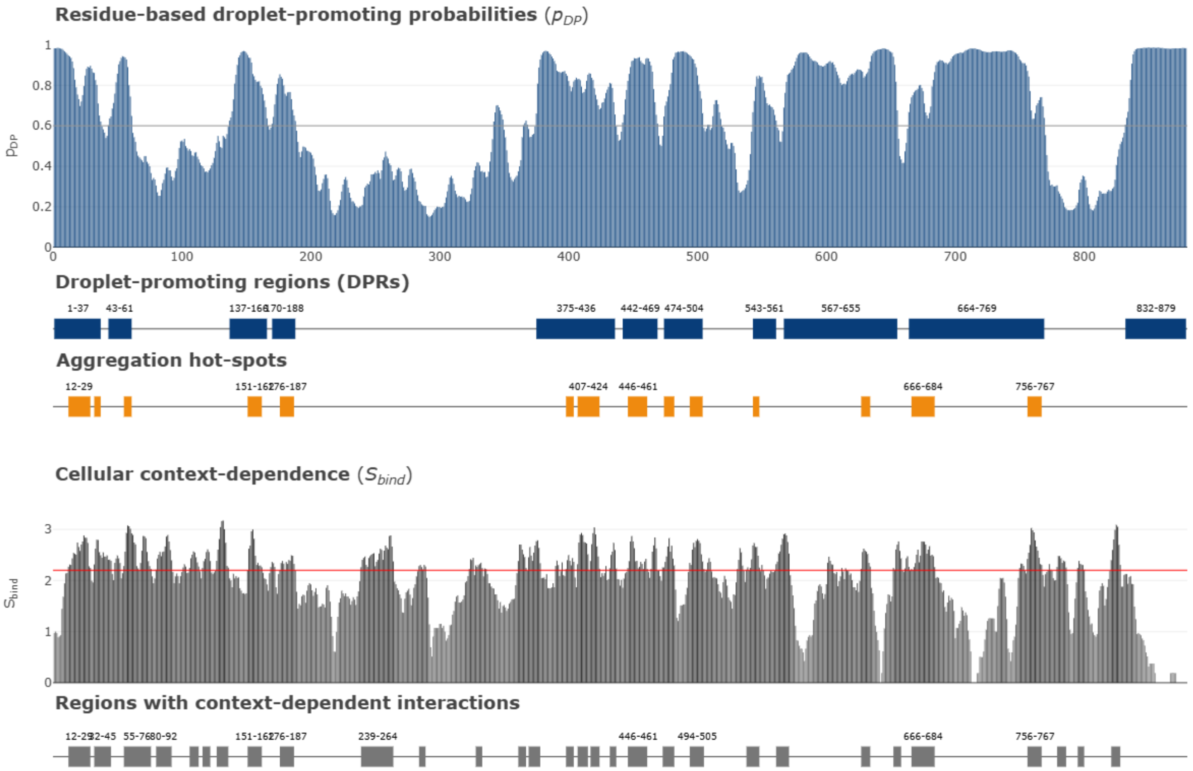


**Fig. S2. FuzDrop prediction of MpARF2**

MpARF2 consists of a DBD (1-394), MR (395-744) and PB1 (745-878) domain. The probability of liquid-liquid phase separation, $P_{\mathrm{LLPS}}$, is 0.9975. The residue-based droplet-promoting probability, $p_{\mathrm{DP}}$, quantifies how likely an amino acid is to contribute to liquid-liquid phase separation. Stretches where $p_{\mathrm{DP}} > 0.6$ are marked as droplet-promoting regions. Aggregation hot-spots are sequence elements that tend to self-associate into aggregates, rather than remaining in a dynamic, liquid-like state. The dependence on cellular context indicates the extent to which the tendency for droplet formation depends on the molecular environment.

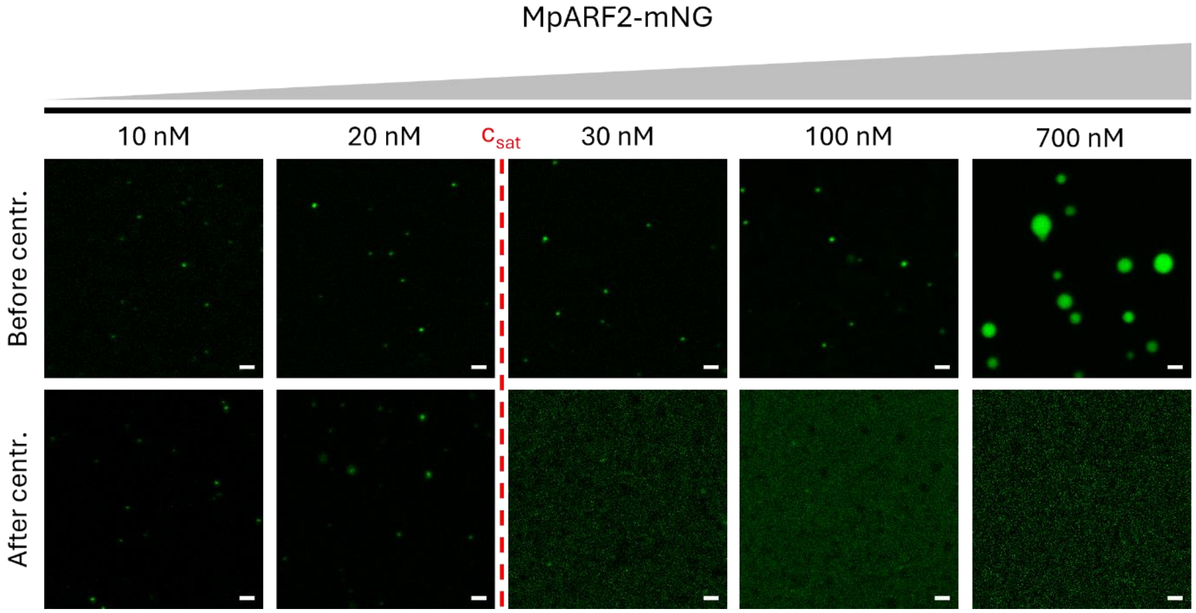


**Fig. S3. MpARF2 nanoclusters/condensates before and after centrifugation**

Confocal fluorescence microscopy images of MpARF2-mNeonGreen solutions at various concentrations before and after centrifugation. Post-centrifugation images above $c_{sat}$ are shown with enhanced intensity to emphasize the absence of protein assemblies. Scale bar: 1 µm.

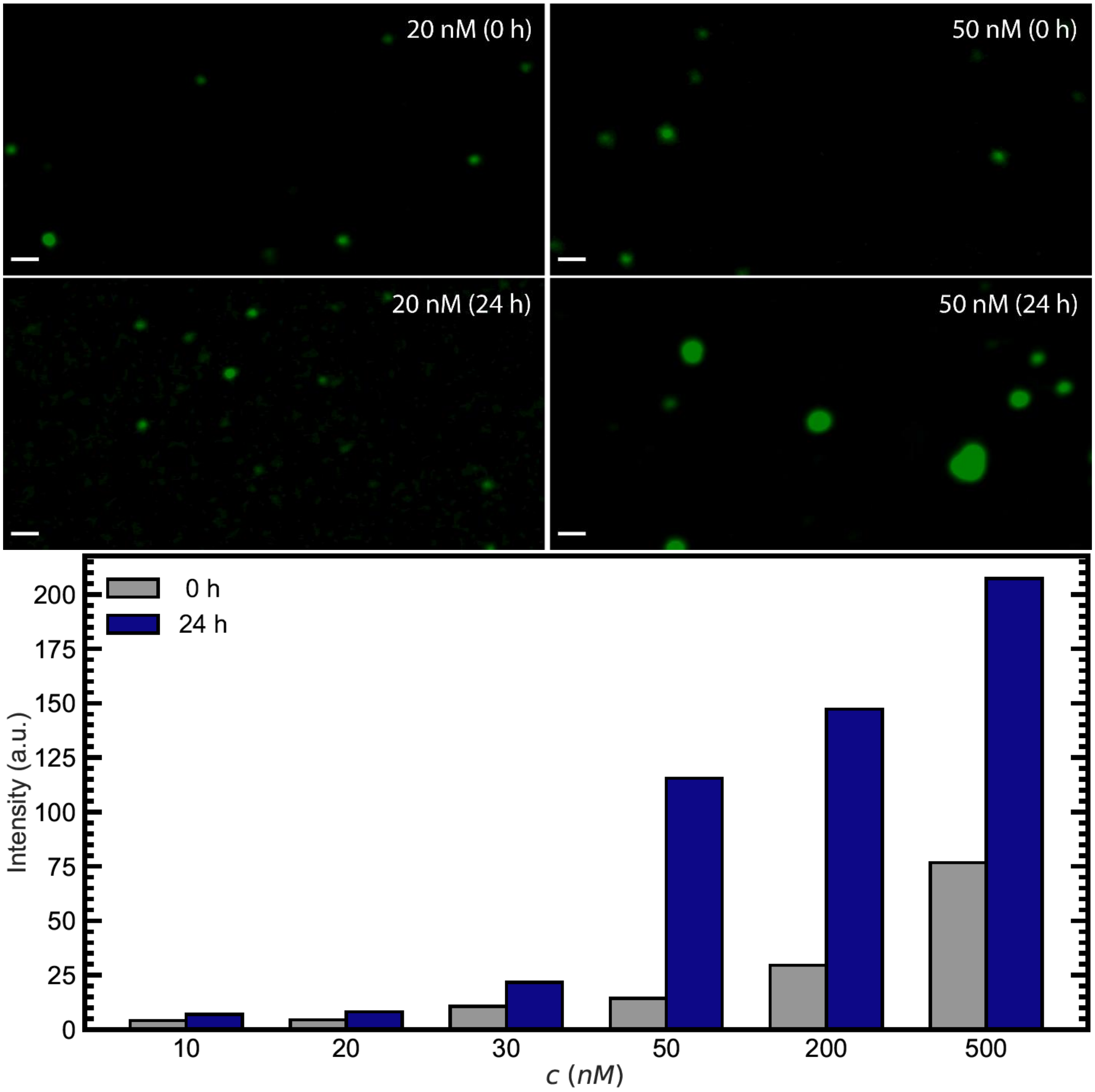


**Fig. S4. Growth kinetics of MpARF2 assemblies**

**Top** Confocal microscopy images of purified MpARF2-mNG after inducing phase separation by dilution to physiological salt concentration. The protein concentrations are just above (50 nM) and below (20 nM) $c_{\text{sat}}$. The transition of diffraction-limited assemblies to micron-scale assemblies indicates coarsening as predicted by phase separation, confirming $c_{\text{sat}}$ and distinct identity of nanoscale assemblies below $c_{\text{sat}}$. Scale bar: 1 µm. **Bottom** Degree of clustering quantified as the average brightness of pixels exceeding the background level intensity by five standard deviations (*y*-axis). Subsaturated solutions (< 28 nM) show a limited brightness increase over 24 hours, whereas solutions containing condensates (> 28 nM) exhibit substantial growth.

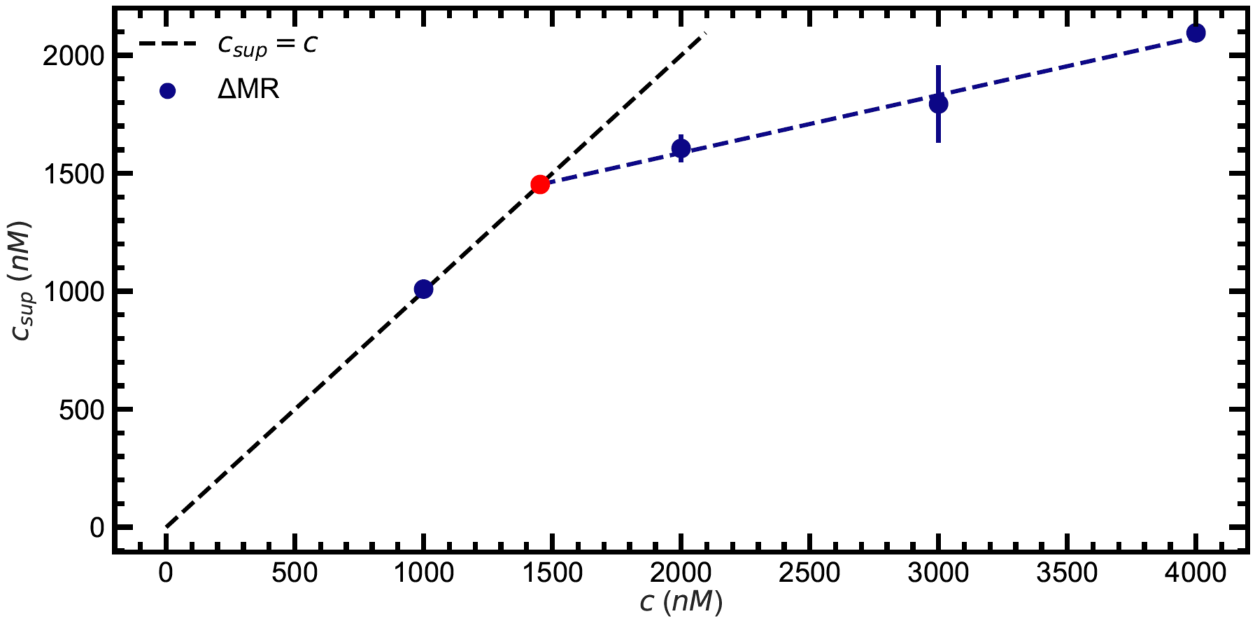


**Fig. S5. Centrifugation assay of the ΔMR protein variant**

The ΔMR concentration in the supernatant, $c_{\mathrm{sup}}$, is measured after centrifugal removal of condensates. The red dot indicates that $c_{\mathrm{sat}}$ is ~1.5 µM.

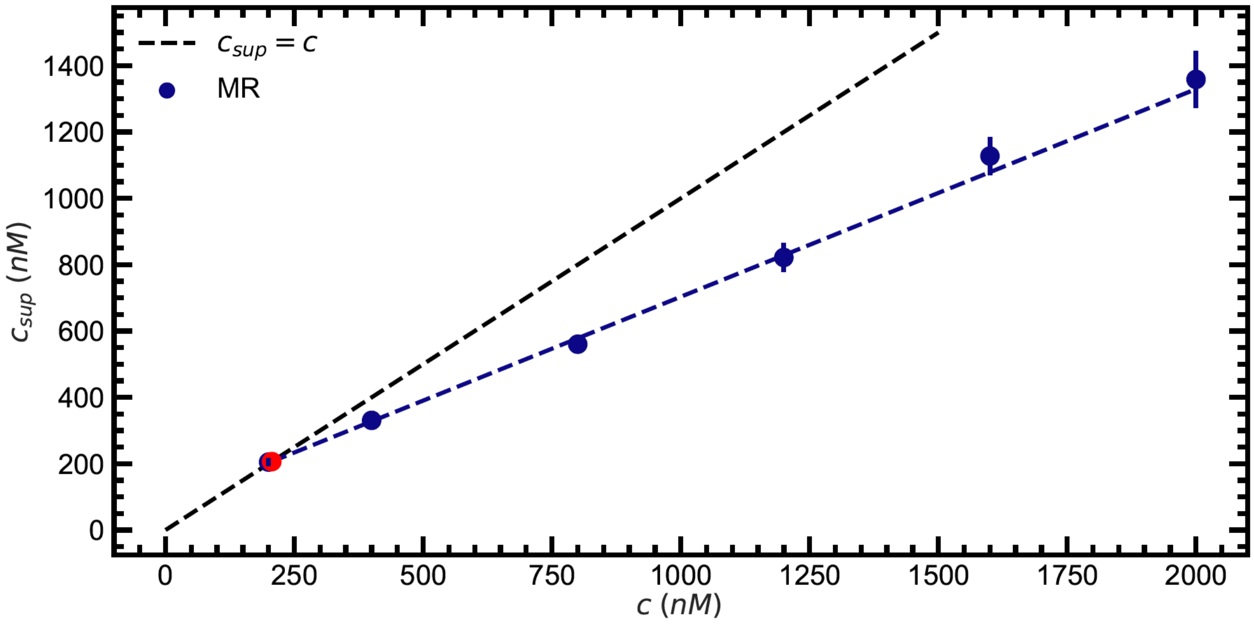


**Fig. S6. Centrifugation assay of the MpARF2-MR protein variant**

The MpARF2-MR concentration in the supernatant, $c_{\text{sup}}$, is measured after centrifugal removal of condensates. The red dot indicates that $c_{\text{sat}}$ is ~200 nM.

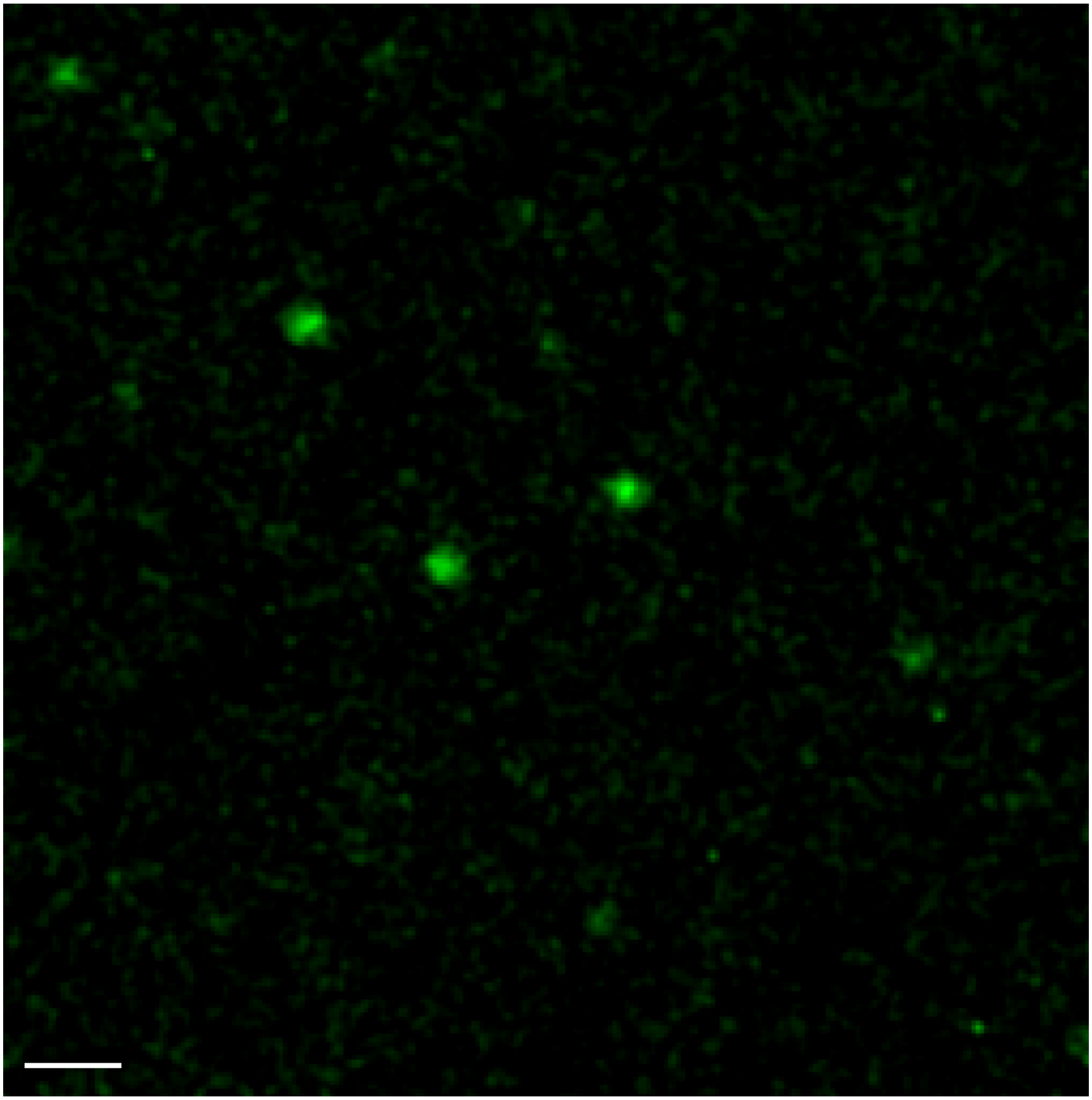

**Fig. S7. Confocal image of MpARF2-MR nanoclusters**

Section of a confocal image of MpARF2-MR in assay buffer (20 mM Tris pH 7.5, 125 mM NaCl, 0.25 mM DTT, 0.25 mM EDTA, 0.001% Tween-20), showing the presence of nanoclusters at a protein concentration of 15 nM, well below the threshold for condensate formation ($c_{\mathrm{sat}} \approx 200$ nM). Scale bar: 1 µm.

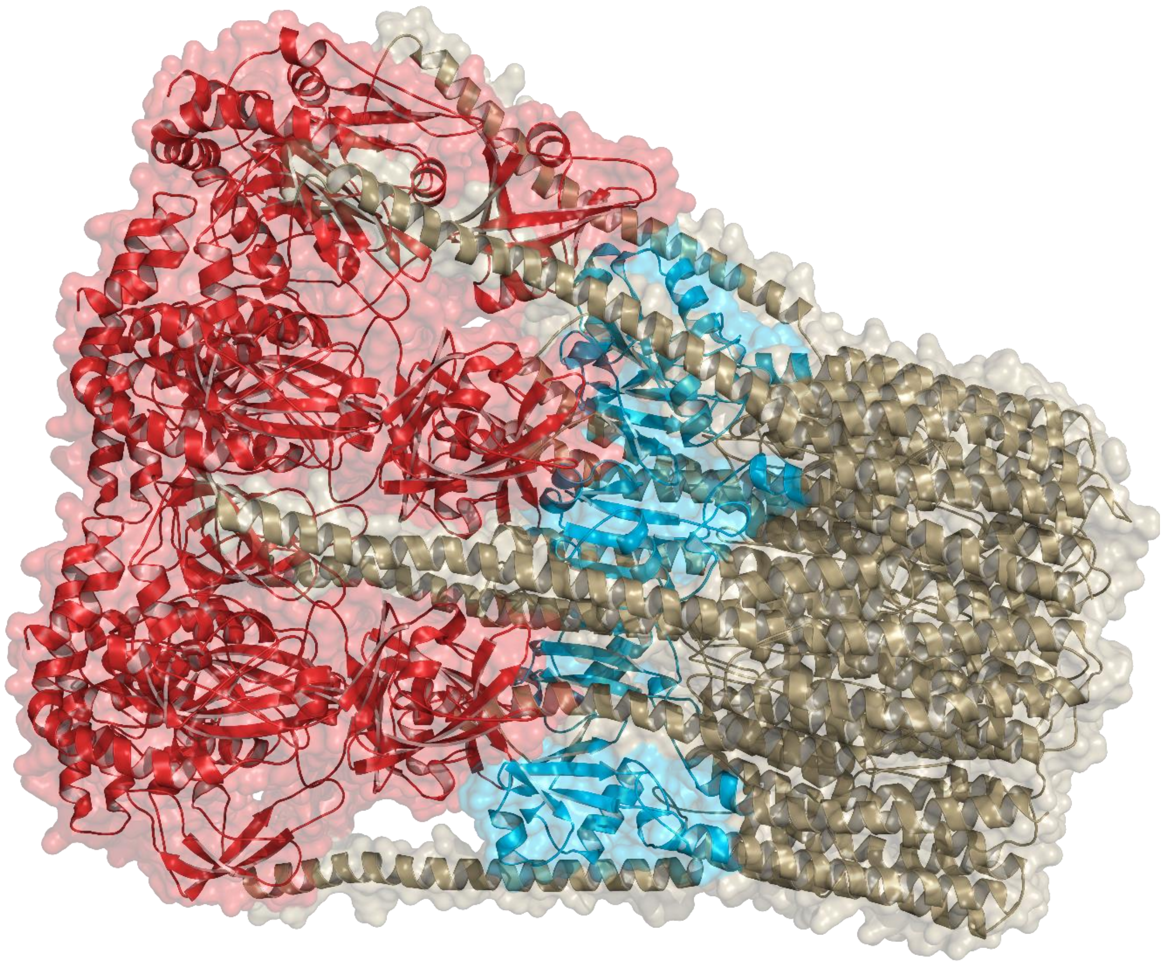

**Fig. S8. AlphaFold-Multimer prediction of MpARF2 pentamers**
Colors refer to the DBD (red), the MR (sand) and the PB1 (cyan) domains.

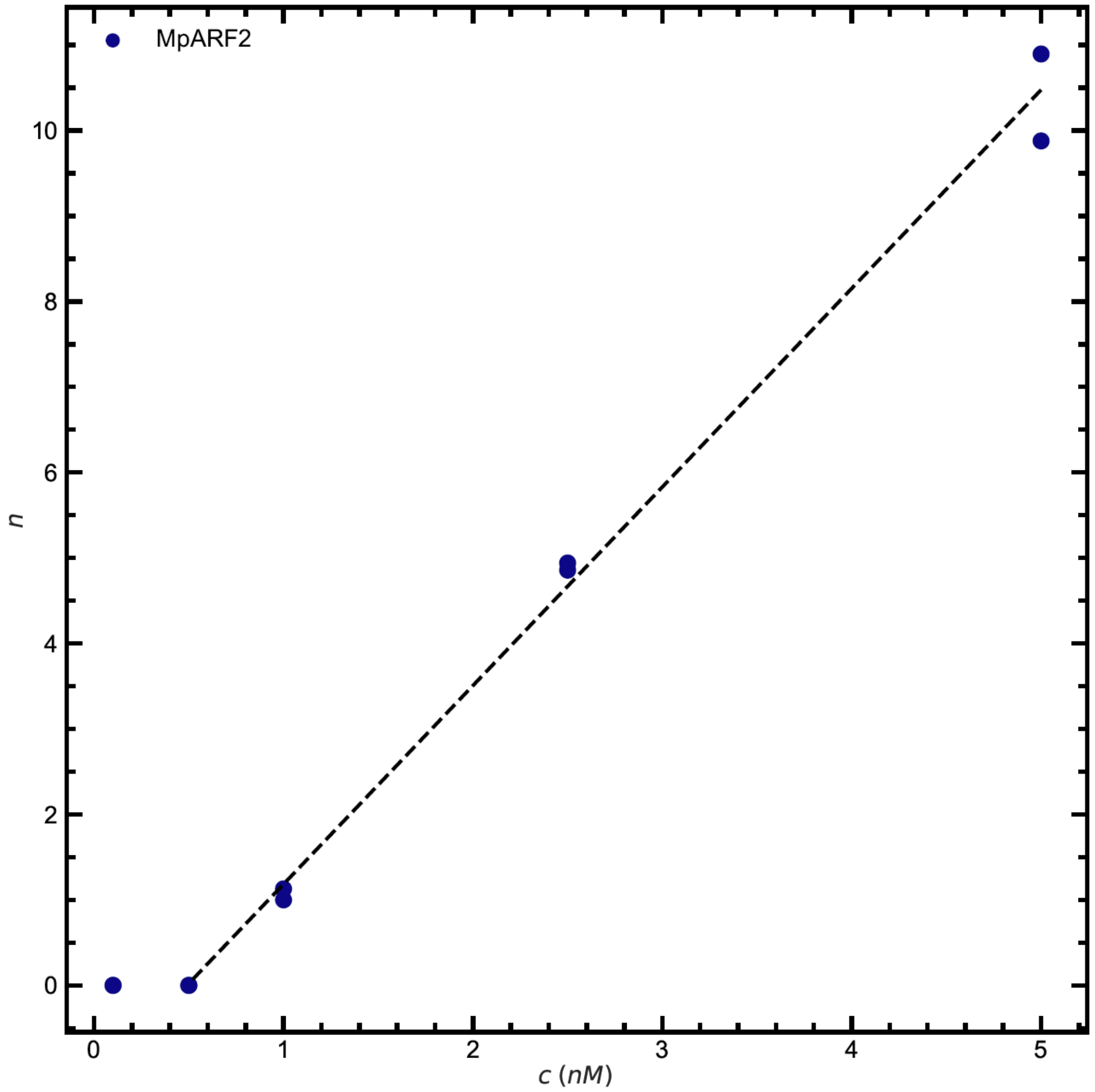


**Fig. S9. MpARF2 nanocluster abundance as a function of protein concentration**

Average number of MpARF2 nanoclusters per field of view (208 × 208 µm) plotted as a function of protein concentration. A sharp increase in cluster number is observed above a critical concentration of ~0.5 nM, followed by a linear increase with concentration. Nanocluster counts were obtained by single-particle tracking analysis (301 frames, 20 ms intervals) performed in duplicate. Trajectories shorter than 10 frames were excluded to minimize false-positive detections.

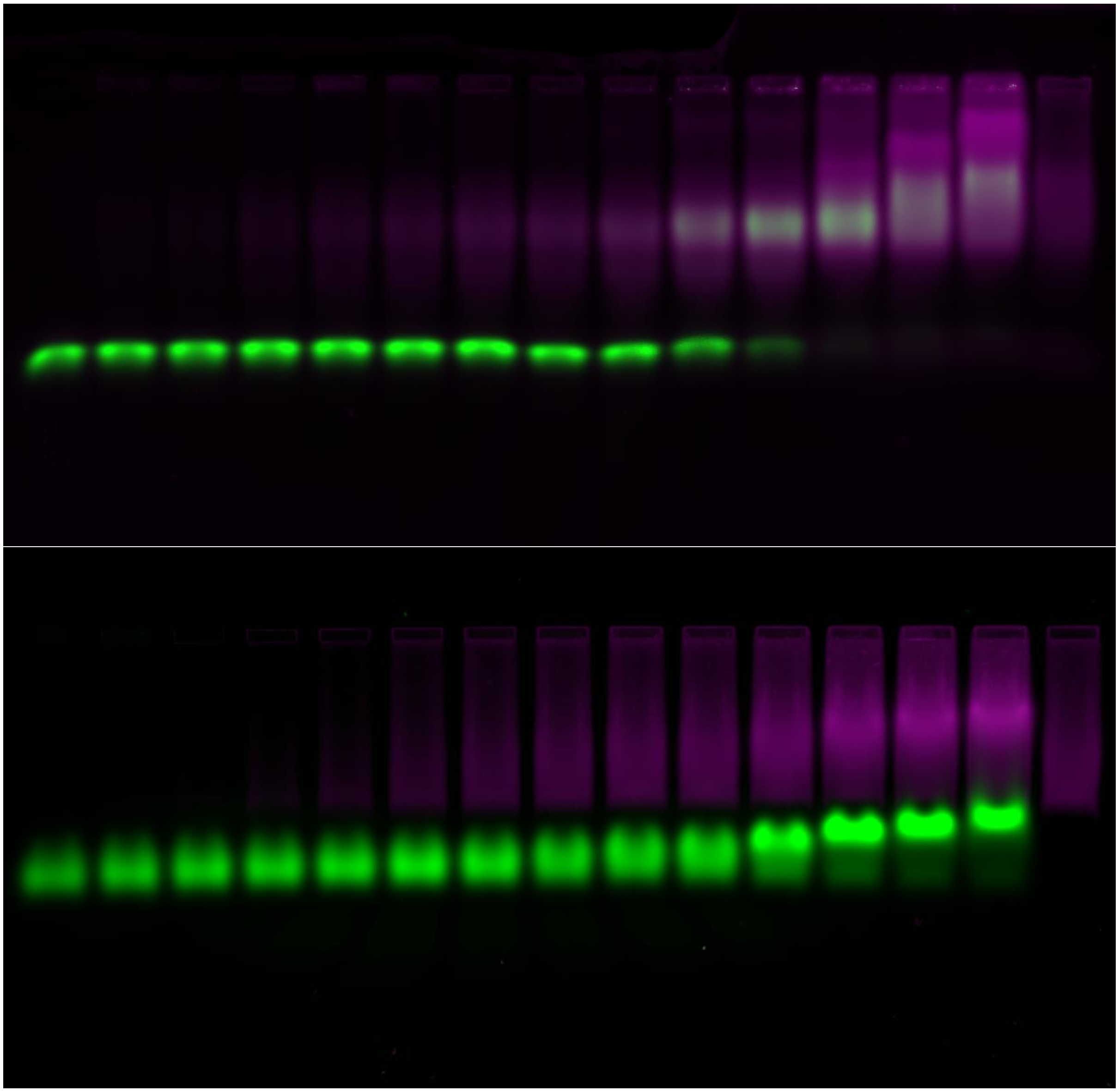

**Fig. S10. EMSA gel of MpARF2-DBD and 19 bp DNA**

EMSA gel of MpARF2-DBD (magenta, log color scale) with 19 bp DNA (green). **Top** Specific DNA (IR7b19). **Bottom** Nonspecific DNA (NSb19). Concentration range: 10-1000 nM logarithmically spaced. First and last wells are controls containing only DNA and protein, respectively. Binding conditions: 20 mM HEPES, 100 mM Tris, 2.5% glycerol, 150 mM NaCl, pH 7.5.

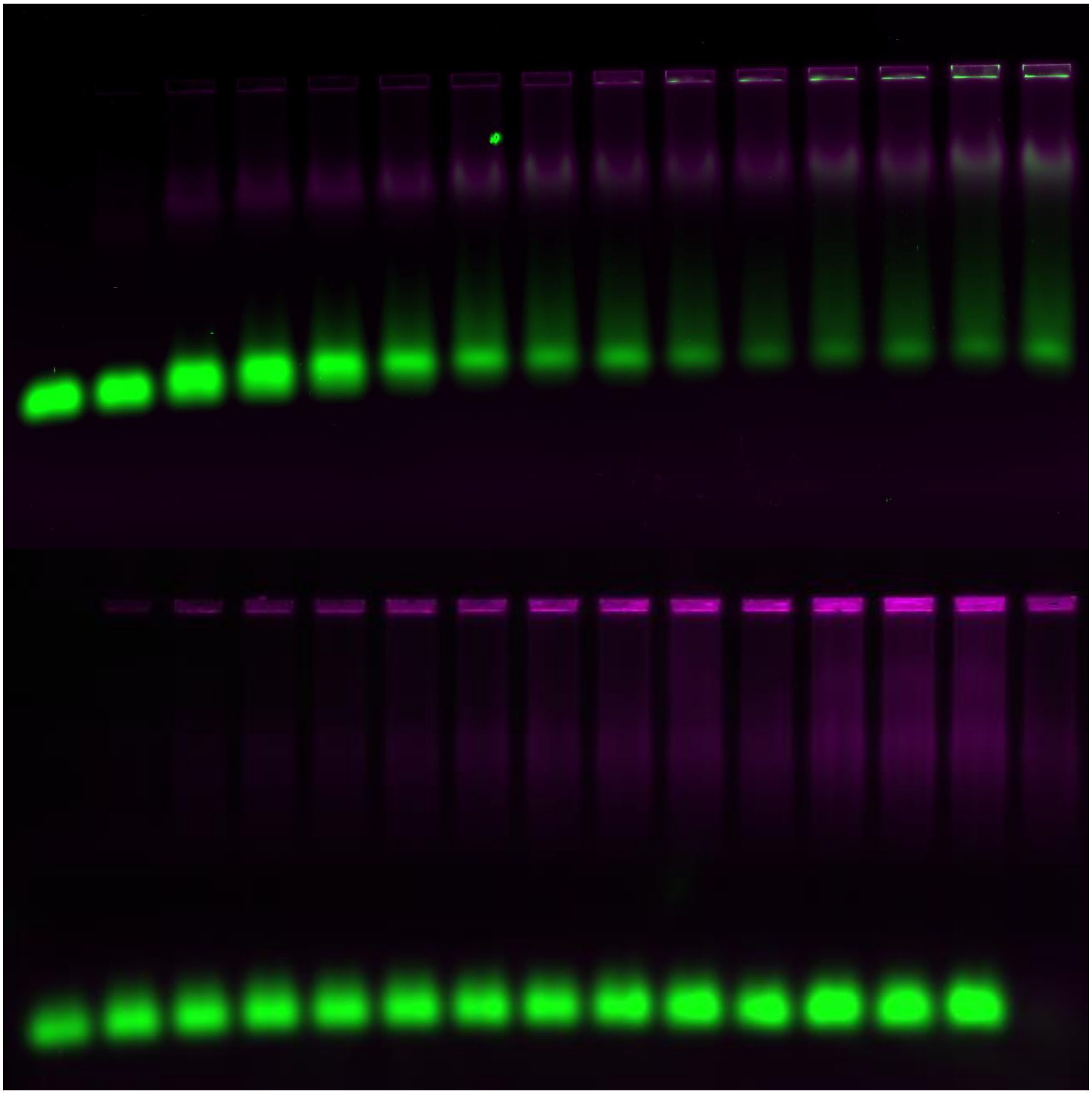

**Fig. S11. EMSA gel of MpARF2 and 19 bp DNA**

EMSA gel of MpARF2 (magenta, log color scale) with 19 bp DNA (green). **Top** Specific DNA (IR7b19). **Bottom** Nonspecific DNA (NSb19). Concentration range: 7.5 – 97.5 nM linearly spaced. First and last wells are controls containing only DNA and protein, respectively. An exception is the last well of the top gel, which erroneously contains DNA and was excluded from analysis. Binding conditions: 20 mM HEPES, 100 mM Tris, 2.5% glycerol, 150 mM NaCl, pH 7.5.

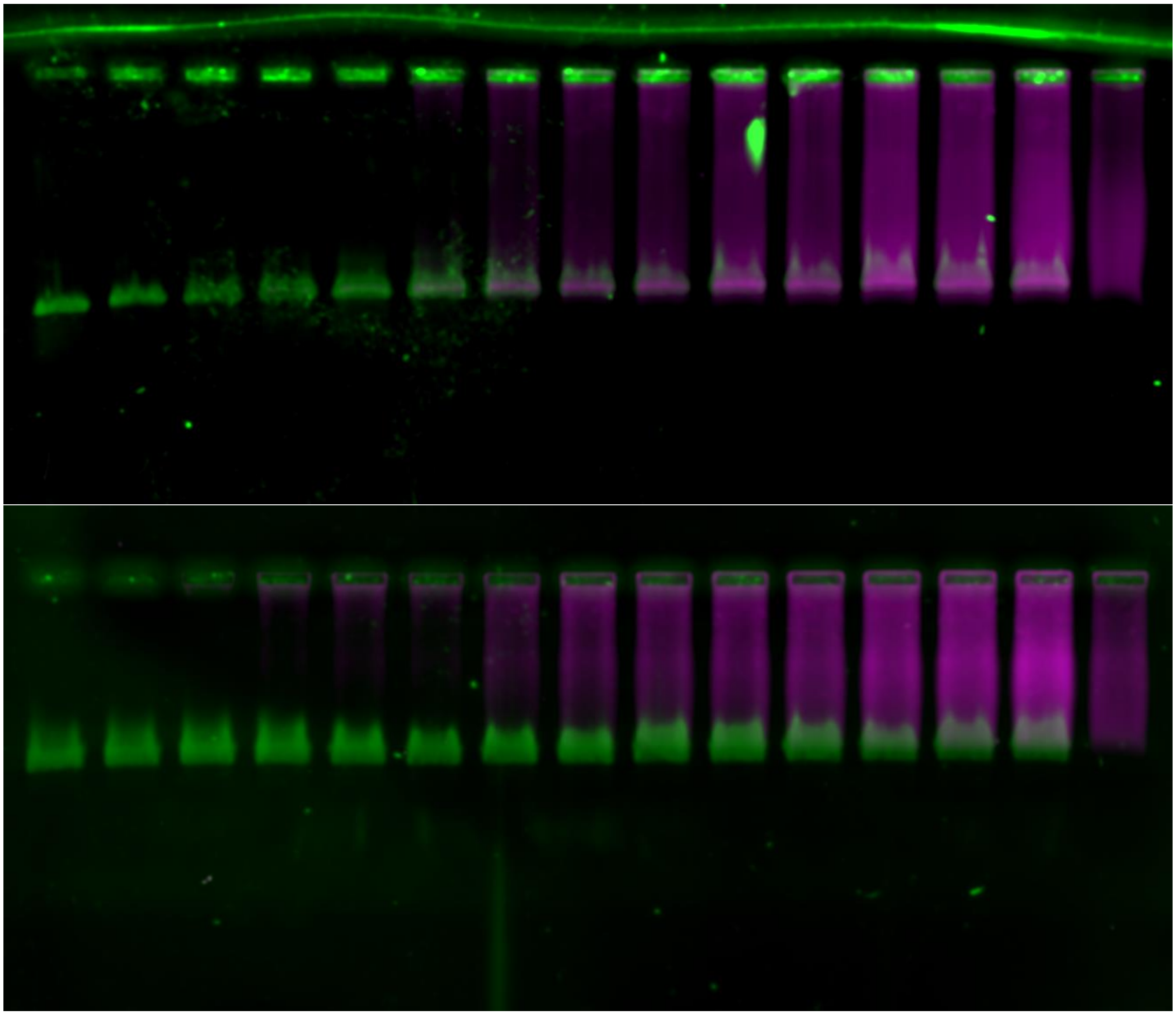

**Fig. S12. EMSA gel of MpARF2-DBD and 1500 bp DNA**

EMSA gel of MpARF2-DBD (magenta, log color scale) with 1500 bp DNA (green). **Top** Specific DNA (IR7b1500). **Bottom** Nonspecific DNA (NSb1500). Concentration range: 10-1000 nM logarithmically spaced. First and last wells are controls containing only DNA and protein, respectively. Binding conditions: 20 mM HEPES, 100 mM Tris, 2.5% glycerol, 150 mM NaCl, pH 7.5.

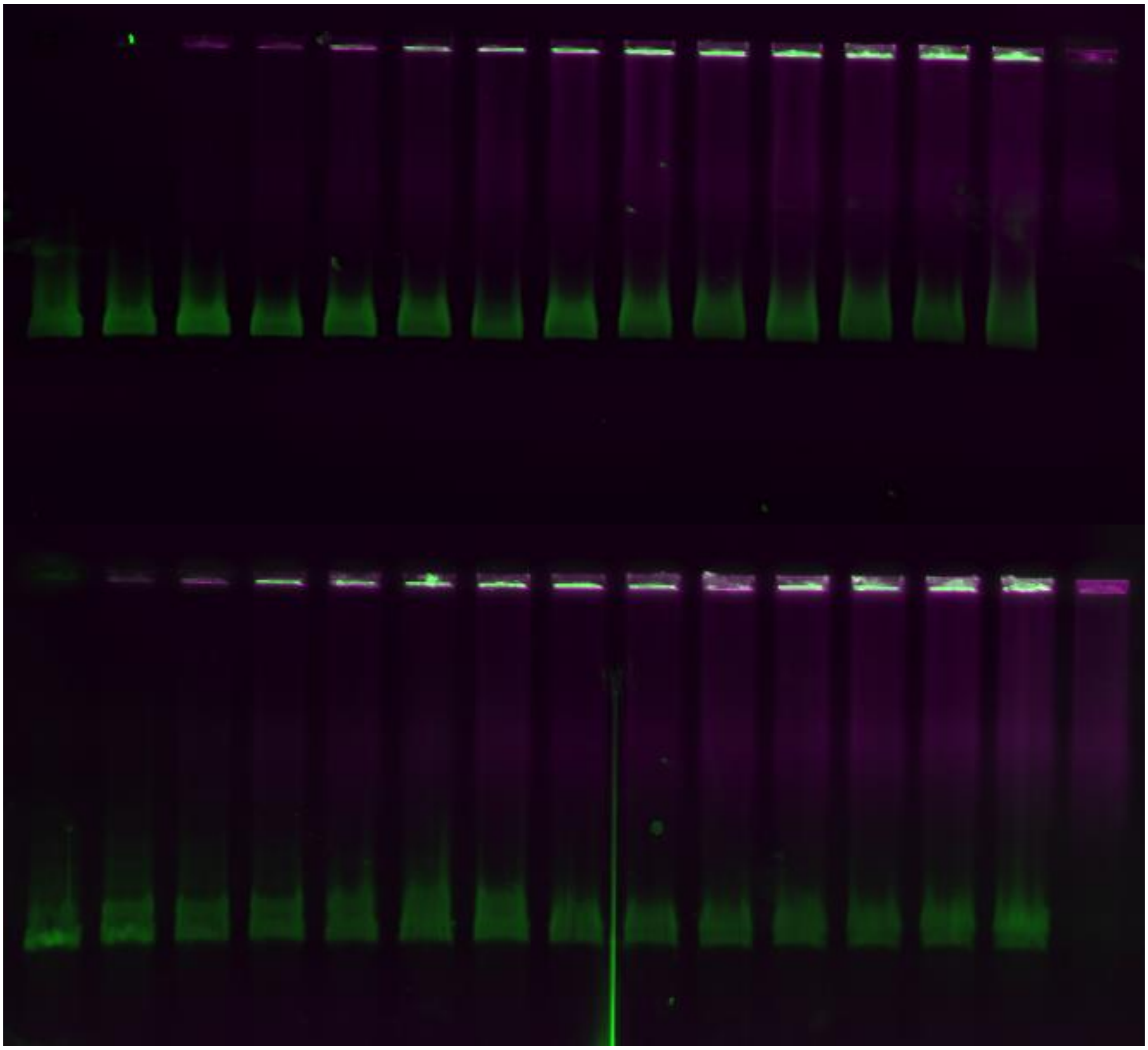

**Fig. S13. EMSA gel of MpARF2 and 1500 bp DNA**

EMSA gel of MpARF2 (magenta, log color scale) with 1500 bp DNA (green). **Top** Specific DNA (IR7b1500). **Bottom** Nonspecific DNA (NSb1500). Concentration range: 7.5 – 97.5 nM linearly spaced. First and last wells are controls containing only DNA and protein, respectively. Binding conditions: 20 mM HEPES, 100 mM Tris, 2.5% glycerol, 150 mM NaCl, pH 7.5.

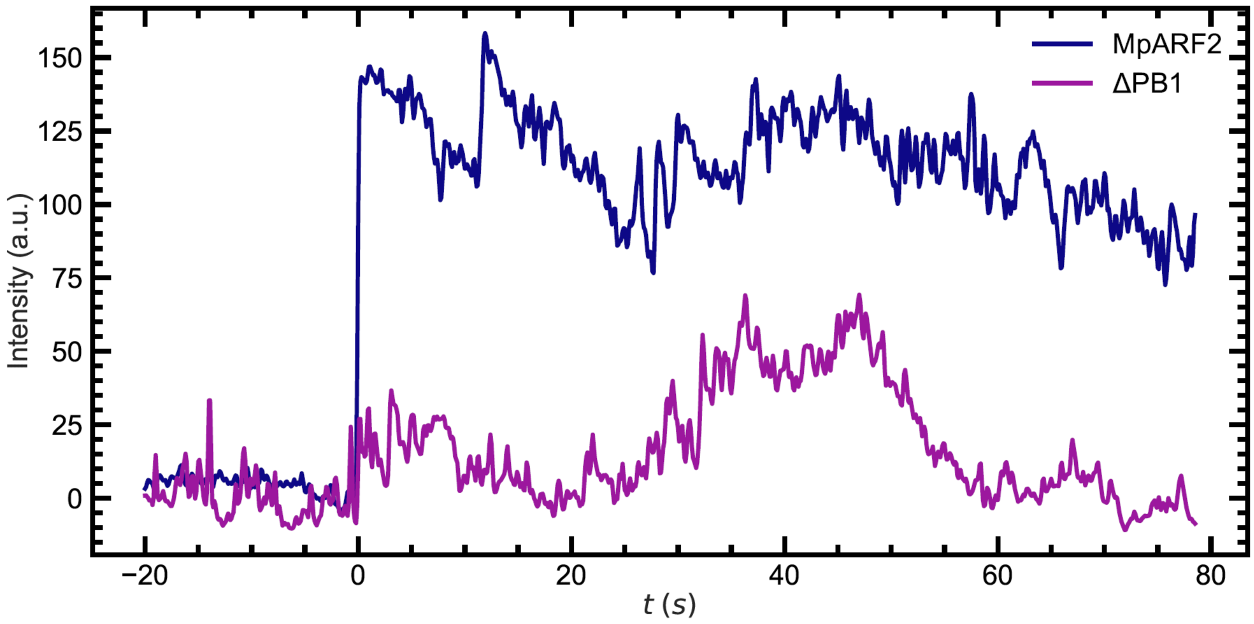


**Fig. S14. Binding kinetics in single-molecule imaging**

Representative time traces of protein intensity at the location of binding to DNA. $t_0$ is chosen as the first frame where DNA-binding is visible by eye. MpARF2 clusters bind stably and instantaneously, yielding a sudden jump in intensity, whereas the ΔPB1 protein variant exhibits gradual and transient binding.

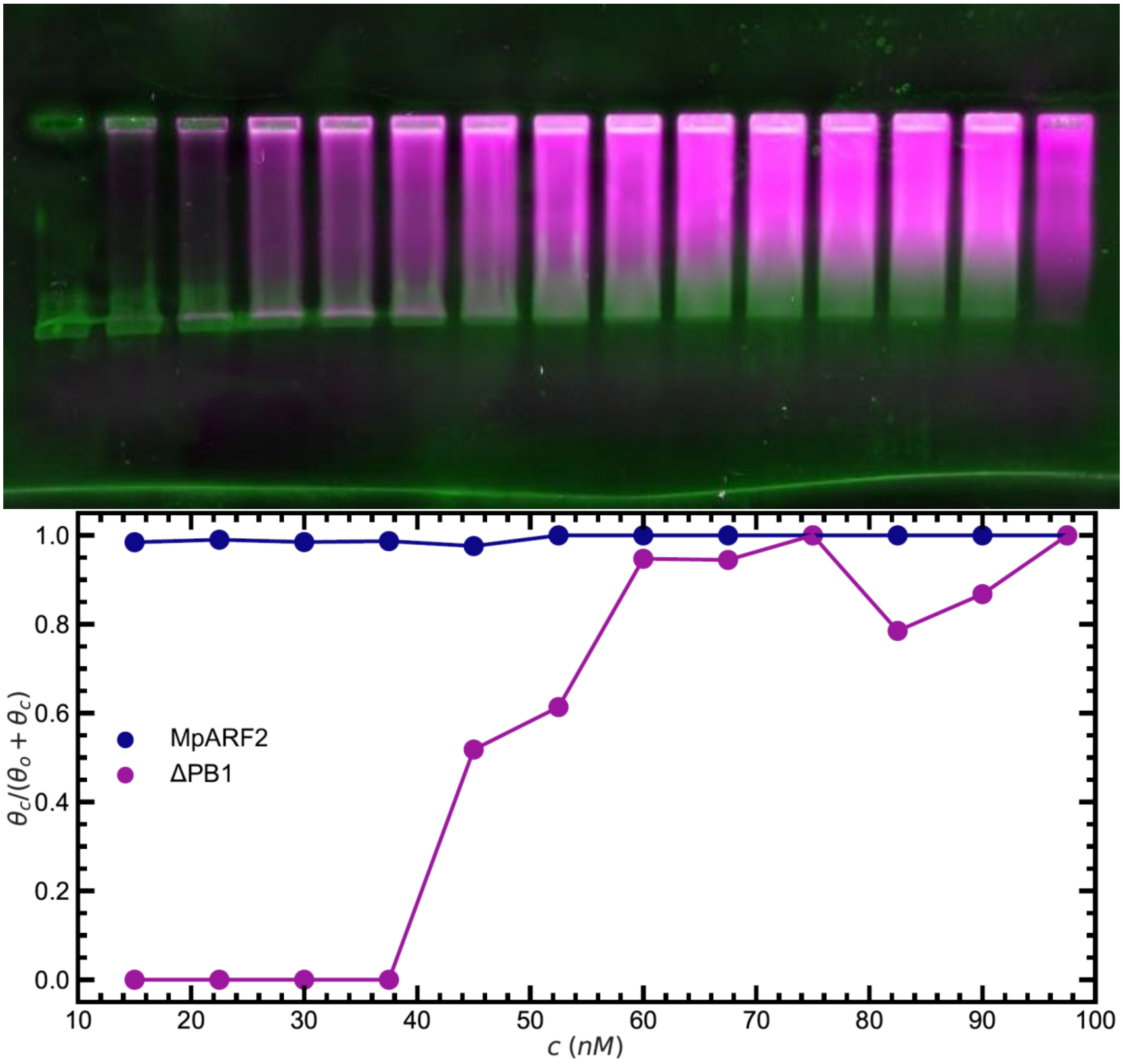


**Fig. S15. EMSA of ΔPB1 on IR7b1500 DNA**

Binding transition of the ΔPB1 protein variant to 1500 bp DNA **Top** EMSA gel of ΔPB1 (magenta, log color scale) and IR7b1500 DNA (green). Protein concentration range: 7.5 – 97.5 nM. The first and last wells are controls, containing only DNA and protein, respectively. **Bottom** EMSA quantification comparing the relative abundances of the oligomeric bound state $\theta_o$ and the clustered bound state, $\theta_c$. MpARF2 binds only as clusters which remain in the well of the gel (blue). In contrast, ΔPB1 (purple) binds primarily as oligomers at low concentrations (<40 nM) and switches to a clustered binding mode at higher concentrations.

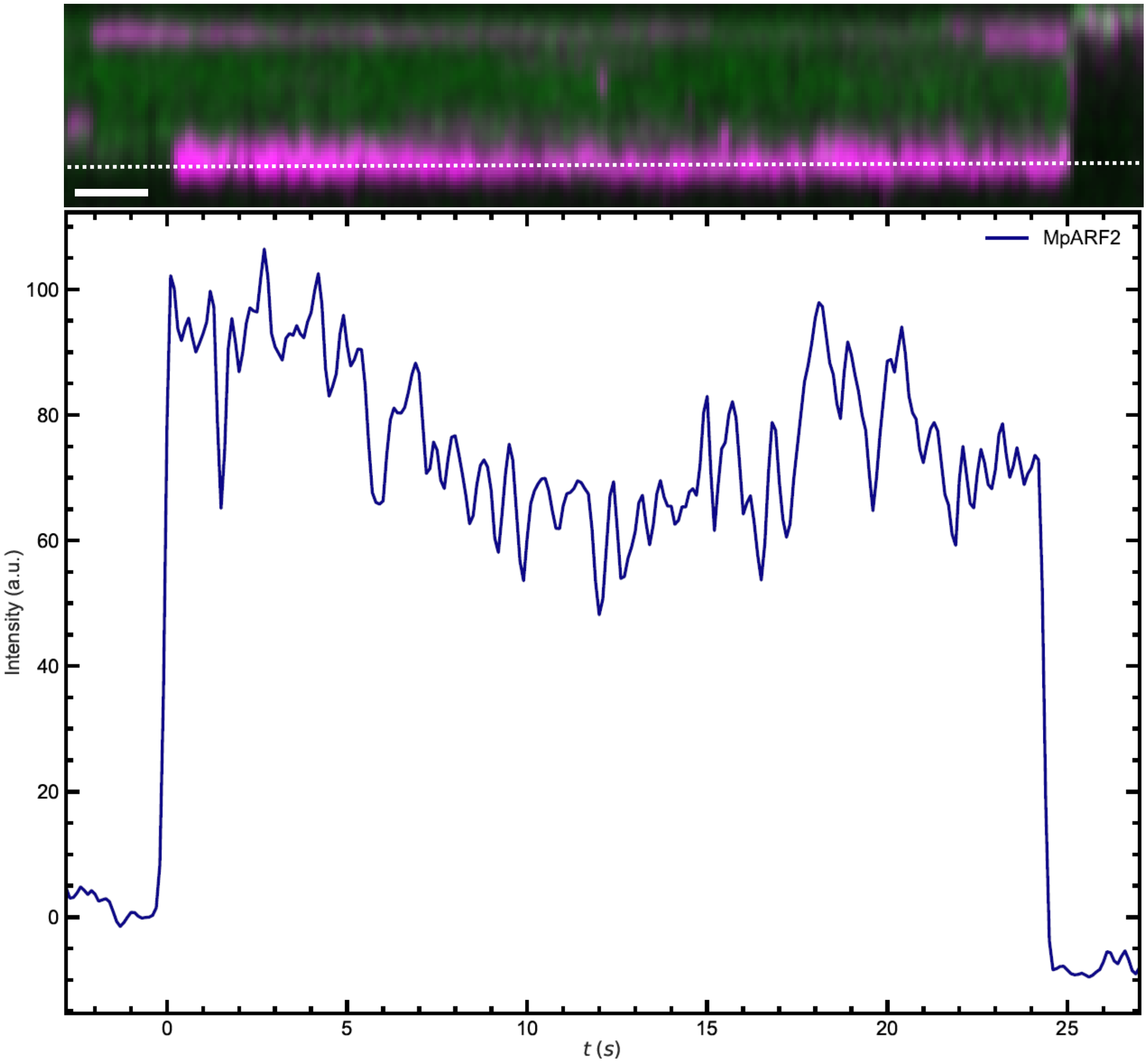


**Fig. S16. Rare example of an MpARF2 nanocluster binding to nonspecific DNA.**

**Top** Kymograph of an MpARF2 nanocluster (magenta) binding to nonspecific DNA (NSkb12, green), as observed using our single-molecule imaging assay. Scale bar: 2 s. **Bottom** Fluorescence intensity time trace along the dotted line in the top figure. Flow is interrupted at $t \approx 24$ s.

**Table S1. dsDNA sequences**

Linearized 3IR7kb12 and NSkb12 are identical to their plasmid sequences in Table S4. AuxREs occuring as part of an IR7 motif are highlighted in orange.

| Name | Sequence (5'-3') |
|---|---|
| MCS | GAGGTATTATTTCAAGGACCGGGATCCGAATTCGAGCGCCGTCGACAAGCTTGCGGCCGCACTGCAGGAAAACCTGTACTTCCAAT |
| UC19' | ACATGTGTTTTTCCATAGGCTCCGCCCCCCTGACGAGCATCACAAAAATCGACGCTCAAGTCAGAGGTGGCGAAACCCGAGAGGACTA<br>TAAAGATACCAGGCGTTTCCCCCTGGAAGCTCCCTCGTGCGCTCTCCTGTTCCGACCCTGCCGCTTACCGGATACCTGACCGCCTTTC<br>TCCCTTCGGGAAGCGTGGCGCTTTCTCATAGCTCACGCTGTAGGTATCTCAGTTCGGTGTAGGTCGTTCGCTCCAAGCTGGGCTGTGT<br>GCACGAACCCCCCGTTCAGCCCGACCGCTGCGCCTTATCCGGTAACTATCGTCTTGAGTCCAACCCGGTATGACACGACTTATCGCCA<br>CTGGCAGCAGCCACTGGTAACAGGATTAGCAGAGCGAGGTATGTAGGCGGTGCTACAGAGTTCTTGAAGTGGTGGCCTAACTACGGCT<br>ACACTAGAAGGACAGTATTTGGTATCTGCGCTCTGCTGAAGCCAGTTACCTTCGGAAAAAGAGTTGGTAGCTCTTGATCCGGCAAACA<br>AACCACCGCTGGTAGCGGTGGTTTTTTTGTTTGCAAGCAGCAGATTACGCGCAGAAAAAAAGGATCTCAAGAAGATCCTTTGATCTTT<br>TCTACTACCAATGCTTAATCAGTGAGGCACCTATCTCAGCGATCTGCCTATTTCGTTCATCCATAGTTGCCTGACTCCCCGTCGTGTA<br>GATAACTACGATACGGGAGGGCTTACCATCTGGCCCCAGTGCTGCAATGATACCGCGAGACCCACGCTCACCGGCTCCAGATTTATCA<br>GCAATAAACCAGCCAGCCGGAAGGGCCGAGCGCAGAAGTGGTCCTGCAACTTTATCCGCCTCCATCCAGTCTATTAATTGTTGCCGGG<br>AAGCTAGAGTAAGTAGTTCGCCAGTTAATAGTTTGCGCAACGTTGTTGCCATTGCTACAGGCATCGTGGTATCACGCTCGTCGTTTGG<br>TATGGCTTCATTCAGCTCCGGTTCCCAACGATCAAGGCGAGTTACATGATCCCCCATGTTGTGCAAAAAAGCGGTTAGCTCCTTCGGT<br>CCTCCGATCGTTGTTAGAAGTAAGTTGGCCGCAGTGTTATCACTCATGGTTATGGCAGCACTGCATAATTCTCTTACCGTCATGCCAT<br>CCGTAAGATGCTTTTCTGTGACTGGTGAGTACTCAACCAAGTCATTCTGAGAATAGTGTATGCGGCGACCGAGTTGCTCTTGCCCGGC<br>GTCAATACGGGATAATACCGCGCCACATAGCAGAACTTTAAAAGTGCTCATCATTGGAAAACGTTCTTCGGGGCGAAAACTCTCAAGG<br>ATCTTACCGCTGTTGAGATCCAGTTCGATGTAACCCACTCGTGCACCCAACTGATCTTCAGCATCTTTTACTTTCACCAGCGTTTCTG<br>GGTGAGCAAAAACAGGAAGGCAAAATGCCGCAAAAAAGGGAATAAGGGCTACCCGGAAATGTTGAATACTCATACTCTTC |
| IR7b19 | TGTCGGCCAAAGGCCGACA |
| NSb19 | TGGCGGGAAAAGTTCAGTT |
| IR7b1500 | GAAGAAGATGGAGCCCTAGATAACGATGATGATGACGATGATGATGATGATGACTCTGAAATGGAGAATCGTGATCGTTTGATTAGGA<br>AATCGAGAAGCCGTGGAGGTAGTACTAGAGGAAATAGGACGACGATTGAAGATCATCATCTTCAGGAGGAGAAAGCTCCGCCACCTCC<br>CCCTTTGGCGAATTCACGGCCAATTCCGCCGCCACGTCAGCATCAGCATCAACATCAGCAACAGCAACAACAACCTTTCTACGATTAC<br>TTCTTCCCTAATGTTGAGAATATGCCTGGAACTACTTTAGAAGATACTCCTCCACAACCACAACCACAACCAACAAGGCCTGTGCCTC<br>CTCAACCACATTCACCAGTCGTTACTGAGGATGACGAAGATGAGGAGGAGGAAGAGGAGGAAGAGGAGGAGGAAGAGGAGACGGTGAT<br>TGAACGGAAACCACTGGTGGAGGAAAGACCGAAGAGAGTAGAGGAAGTGACGATTGAATTGGAAAAAGTTACTAATTTGAGAGGGATG<br>AAGAAGAGTAAAGGGATAGGGATTCCCGGAGAGAGGAGAGGAATGCGAATGCCGGTGACTGCGACGCATTTGGCGAATGTATTCATTG<br>AGCTTGATGATAATTTCTTGAAAGCTTCTGAAAGTGCTCATGATGTTTCTAAGATGCTTGAAGCTACTAGGCTCCATTACCATTCTAA<br>TTTTGCAGATAACCGAGGTAATTTTGATTCGATAAATGTCGGACGTGCACCGACATCTTTATCCTTTTTCTTTTGATGCTTCATTCTC<br>TCTTTTCTTTTTTTTCCTCTGTAGCGTTTCTAGGAATCTCTCTCCAACGTTTAGGTTTCTTCTCAGATTTAGAAAGTTTCTTCTCTAT<br>CTGAAGAATTTTCATTTCGATTTCGCTTCTTCCTTCACCCTAATCCGTAATCGAATGCCATTTCGAAGATTGTGTTGAAGAGTAGAGA<br>GGAAGAAGAAGACTAAGAAGCTAGGGTTTCTCTGATGAGTAGATGATGCGTGAACCGCTTGTTAGCGAAGAAGAAGAAGAAGAAGCTA<br>CTGAAGTTTTACTAGTGGAGAAGACGAAGTTGTGTAAGAGACGAGGAGACGAAGAAAAAACGGAAGAGAGAAGAGACGATTTGTTGTT<br>ACTAGCTTTAACGCCGATGGTACGATCGAAATCTCAAGGAACTACTCGGCGCGTCACTCCCACGCCACCTCCTGTTGACGTGGAGAAA<br>CCGTTACCAAACGGAGATCTCTACATGGGAACGTTTTCCGGCGGGTTTCCCAACGGATCCGGTAAGTATTTGTGGAAGGATGGGTGTA<br>TGTACGAAGGCGAGTGGAAACGTGGTAAAGCGAGTGGTAAAGGCAAGTTTTCGTGGCCGAGTGGTGCGACTTATGAAGGAGAGTTCAA<br>ATCTGGGAGAATGGAAGGATCTGGGACTTTTGTTGGTGTTGATGGTGATACTTATCGTGGCTCTTGGGTTGCTGATCGGAAACAAGGT<br>CATG |
| NSb1500 | GAAGAAGATGGAGCCCTAGATAACGATGATGATGACGATGATGATGATGATGACTCTGAAATGGAGAATCGTGATCGTTTGATTAGGA<br>AATCGAGAAGCCGTGGAGGTAGTACTAGAGGAAATAGGACGACGATTGAAGATCATCATCTTCAGGAGGAGAAAGCTCCGCCACCTCC<br>CCCTTTGGCGAATTCACGGCCAATTCCGCCGCCACGTCAGCATCAGCATCAACATCAGCAACAGCAACAACAACCTTTCTACGATTAC<br>TTCTTCCCTAATGTTGAGAATATGCCTGGAACTACTTTAGAAGATACTCCTCCACAACCACAACCACAACCAACAAGGCCTGTGCCTC<br>CTCAACCACATTCACCAGTCGTTACTGAGGATGACGAAGATGAGGAGGAGGAAGAGGAGGAAGAGGAGGAGGAAGAGGAGACGGTGAT<br>TGAACGGAAACCACTGGTGGAGGAAAGACCGAAGAGAGTAGAGGAAGTGACGATTGAATTGGAAAAAGTTACTAATTTGAGAGGGATG<br>AAGAAGAGTAAAGGGATAGGGATTCCCGGAGAGAGGAGAGGAATGCGAATGCCGGTGACTGCGACGCATTTGGCGAATGTATTCATTG<br>AGCTTGATGATAATTTCTTGAAAGCTTCTGAAAGTGCTCATGATGTTTCTAAGATGCTTGAAGCTACTAGGCTCCATTACCATTCTAA<br>TTTTGCAGATAACCGAGGTAATTTTGATTCGATAAATCTTTATCCTTTTTCTTTTGATGCTTCATTCTCTCTTTTCTTTTTTTTCCTC<br>TGTAGCGTTTCTAGGAATCTCTCTCCAACGTTTAGGTTTCTTCTCAGATTTAGAAAGTTTCTTCTCTATCTGAAGAATTTTCATTTCG<br>ATTTCGCTTCTTCCTTCACCCTAATCCGTAATCGAATGCCATTTCGAAGATTGTGTTGAAGAGTAGAGAGGAAGAAGAAGACTAAGAA<br>GCTAGGGTTTCTCTGATGAGTAGATGATGCGTGAACCGCTTGTTAGCGAAGAAGAAGAAGAAGAAGCTACTGAAGTTTTACTAGTGGA<br>GAAGACGAAGTTGTGTAAGAGACGAGGAGACGAAGAAAAAACGGAAGAGAGAAGAGACGATTTGTTGTTACTAGCTTTAACGCCGATG<br>GTACGATCGAAATCTCAAGGAACTACTCGGCGCGTCACTCCCACGCCACCTCCTGTTGACGTGGAGAAACCGTTACCAAACGGAGATC<br>TCTACATGGGAACGTTTTCCGGCGGGTTTCCCAACGGATCCGGTAAGTATTTGTGGAAGGATGGGTGTATGTACGAAGGCGAGTGGAA<br>ACGTGGTAAAGCGAGTGGTAAAGGCAAGTTTTCGTGGCCGAGTGGTGCGACTTATGAAGGAGAGTTCAAATCTGGGAGAATGGAAGGA<br>TCTGGGACTTTTGTTGGTGTTGATGGTGATACTTATCGTGGCTCTTGGGTTGCTGATCGGAAACAAGGTCATG |

### Table S2. ssDNA sequences

| Set name | Code | Sequence |
|---|---|---|
| pET28 | BPJ030 | AGAAGAACCAGAAGAACCCATGGTATATCTCCTTC |
| | BPJ031 | AAATCTTCTGGTCACCATCACCATCACCATTGAGATCCGGCTGCTAAC |
| MBP | BPJ002 | TCCTTGAAATAATACCTCTAGCTCGATCCCATTAGTCTGCG |
| | BPJ029 | GGTTCTTCTGGTTCTTCTATGAAAATCGAAGAAGG |
| mNG | BPJ005 | AAAACCTGTACTTCCAATCGGACGAGATGGTGAGCAAGGGCGA |
| | BPJ032 | ATGGTGATGGTGATGGTGACCAGAAGATTTCTTGTACAGCTCGTCCATGC |
| pET_MCS | BPJ161 | AAGCTTGCGGCCGCACT |
| | BPJ162 | TTCGGATCCCGGTCCTTGAAAT |
| MpARF2 | BPJ163 | CAAGGACCGGGATCCGAAATGTCAGAAGCA |
| | BPJ166 | CAGTGCGGCCGCAAGCTTCATGTCGTCGCCGCG |
| DBD | BPJ163 | CAAGGACCGGGATCCGAAATGTCAGAAGCA |
| | BPJ165 | CAGTGCGGCCGCAAGCTTGAAGGGCTCCACTTCCCATG |
| MR | BPJ013 | GAATTCTCTCCCTCCACGACCATCA |
| | BPJ014 | AAGCTTATTGGATCTCTGGGGACCC |
| ΔDBD | BPJ013 | GAATTCTCTCCCTCCACGACCATCA |
| | BPJ043 | AAGCTTCATGTCGTCGCCGCGA |
| ΔMR1 | BPJ061 | GACGTCTTGAAGGGCTCCACTTCCCATGGT |
| | BPJ163 | CAAGGACCGGGATCCGAAATGTCAGAAGCA |
| ΔMR2 | BPJ062 | GAGCCCTTCAAGACGTCGCAAAATTTCCAACAAG |
| | BPJ166 | CAGTGCGGCCGCAAGCTTCATGTCGTCGCCGCG |
| ΔPB1 | BPJ163 | CAAGGACCGGGATCCGAAATGTCAGAAGCA |
| | BPJ164 | CAGTGCGGCCGCAAGCTTATTGGATCTCTGGGGACCCTTATC |
| A1 | KMA065 | TCGACGGTACCGCGGGCCCGATCTCTCTATTCGCTATTCCCAGATC |
| | KMA066 | ATTATAATGCCCACCACAGCTGATTGTATGAG |
| A2 | KMA067 | GCTGTGGTGGGCATTATAATCCTTATATTACTAGTGAGTAGTGAC |
| | KMA068 | TTGCTCACCATGGTGGCGACTTTATCGAATCAAAATTACCTCGGTTATCTG |
| B1 | KMA069 | TCGACGGTACCGCGGGCCCGTCTTTATCCTTTTTCTTTTGATGCTTCATTC |
| | KMA070 | TACAAAAATGGAGTACCTAAAAATTCACAACTTGGTTAGA |
| B2 | KMA071 | TTAGGTACTCCATTTTTGTAAATATGTACGTACACATAATGCT |
| | KMA072 | TTGCTCACCATGGTGGCGACAGTCATCTTCTCCACACTTTTCATTG |
| C1 | KMA081 | TCGACGGTACCGCGGGCCCGGATTATACCCACTGAAGGGTATTTCCG |
| | KMA082 | GAAGATGACTTCCGGTCTCAGACTCTCTTAAAAC |
| C2 | kMA083 | TGAGACCGGAAGTCATCTTCTCCACACTTTTCATTG |
| | KMA084 | TTGCTCACCATGGTGGCGACCATTTTTGTAAATATGTACGTACACATAATGCT |
| filler | KMA033 | CATTATTATCACCCTTTTCATGATGACCCTT |
| | JJR537 | CCTATGGAAAAACACATGTGCTCTTCACCCTAGCGCTGGAAACTGAACTTTTCCCGCC |
| pUC19’ | JJR547 | ACATGTGTTTTTCCATAGGCTCCGCC |
| | KMA035 | TGAAAAGGGTGATAATAATGGTTTCTTAGACGTCAGGTG |
| backbone | KMA117 | TACAAAAATGAGCGCTAGGGTGAAGAGCACATG |
| | KMA096 | ATAGAGAGATTGTCGGATGCGTACCGACAGCTGGAAACTGAACTTTTCCCGCC |
| A_IR7 | KMA097 | AGTTTCCAGCTGTCGGTACGCATCCGACAATCTCTCTATTCGCTATTCCCAGATC |
| | KMA098 | AGGATAAAGATGTCGGTGCACGTCCGACATTTATCGAATCAAAATTACCTCGGTTATCTG |
| B_IR7 | KMA099 | ATTCGATAAATGTCGGACGTGCACCGACATCTTTATCCTTTTTCTTTTGATGCTTCATTC |
| | KMA100 | GGGTATAATCTGTCGGGTCACTGCCGACAAGTCATCTTCTCCACACTTTTCATTG |
| C_IR7 | KMA101 | GAAGATGACTTGTCGGCAGTGACCCGACAGATTATACCCACTGAAGGGTATTTCCG |
| | KMA102 | TCACCCTAGCGCTCATTTTTGTAAATATGTACGTACACATAATGCT |
| A_NS | KMA119 | AGTTTCCAGCATCTCTCTATTCGCTATTCCCAGATC |
| | KMA120 | ATAGAGAGATGCTGGAAACTGAACTTTTCCCGCC |
| B_NS | KMA121 | ATTCGATAAATCTTTATCCTTTTTCTTTTGATGCTTCATTC |
| | KMA122 | AGGATAAAGATTTATCGAATCAAAATTACCTCGGTTATCTG |
| C_NS | KMA123 | GAAGATGACTGATTATACCCACTGAAGGGTATTTCCG |
| | KMA124 | TCACCCTAGCGCTCATTTTTGTAAATATGTACGTACACATAATGCT |
| b1500 | KMA130 | GAAGAAGATGGAGCCCTAGATAACG |
| | KMA131 | CATGACCTTGTTTCCGATCAGC |
| λ-DNA | CosR-bio | /5Phos/AGGTCGCCGCCCAAAAAAAAAAAA/3Bio/ |
| | CosL | /5Phos/GGGCGGCGACCTAAAAAAAAAAAA |

**Table S3. Strategies used to generate MpRF2 variant plasmids**

Overview of the methods employed to construct expression plasmids encoding MpARF2 protein variants. The table specifies the amino acid residues included in each variant relative to the full-length protein, the linearization strategy used for the host plasmid, and the primer sets applied for insert amplification.

| Protein variant | Residues | Linearization | Primers |
|---|---|---|---|
| MpARF2 | M1-M878 | BPJ161/BPJ162 | BPJ163/BPJ166 |
| MpARF2-DBD | M1-F396 | BPJ161/BPJ162 | BPJ163/BPJ165 |
| MpARF2-MR | S397-N744 | EcoRI+HindIII | BPJ013/BPJ014 |
| ΔDBD | S397-M878 | EcoRI+HindIII | BPJ013/BPJ043 |
| ΔMR | M1-F396 & K745-M878 | BPJ161/BPJ162 | BPJ061/BPJ163<br>BPJ062/BPJ166 |
| ΔPB1 | M1-N744 | BPJ161/BPJ162 | BPJ163/BPJ164 |

**Table S4. Overview of plasmids used for generating protein variants and DNA templates**

AuxREs occuring as part of an IR7 motif are highlighted in orange.

| Name | Sequence |
|---|---|
| pET_MBP-MCS-mNG-His | CGGGATCTCGACGCTCTCCCTTATGCGACTCCTGCATTAGGAAGCAGCCCAGTAGTAGGTTGAGGCCGTTGAGCACCGCCGCCGCAAGGAATGGTGCATGC<br>AAGGAGATGGCGCCCAACAGTCCCCCGGCCACGGGGCCTGCCACCATACCCACGCCGAAACAAGCGCTCATGAGCCCGAAGTGGCGAGCCCGATCTTCCCC<br>ATCGGTGATGTCGGCGATATAGGCGCCAGCAACCGCACCTGTGGCGCCGGTGATGCCGGCCACGATGCGTCCGGCGTAGAGGATCGAGATCTCGATCCCGC<br>GAAATTAATACGACTCACTATAGGGGAATTGTGAGCGGATAACAATTCCCCTCTAGAAATAATTTTGTTTAACTTTAAGAAGGAGATATACCATGGGTTCT<br>TCTGGTTCTTCTATGAAAATCGAAGAAGGTAAACTGGTAATCTGGATTAACGGCGATAAAGGCTATAACGGTCTCGCTGAAGTCGGTAAGAAATTCGAGAA<br>AGATACCGGAATTAAAGTCACCGTTGAGCATCCGGATAAACTGGAAGAGAAATTCCCACAGGTTGCGGCAACTGGCGATGGCCCTGACATTATCTTCTGGG<br>CACACGACCGCTTTGGTGGCTACGCTCAATCTGGCCTGTTGGCTGAAATCACCCCGGACAAAGCGTTCCAGGACAAGCTGTATCCGTTTACCTGGGATGCC<br>GTACGTTACAACGGCAAGCTGATTGCTTACCCGATCGCTGTTGAAGCGTTATCGCTGATTTATAACAAAGATCTGCTGCCGAACCCGCCAAAAACCTGGGA<br>AGAGATCCCGGCGCTGGATAAAGAACTGAAAGCGAAAGGTAAGAGCGCGCTGATGTTCAACCTGCAAGAACCGTACTTCACCTGGCCGCTGATTGCTGCTG<br>ACGGGGGTTATGCGTTCAAGTATGAAAACGGCAAGTACGACATTAAAGACGTGGGCGTGGATAACGCTGGCGCGAAAGCGGGTCTGACCTTCCTGGTTGAC<br>CTGATTAAAAACAAACACATGAATGCAGACACCGATTACTCCATCGCAGAAGCTGCCTTTAATAAAGGCGAAACAGCGATGACCATCAACGGCCCGTGGGC<br>ATGGTCCAACATCGACACCAGCAAAGTGAATTATGGTGTAACGGTACTGCCGACCTTCAAGGGTCAACCATCCAAACCGTTCGTTGGCGTGCTGAGCGCAG<br>GTATTAACGCCGCCAGTCCGAACAAAGAGCTGGCAAAAGAGTTCCTCGAAAACTATCTGCTGACTGATGAAGGTCTGGAAGCGGTTAATAAAGACAAACCG<br>CTGGGTGCCGTAGCGCTGAAGTCTTACGAGGAAGAGTTGGCGAAAGATCCACGTATTGCCGCCACTATGGAAAACGCCCAGAAAGGTGAAATCATGCCGAA<br>CATCCCGCAGATGTCCGCTTTCTGGTATGCCGTGCGTACTGCGGTGATCAACGCCGCCAGCGGTCGTCAGACTGTCGATGAAGCCCTGAAAGACGCGCAGA<br>CTAATGGGATCGAGCTAGAGGTATTATTTCAAGGACCGGGATCCGAATTCGAGCGCCGTCGACAAGCTTGCGGCCGCACTGCAGGAAAACCTGTACTTCCA<br>ATCGGACGAGATGGTGAGCAAGGGCGAGGAGGATAACATGGCCTCTCTCCCAGCGACACATGAGTTACACATCTTTGGCTCCATCAACGGTGTGGACTTTG<br>ACATGGTGGGTCAGGGCACCGGCAATCCAAATGATGGTTATGAGGAGTTAAACCTGAAGTCCACCAAGGGTGACCTCCAGTTCTCCCCCTGGATTCTGGTC<br>CCTCATATCGGGTATGGCTTCCATCAGTACCTGCCCTACCCTGACGGGATGTCGCCTTTCCAGGCCGCCATGGTAGATGGCTCCGGATACCAAGTCCATCG<br>CACAATGCAGTTTGAAGATGGTGCCTCCCTTACTGTTAACTACCGCTACACCTACGAGGGAAGCCACATCAAAGGAGAGGCCCAGGTGAAGGGGACTGGTT<br>TCCCTGCTGACGGTCCTGTGATGACCAACTCGCTGACCGCTGCGGACTGGTGCAGGTCGAAGAAGACTTACCCCAACGACAAAACCATCATCAGTACCTTT<br>AAGTGGAGTTACACCACTGGAAATGGCAAGCGCTACCGGAGCACTGCGCGGACCACCTACACCTTTGCCAAGCCAATGGCGGCTAACTATCTGAAGAACCA<br>GCCGATGTACGTGTTCCGTAAGACGGAGCTCAAGCACTCCAAGACCGAGCTCAACTTCAAGGAGTGGCAAAAGGCCTTTACCGATGTGATGGGCATGGACG<br>AGCTGTACAAGAAATCTTCTGGTCACCATCACCATCACCATTGAGATCCGGCTGCTAACAAAGCCCGAAAGGAAGCTGAGTTGGCTGCTGCCACCGCTGAG<br>CAATAACTAGCATAACCCCTTGGGGCCTCTAAACGGGTCTTGAGGGGTTTTTTGCTGAAAGGAGGAACTATATCCGGATTGGCGAATGGGACGCGCCCTGT<br>AGCGGCGCATTAAGCGCGGCGGGTGTGGTGGTTACGCGCAGCGTGACCGCTACACTTGCCAGCGCCCTAGCGCCCGCTCCTTTCGCTTTCTTCCCTTCCTT<br>TCTCGCCACGTTCGCCGGCTTTCCCCGTCAAGCTCTAAATCGGGGGCTCCCTTTAGGGTTCCGATTTAGTGCTTTACGGCACCTCGACCCCAAAAAACTTG<br>ATTAGGGTGATGGTTCACGTAGTGGGCCATCGCCCTGATAGACGGTTTTTCGCCCTTTGACGTTGGAGTCCACGTTCTTTAATAGTGGACTCTTGTTCCAA<br>ACTGGAACAACACTCAACCCTATCTCGGTCTATTCTTTTGATTTATAAGGGATTTTGCCGATTTCGGCCTATTGGTTAAAAAATGAGCTGATTTAACAAAA<br>ATTTAACGCGAATTTTAACAAACTAGTAACGTTTACAATTTCAGGTGGCACTTTTCGGGGAAATGTGCGCGGAACCCCTATTTGTTTATTTTTCTAAATAC<br>ATTCAAATATGTATCCGCTCATGAATTAATTCTTAGAAAAACTCATCGAGCATCAAATGAAACTGCAATTTATTCATATCAGGATTATCAATACCATATTT<br>TTGAAAAAGCCGTTTCTGTAATGAAGGAGAAAACTCACCGAGGCAGTTCCATAGGATGGCAAGATCCTGGTATCGGTCTGCGATTCCGACTCGTCCAACAT<br>CAATACAACCTATTAATTTCCCCTCGTCAAAAATAAGGTTATCAAGTGAGAAATCACCATGAGTGACGACTGAATCCGGTGAGAATGGCAAAAGTTTATGC<br>ATTTCTTTCCAGACTTGTTCAACAGGCCAGCCATTACGCTCGTCATCAAAATCACTCGCATCAACCAAACCGTTATTCATTCGTGATTGCGCCTGAGCGAG<br>ACGAAATACGCGATCGCTGTTAAAAGGACAATTACAAACAGGAATCGAATGCAACCGGCGCAGGAACACTGCCAGCGCATCAACAATGTTTTCACCTGAAT<br>CAGGATATTCTTCTAATACCTGGAATGCTGTTTTCCCGGGGATCGCAGTGGTGAGTAACCATGCATCATCAGGAGTACGGATAAAATGCTTGATGGTCGGA<br>AGAGGCATAAATTCCGTCAGCCAGTTTAGTCTGACCATCTCATCTGTAACATCATTGGCAACGCTACCTTTGCCATGTTTCAGAAACAACTCTGGCGCATC<br>GGGCTTCCCATACAATCGATAGATTGTCGCACCTGATTGCCCGACATTATCGCGAGCCCATTTATACCCATATAAATCAGCATCCATGTTGGAATTTAATC<br>GCGGCCTAGAGCAAGACGTTTCCCGTTGAATATGGCTCATAACACCCCTTGTATTACTGTTTATGTAAGCAGACAGTTTTATTGTTCATGACCAAAATCCC<br>TTAACGTGAGTTTTCGTTCCACTGAGCGTCAGACCCCGTAGAAAAGATCAAAGGATCTTCTTGAGATCCTTTTTTTCTGCGCGTAATCTGCTGCTTGCAAA<br>CAAAAAAACCACCGCTACCAGCGGTGGTTTGTTTGCCGGATCAAGAGCTACCAACTCTTTTTCCGAAGGTAACTGGCTTCAGCAGAGCGCAGATACCAAAT<br>ACTGTCCTTCTAGTGTAGCCGTAGTTAGGCCACCACTTCAAGAACTCTGTAGCACCGCCTACATACCTCGCTCTGCTAATCCTGTTACCAGTGGCTGCTGC<br>CAGTGGCGATAAGTCGTGTCTTACCGGGTTGGACTCAAGACGATAGTTACCGGATAAGGCGCAGCGGTCGGGCTGAACGGGGGGTTCGTGCACACAGCCCA<br>GCTTGGAGCGAACGACCTACACCGAACTGAGATACCTACAGCGTGAGCTATGAGAAAGCGCCACGCTTCCCGAAGGGAGAAAGGCGGACAGGTATCCGGTA<br>AGCGGCAGGGTCGGAACAGGAGAGCGCACGAGGGAGCTTCCAGGGGGAAACGCCTGGTATCTTTATAGTCCTGTCGGGTTTCGCCACCTCTGACTTGAGCG<br>TCGATTTTTGTGATGCTCGTCAGGGGGGCGGAGCCTATGGAAAAACGCCAGCAACGCGGCCTTTTTACGGTTCCTGGCCTTTTGCTGGCCTTTTGCTCACA<br>TGTTCTTTCCTGCGTTATCCCCTGATTCTGTGGATAACCGTATTACCGCCTTTGAGTGAGCTGATACCGCTCGCCGCAGCCGAACGACCGAGCGCAGCGAG<br>TCAGTGAGCGAGGAAGCGGAAGAGCGCCTGATGCGGTATTTTCTCCTTACGCATCTGTGCGGTATTTCACACCGCATATATGGTGCACTCTCAGTACAATC<br>TGCTCTGATGCCGCATAGTTAAGCCAGTATACACTCCGCTATCGCTACGTGACTGGGTCATGGCTGCGCCCCGACACCCGCCAACACCCGCTGACGCGCCC<br>TGACGGGCTTGTCTGCTCCCGGCATCCGCTTACAGACAAGCTGTGACCGTCTCCGGGAGCTGCATGTGTCAGAGGTTTTCACCGTCATCACCGAAACGCGC<br>GAGGCAGCTGCGGTAAAGCTCATCAGCGTGGTCGTGAAGCGATTCACAGATGTCTGCCTGTTCATCCGCGTCCAGCTCGTTGAGTTTCTCCAGAAGCGTTA<br>ATGTCTGGCTTCTGATAAAGCGGGCCATGTTAAGGGCGGTTTTTTCCTGTTTGGTCACTGATGCCTCCGTGTAAGGGGGATTTCTGTTCATGGGGGTAATG<br>ATACCGATGAAACGAGAGAGGATGCTCACGATACGGGTTACTGATGATGAACATGCCCGGTTACTGGAACGTTGTGAGGGTAAACAACTGGCGGTATGGAT<br>GCGGCGGGACCAGAGAAAAATCACTCAGGGTCAATGCCAGCGCTTCGTTAATACAGATGTAGGTGTTCCACAGGGTAGCCAGCAGCATCCTGCGATGCAGA<br>TCCGGAACATAATGGTGCAGGGCGCTGACTTCCGCGTTTCCAGACTTTACGAAACACGGAAACCGAAGACCATTCATGTTGTTGCTCAGGTCGCAGACGTT<br>TTGCAGCAGCAGTCGCTTCACGTTCGCTCGCGTATCGGTGATTCATTCTGCTAACCAGTAAGGCAACCCCGCCAGCCTAGCCGGGTCCTCAACGACAGGAG<br>CACGATCATGCGCACCCGTGGGGCCGCCATGCCGGCGATAATGGCCTGCTTCTCGCCGAAACGTTTGGTGGCGGGACCAGTGACGAAGGCTTGAGCGAGGG<br>CGTGCAAGATTCCGAATACCGCAAGCGACAGGCCGATCATCGTCGCGCTCCAGCGAAAGCGGTCCTCGCCGAAAATGACCCAGAGCGCTGCCGGCACCTGT<br>CCTACGAGTTGCATGATAAAGAAGACAGTCATAAGTGCGGCGACGATAGTCATGCCCCGCGCCCACCGGAAGGAGCTGACTGGGTTGAAGGCTCTCAAGGG<br>CATCGGTCGAGATCCCGGTGCCTAATGAGTGAGCTAACTTACATTAATTGCGTTGCGCTCACTGCCCGCTTTCCAGTCGGGAAACCTGTCGTGCCAGCTGC<br>ATTAATGAATCGGCCAACGCGCGGGGAGAGGCGGTTTGCGTATTGGGCGCCAGGGTGGTTTTTCTTTTCACCAGTGAGACGGGCAACAGCTGATTGCCCTT<br>CACCGCCTGGCCCTGAGAGAGTTGCAGCAAGCGGTCCACGCTGGTTTGCCCCAGCAGGCGAAAATCCTGTTTGATGGTGGTTAACGGCGGGATATAACATG<br>AGCTGTCTTCGGTATCGTCGTATCCCACTACCGAGATATCCGCACCAACGCGCAGCCCGGACTCGGTAATGGCGCGCATTGCGCCCAGCGCCATCTGATCG<br>TTGGCAACCAGCATCGCAGTGGGAACGATGCCCTCATTCAGCATTTGCATGGTTTGTTGAAAACCGGACATGGCACTCCAGTCGCCTTCCCGTTCCGCTAT<br>CGGCTGAATTTGATTGCGAGTGAGATATTTATGCCAGCCAGCCAGACGCAGACGCGCCGAGACAGAACTTAATGGGCCCGCTAACAGCGCGATTTGCTGGT<br>GACCCAATGCGACCAGATGCTCCACGCCCAGTCGCGTACCGTCTTCATGGGAGAAAATAATACTGTTGATGGGTGTCTGGTCAGAGACATCAAGAAATAAC<br>GCCGGAACATTAGTGCAGGCAGCTTCCACAGCAATGGCATCCTGGTCATCCAGCGGATAGTTAATGATCAGCCCACTGACGCGTTGCGCGAGAAGATTGTG<br>CACCGCCGCTTTACAGGCTTCGACGCCGCTTCGTTCTACCATCGACACCACCACGCTGGCACCCAGTTGATCGGCGCGAGATTTAATCGCCGCGACAATTT<br>GCGACGGCGCGTGCAGGGCCAGACTGGAGGTGGCAACGCCAATCAGCAACGACTGTTTGCCCGCCAGTTGTTGTGCCACGCGGTTGGGAATGTAATTCAGC<br>TCCGCCATCGCCGCTTCCACTTTTTCCCGCGTTTTCGCAGAAACGTGGCTGGCCTGGTTCACCACGCGGGAAACGGTCTGATAAGAGACACCGGCATACTC<br>TGCGACATCGTATAACGTTACTGGTTTCACATTCACCACCCTGAATTGACTCTCTTCCGGGCGCTATCATGCCATACCGCGAAAGGTTTTGCGCCATTCGA<br>TGGTGTC |

| | |
|---|---|
| pUC19’ | GCTAGGGTGAAGAGCACATGTGTTTTTCCATAGGCTCCGCCCCCCTGACGAGCATCACAAAAATCGACGCTCAAGTCAGAGGTGGCGAAACCCGACAGGAC<br>TATAAAGATACCAGGCGTTTCCCCCTGGAAGCTCCCTCGTGCGCTCTCCTGTTCCGACCCTGCCGCTTACCGGATACCTGTCCGCCTTTCTCCCTTCGGGA<br>AGCGTGGCGCTTTCTCATAGCTCACGCTGTAGGTATCTCAGTTCGGTGTAGGTCGTTCGCTCCAAGCTGGGCTGTGTGCACGAACCCCCCGTTCAGCCCGA<br>CCGCTGCGCCTTATCCGGTAACTATCGTCTTGAGTCCAACCCGGTAAGACACGACTTATCGCCACTGGCAGCAGCCACTGGTAACAGGATTAGCAGAGCGA<br>GGTATGTAGGCGGTGCTACAGAGTTCTTGAAGTGGTGGCCTAACTACGGCTACACTAGAAGAACAGTATTTGGTATCTGCGCTCTGCTGAAGCCAGTTACC<br>TTCGGAAAAAGAGTTGGTAGCTCTTGATCCGGCAAACAAACCACCGCTGGTAGCGGTGGTTTTTTTGTTTGCAAGCAGCAGATTACGCGCAGAAAAAAAGG<br>ATCTCAAGAAGATCCTTTGATCTTTTCTACGGGGTCTGACGCTCAGTGGAACGAAAACTCACGTTAAGGGATTTTGGTCATGAGATTATCAAAAAGGATCT<br>TCACCTAGATCCTTTTAAATTAAAAATGAAGTTTTAAATCAATCTAAAGTATATATGAGTAAACTTGGTCTGACAGTTACCAATGCTTAATCAGTGAGGCA<br>CCTATCTCAGCGATCTGTCTATTTCGTTCATCCATAGTTGCCTGACTCCCCGTCGTGTAGATAACTACGATACGGGAGGGCTTACCATCTGGCCCCAGTGC<br>TGCAATGATACCGCGAGACCCACGCTCACCGGCTCCAGATTTATCAGCAATAAACCAGCCAGCCGGAAGGGCCGAGCGCAGAAGTGGTCCTGCAACTTTAT<br>CCGCCTCCATCCAGTCTATTAATTGTTGCCGGGAAGCTAGAGTAAGTAGTTCGCCAGTTAATAGTTTGCGCAACGTTGTTGCCATTGCTACAGGCATCGTG<br>GTGTCACGCTCGTCGTTTGGTATGGCTTCATTCAGCTCCGGTTCCCAACGATCAAGGCGAGTTACATGATCCCCCATGTTGTGCAAAAAAGCGGTTAGCTC<br>CTTCGGTCCTCCGATCGTTGTCAGAAGTAAGTTGGCCGCAGTGTTATCACTCATGGTTATGGCAGCACTGCATAATTCTCTTACTGTCATGCCATCCGTAA<br>GATGCTTTTCTGTGACTGGTGAGTACTCAACCAAGTCATTCTGAGAATAGTGTATGCGGCGACCGAGTTGCTCTTGCCCGGCGTCAATACGGGATAATACC<br>GCGCCACATAGCAGAACTTTAAAAGTGCTCATCATTGGAAAACGTTCTTCGGGGCGAAAACTCTCAAGGATCTTACCGCTGTTGAGATCCAGTTCGATGTA<br>ACCCACTCGTGCACCCAACTGATCTTCAGCATCTTTTACTTTCACCAGCGTTTCTGGGTGAGCAAAAACAGGAAGGCAAAATGCCGCAAAAAAGGGAATAA<br>GGGCGACACGGAAATGTTGAATACTCATACTCTTCCTTTTTCAATATTATTGAAGCATTTATCAGGGTTATTGTCTCATGAGCGGATACATATTTGAATGT<br>ATTTAGAAAAATAAACAAATAGGGGTTCCGCGCACATTTCCCCGAAAAGTGCCACCTGACGTCTAAGAAACCATTATTATCACCCTTTTCATGATGACCCT<br>TTTCAACTCTATCCTTTTGTTTTATCCATTTTGTTTCTACCCTCTTTAAATTTTTAATGACCATTTTACCCTTATTTGGAAATTGTTTTAGATTAACAAAA<br>TGAAATATTAATATTTTTCCGCCTAAAATATTCCAAAATAAATTTCCCGCCAAAAATTTTCAAAAAAAAAATTCCGCCAAAAATTTTCAAAAAACAAATTC<br>CCGCCAAATTTTGTTTTCAAAAATATTTTTCCCGCCAATTTTTTTTTTGGCGGGAAATAGATTTACAAGGGATTTTTAAAGAGTTTTAAAAGATTTACAAG<br>AGATTTTAAAGGGGTTTTATAAGATTTACAAGAGATTTTAAAAGGGTGATTTACAAGGGATTTTAAAAGGTTTTTAAAATATTTACAAGAGATTTTAAAAG<br>GGTTTTAAAAGATTTACAAGAGATTTTAAAGAGATTTAAAAGATTTACAAAGGTCTTTTAAAGATTTTTAAACGGTTTAAAATTATTTTAAAATGATTTTA<br>AAATATTTACAAGGGATTTTAAAAGGGTTTTACAATATTTACAACAGATTTTAAAGGACTTTAAAATATTTACAAAAGATCTGTAAAGATTCTTAAAGGGT<br>TTAAAAGTGTTTTGAAAGATTTACAAGAGATTTTAAAACCTTTTAAAATTACTTGTAAATTTTTTAAAACCCTTGTAAATCTTTTAAAATCTCTTTTAAAT<br>CTTTTAAAACCTTTTTAAAATCTCTTGTAAATCTTTTAAGACCCTTTTAAAATCCCTTGTAAATCTATTTGCCGCAAACAAATATTTACAAGGATTTTTAA<br>AATTATTTTTTTGGCTCAGAAAAAAATTTGTTACGGAACAAAAAAATTTGGCGGGAAACTTTTTTTTTGGCGGGAAAAAAAGTTCGGCGGGAAATCTTTTTTGG<br>CGGGAAAAAAAAAATTTTGGCGCCAATAAATTTGGCGGGAAAAGTTCAGTTTCCAGC |
| pBlockA | TAGTTATTAATAGTAATCAATTACGGGGTCATTAGTTCATAGCCCATATATGGAGTTCCGCGTTACATAACTTACGGTAAATGGCCCGCCTGGCTGACCGC<br>CCAACGACCCCCGCCCATTGACGTCAATAATGACGTATGTTCCCATAGTAACGCCAATAGGGACTTTCCATTGACGTCAATGGGTGGAGTATTTACGGTAA<br>ACTGCCCACTTGGCAGTACATCAAGTGTATCATATGCCAAGTACGCCCCCTATTGACGTCAATGACGGTAAATGGCCCGCCTGGCATTATGCCCAGTACAT<br>GACCTTATGGGACTTTCCTACTTGGCAGTACATCTACGTATTAGTCATCGCTATTACCATGGTGATGCGGTTTTGGCAGTACATCAATGGGCGTGGATAGC<br>GGTTTGACTCACGGGGATTTCCAAGTCTCCACCCCATTGACGTCAATGGGAGTTTGTTTTGGCACCAAAATCAACGGGACTTTCCAAAATGTCGTAACAAC<br>TCCGCCCCATTGACGCAAATGGGCGGTAGGCGTGTACGGTGGGAGGTCTATATAAGCAGAGCTGGTTTAGTGAACCGTCAGATCCGCTAGCGCTACCGGAC<br>TCAGATCTCGAGCTCAAGCTTCGAATTCTGCAGTCGACGGTACCGCGGGCCCGATCTCTCTATTCGCTATTCCCAGATCTCCATGACCCACTTCACTCCCT<br>GCGCGTCCGGTACAATCGCCACTTCCATAACACCAAGAGTCAATACAAAAAAAAAGCCGACTCCATTATTATCCCTCTCTCTACAAAAAAAACAGTAGAGA<br>GAGATATAGAGAGCATCAATGGCGTCGTCGTCAAGACTTTGCGATTCGTGCAAATCGACGGCAGCAACTCTATTCTGCCGCGCAGACGCCGCGTTTCTCTG<br>CGGCGACTGCGACGGTAAGATCCACACAGCTAACAAACTCGCATCACGTCACGAACGAGTTTGGCTATGTGAAGTATGCGAACAAGCACCAGCACACGTCA<br>CGTGCAAAGCTGACGCCGCTGCGTTATGCGTCACGTGCGACCGTGATATCCACTCAGCTAATCCACTCTCTCGCCGCCACGAGCGCGTACCAATCACTCCT<br>TTCTACGACGCTGTAGGTCCAGCAAAATCCGCTTCCTCCTCCGTCAATTTCGTAGATGAAGATGGCGGTGACGTCACTGCTTCGTGGCTTTTAGCTAAAGA<br>AGGAATCGAAATCACTAATTTGTTCTCCGATCTTGATTATCCGAAGATTGAGGTAACGTCGGAGGAGAATAGCTCCGGTAACGACGGAGTTGTTCCTGTGC<br>AGAATAAGTTGTTTCTTAACGAAGATTACTTCAATTTCGATCTCTCTGCTTCCAAAATTTCTCAACAAGGATTCAATTTCATCAACCAAACTGTACGAATC<br>TCTCTCTATCTTCTTAAGCTTCCATGGATCTGGAAAATTTTCAGATCTAGATCTGAAAGTTTGGATCTGATTTGAGTTTGTGAAATGTTGTTAGGTTTCGA<br>CGAGAACGATAGATGTACCGTTGGTGCCTGAAAGTGGAGGAGTGACGGCGGAGATGACGAACACGGAGACGCCAGCTGTGCAGTTATCACCGGCGGAGAGG<br>GAAGCAAGGGTTTTGAGGTATAGAGAGAAGAGGAAGAATCGGAAATTTGAGAAGACGATTAGGTATGCGTCGCGTAAAGCTTACGCTGAGATGAGGCCGAG<br>AATCAAAGGACGTTTTGCTAAGAGAACAGATTCGAGAGAGAATGATGGTGGAGACGTCGGAGTTTATGGCGGATTCGGTGTTGTTCCGAGTTTCTGATATT<br>TTCCGGTGAAAGAAACATGGTAGTAGTATATCGGTTAATAGCAGATTAAGGATAATTAAAACTAAAATGTTTCCTGATTTAATCAGGGGCTAAGTTATTGT<br>TAGAAGATTGTTGTTTTTTGAATTAATTCTCACAATTGTATCCTAAATTATATAGTTTACTTAATGTTGAACTAATAAATTTTGATTTACAATCTCTGTAA<br>GAAATCGATATGCAAAAATAAATGAGTTGAGCTATTAAGATTAACATTCATATAGAACACAAACTCATACAATCAGCTGTGGTGGGAAAAAAAGCATTATA<br>ATCCTTATATTACTAGTGAGTAGTGACTGATACAAAAGATTTTTATACGTTAAAAGTTTAAAAATTCAAAAGTACAGAAAACGTTATATTCGCTTGTTAC<br>CTTTTTTAACTTTATCCACGTCATTACCCCTGAGAGTAGCCCAACGAAAAATGGAAAATTTAACCCAAAAAACGGAAAATATTAGAGAAAGCGTTAATGCC<br>TTTTCTAACAACTAAACGACGACGTATCAGTTCAATCCTCTCTCTATTATTATTCCTTAAAACTTAACTCTAAAGCTTTCGAGTATTCTTCAATCTCTTCT<br>TCTCCGATCGATCTGAAGTAAGAGAGCTTCTTAAGGAAGAAGAAGATGGGTTGTGCTCAATCAAAGATCGAGAACGAAGAAGCAGTTACTCGTTGCAAAGA<br>ACGAAAACAATTGATGAAAGACGCCGTCACTGCTCGTAACGCTTTCGCCGCCGCTCACTCAGCTTACGCTATGGCTCTTAAAAACACCGGAGCTGCTCTTT<br>CCGATTACTCTCACGGCGAGTTTTTAGTCTCTAATCACTCGTCTTCCTCCGCAGCTGCAGCAATCGCTTCTACTTCTTCTCTTCCCACTGCTATATCTCCT<br>CCTCTTCCTTCTTCCACCGCTCCGGTTTCTAATTCAACCGCTTCTTCTTCCTCCGCTGCGGTTCCTCAGCCGATTCCTGATACTCTTCCTCCTCCTCCTCC<br>TCCACCACCGCTTCCTCTTCAACGTGCTGCTACTATGCCGGAGATGAACGGTAGATCCGGTGGTGGTCATGCTGGTAGTGGACTCAACGGAATTGAAGAAG<br>ATGGAGCCCTAGATAACGATGATGATGACGATGATGATGATGATGACTCTGAAATGGAGAATCGTGATCGTTTGATTAGGAAATCGAGAAGCCGTGGAGGT<br>AGTACTAGAGGAAATAGGACGACGATTGAAGATCATCATCTTCAGGAGGAGAAAGCTCCGCCACCTCCCCCTTTGGCGAATTCACGGCCAATTCCGCCGCC<br>ACGTCAGCATCAGCATCAACATCAGCAACAGCAACAACAACCTTTCTACGATTACTTCTTCCCTAATGTTGAGAATATGCCTGGAACTACTTTAGAAGATA<br>CTCCTCCACAACCACAACCACAACCAACAAGGCCTGTGCCTCCTCAACCACATTCACCAGTCGTTACTGAGGATGACGAAGATGAGGAGGAGGAAGAGGAG<br>GAAGAGGAGGAGGAAGAGGAGACGGTGATTGAACGGAAACCACTGGTGGAGGAAAGACCGAAGAGAGTAGAGGAAGTGACGATTGAATTGGAAAAAGTTAC<br>TAATTTGAGAGGGATGAAGAAGAGTAAAGGGATAGGGATTCCCGGAGAGAGGAGAGGAATGCGAATGCCGGTGACTGCGACGCATTTGGCGAATGTATTCA<br>TTGAGCTTGATGATAATTTCTTGAAAGCTTCTGAAAGTGCTCATGATGTTTCTAAGATGCTTGAAGCTACTAGGCTCCATTACCATTCTAATTTTGCAGAT<br>AACCGAGGTAATTTTGATTCGATAAAGTCGCCACCATGGTGAGCAAGGGCGAGGAGCTGTTCACCGGGGTGGTGCCCATCCTGGTCGAGCTGGACGGCGAC<br>GTAAACGGCCACAAGTTCAGCGTGTCCGGCGAGGGCGAGGGCGATGCCACCTACGGCAAGCTGACCCTGAAGTTCATCTGCACCACCGGCAAGCTGCCCGT<br>GCCCTGGCCCACCCTCGTGACCACCCTGACCTACGGCGTGCAGTGCTTCAGCCGCTACCCCGACCACATGAAGCAGCACGACTTCTTCAAGTCCGCCATGC<br>CCGAAGGCTACGTCCAGGAGCGCACCATCTTCTTCAAGGACGACGGCAACTACAAGACCCGCGCCGAGGTGAAGTTCGAGGGCGACACCCTGGTGAACCGC<br>ATCGAGCTGAAGGGCATCGACTTCAAGGAGGACGGCAACATCCTGGGGCACAAGCTGGAGTACAACTACAACAGCCACAACGTCTATATCATGGCCGACAA<br>GCAGAAGAACGGCATCAAGGTGAACTTCAAGATCCGCCACAACATCGAGGACGGCAGCGTGCAGCTCGCCGACCACTACCAGCAGAACACCCCCATCGGCG<br>ACGGCCCCGTGCTGCTGCCCGACAACCACTACCTGAGCACCCAGTCCGCCCTGAGCAAAGACCCCAACGAGAAGCGCGATCACATGGTCCTGCTGGAGTTC<br>GTGACCGCCGCCGGGATCACTCTCGGCATGGACGAGCTGTACAAGTAAAGCGGCCGCGACTCTAGATCATAATCAGCCATACCACATTTGTAGAGGTTTTA<br>CTTGCTTTAAAAAACCTCCCACACCTCCCCCTGAACCTGAAACATAAAATGAATGCAATTGTTGTTGTTAACTTGTTTATTGCAGCTTATAATGGTTACAA<br>ATAAAGCAATAGCATCACAAATTTCACAAATAAAGCATTTTTTTCACTGCATTCTAGTTGTGGTTTGTCCAAACTCATCAATGTATCTTAAGGCGTAAATT<br>GTAAGCGTTAATATTTTGTTAAAATTCGCGTTAAATTTTTGTTAAATCAGCTCATTTTTTAACCAATAGGCCGAAATCGGCAAAATCCCTTATAAATCAAA<br>AGAATAGACCGAGATAGGGTTGAGTGTTGTTCCAGTTTGGAACAAGAGTCCACTATTAAAGAACGTGGACTCCAACGTCAAAGGGCGAAAAACCGTCTATC<br>AGGGCGATGGCCCACTACGTGAACCATCACCCTAATCAAGTTTTTTGGGGTCGAGGTGCCGTAAAGCACTAAATCGGAACCCTAAAGGGAGCCCCCGATTT<br>AGAGCTTGACGGGGAAAGCCGGCGAACGTGGCGAGAAAGGAAGGGAAGAAAGCGAAAGGAGCGGGCGCTAGGGCGCTGGCAAGTGTAGCGGTCACGCTGCG |

| | |
|---|---|
| | CGTAACCACCACACCCGCCGCGCTTAATGCGCCGCTACAGGGCGCGTCAGGTGGCACTTTTCGGGGAAATGTGCGCGGAACCCCTATTTGTTTATTTTTCT<br>AAATACATTCAAATATGTATCCGCTCATGAGACAATAACCCTGATAAATGCTTCAATAATATTGAAAAAGGAAGAGTCCTGAGGCGGAAAGAACCAGCTGT<br>GGAATGTGTGTCAGTTAGGGTGTGGAAAGTCCCCAGGCTCCCCAGCAGGCAGAAGTATGCAAAGCATGCATCTCAATTAGTCAGCAACCAGGTGTGGAAAG<br>TCCCCAGGCTCCCCAGCAGGCAGAAGTATGCAAAGCATGCATCTCAATTAGTCAGCAACCATAGTCCCGCCCCTAACTCCGCCCATCCCGCCCCTAACTCC<br>GCCCAGTTCCGCCCATTCTCCGCCCCATGGCTGACTAATTTTTTTTATTTATGCAGAGGCCGAGGCCGCCTCGGCCTCTGAGCTATTCCAGAAGTAGTGAG<br>GAGGCTTTTTTGGAGGCCTAGGCTTTTGCAAAGATCGATCAAGAGACAGGATGAGGATCGTTTCGCATGATTGAACAAGATGGATTGCACGCAGGTTCTCC<br>GGCCGCTTGGGTGGAGAGGCTATTCGGCTATGACTGGGCACAACAGACAATCGGCTGCTCTGATGCCGCCGTGTTCCGGCTGTCAGCGCAGGGGCGCCCGG<br>TTCTTTTTGTCAAGACCGACCTGTCCGGTGCCCTGAATGAACTGCAAGACGAGGCAGCGCGGCTATCGTGGCTGGCCACGACGGGCGTTCCTTGCGCAGCT<br>GTGCTCGACGTTGTCACTGAAGCGGGAAGGGACTGGCTGCTATTGGGCGAAGTGCCGGGGCAGGATCTCCTGTCATCTCACCTTGCTCCTGCCGAGAAAGT<br>ATCCATCATGGCTGATGCAATGCGGCGGCTGCATACGCTTGATCCGGCTACCTGCCCATTCGACCACCAAGCGAAACATCGCATCGAGCGAGCACGTACTC<br>GGATGGAAGCCGGTCTTGTCGATCAGGATGATCTGGACGAAGAGCATCAGGGGCTCGCGCCAGCCGAACTGTTCGCCAGGCTCAAGGCGAGCATGCCCGAC<br>GGCGAGGATCTCGTCGTGACCCATGGCGATGCCTGCTTGCCGAATATCATGGTGGAAAATGGCCGCTTTTCTGGATTCATCGACTGTGGCCGGCTGGGTGT<br>GGCGGACCGCTATCAGGACATAGCGTTGGCTACCCGTGATATTGCTGAAGAGCTTGGCGGCGAATGGGCTGACCGCTTCCTCGTGCTTTACGGTATCGCCG<br>CTCCCGATTCGCAGCGCATCGCCTTCTATCGCCTTCTTGACGAGTTCTTCTGAGCGGGACTCTGGGGTTCGAAATGACCGACCAAGCGACGCCCAACCTGC<br>CATCACGAGATTTCGATTCCACCGCCGCCTTCTATGAAAGGTTGGGCTTCGGAATCGTTTTCCGGGACGCCGGCTGGATGATCCTCCAGCGCGGGGATCTC<br>ATGCTGGAGTTCTTCGCCCACCCTAGGGGGAGGCTAACTGAAACACGGAAGGAGACAATACCGGAAGGAACCCGCGCTATGACGGCAATAAAAAGACAGAA<br>TAAAACGCACGGTGTTGGGTCGTTTGTTCATAAACGCGGGGTTCGGTCCCAGGGCTGGCACTCTGTCGATACCCCACCGAGACCCCATTGGGGCCAATACG<br>CCCGCGTTTCTTCCTTTTCCCCACCCCACCCCCCAAGTTCGGGTGAAGGCCCAGGGCTCGCAGCCAACGTCGGGGCGGCAGGCCCTGCCATAGCCTCAGGT<br>TACTCATATATACTTTAGATTGATTTAAAACTTCATTTTTAATTTAAAAGGATCTAGGTGAAGATCCTTTTTGATAATCTCATGACCAAAATCCCTTAACG<br>TGAGTTTTCGTTCCACTGAGCGTCAGACCCCGTAGAAAAGATCAAAGGATCTTCTTGAGATCCTTTTTTTCTGCGCGTAATCTGCTGCTTGCAAACAAAAA<br>AACCACCGCTACCAGCGGTGGTTTGTTTGCCGGATCAAGAGCTACCAACTCTTTTTCCGAAGGTAACTGGCTTCAGCAGAGCGCAGATACCAAATACTGTC<br>CTTCTAGTGTAGCCGTAGTTAGGCCACCACTTCAAGAACTCTGTAGCACCGCCTACATACCTCGCTCTGCTAATCCTGTTACCAGTGGCTGCTGCCAGTGG<br>CGATAAGTCGTGTCTTACCGGGTTGGACTCAAGACGATAGTTACCGGATAAGGCGCAGCGGTCGGGCTGAACGGGGGGTTCGTGCACACAGCCCAGCTTGG<br>AGCGAACGACCTACACCGAACTGAGATACCTACAGCGTGAGCTATGAGAAAGCGCCACGCTTCCCGAAGGGAGAAAGGCGGACAGGTATCCGGTAAGCGGC<br>AGGGTCGGAACAGGAGAGCGCACGAGGGAGCTTCCAGGGGGAAACGCCTGGTATCTTTATAGTCCTGTCGGGTTTCGCCACCTCTGACTTGAGCGTCGATT<br>TTTGTGATGCTCGTCAGGGGGGCGGAGCCTATGGAAAAACGCCAGCAACGCGGCCTTTTTACGGTTCCTGGCCTTTTGCTGGCCTTTTGCTCACATGTTCT<br>TTCCTGCGTTATCCCCTGATTCTGTGGATAACCGTATTACCGCCATGCAT |
| pBlockB | TAGTTATTAATAGTAATCAATTACGGGGTCATTAGTTCATAGCCCATATATGGAGTTCCGCGTTACATAACTTACGGTAAATGGCCCGCCTGGCTGACCGC<br>CCAACGACCCCCGCCCATTGACGTCAATAATGACGTATGTTCCCATAGTAACGCCAATAGGGACTTTCCATTGACGTCAATGGGTGGAGTATTTACGGTAA<br>ACTGCCCACTTGGCAGTACATCAAGTGTATCATATGCCAAGTACGCCCCCTATTGACGTCAATGACGGTAAATGGCCCGCCTGGCATTATGCCCAGTACAT<br>GACCTTATGGGACTTTCCTACTTGGCAGTACATCTACGTATTAGTCATCGCTATTACCATGGTGATGCGGTTTTGGCAGTACATCAATGGGCGTGGATAGC<br>GGTTTGACTCACGGGGATTTCCAAGTCTCCACCCCATTGACGTCAATGGGAGTTTGTTTTGGCACCAAAATCAACGGGACTTTCCAAAATGTCGTAACAAC<br>TCCGCCCCATTGACGCAAATGGGCGGTAGGCGTGTACGGTGGGAGGTCTATATAAGCAGAGCTGGTTTAGTGAACCGTCAGATCCGCTAGCGCTACCGGAC<br>TCAGATCTCGAGCTCAAGCTTCGAATTCTGCAGTCGACGGTACCGCGGGCCCGTCTTTATCCTTTTTCTTTTGATGCTTCATTCTCTCTTTTCTTTTTTTT<br>CCTCTGTAGCGTTTCTAGGAATCTCTCTCCAACGTTTAGGTTTCTTCTCAGATTTAGAAAGTTTCTTCTCTATCTGAAGAATTTTCATTTCGATTTCGCTT<br>CTTCCTTCACCCTAATCCGTAATCGAATGCCATTTCGAAGATTGTGTTGAAGAGTAGAGAGGAAGAAGAAGACTAAGAAGCTAGGGTTTCTCTGATGAGTA<br>GATGATGCGTGAACCGCTTGTTAGCGAAGAAGAAGAAGAAGAAGCTACTGAAGTTTTACTAGTGGAGAAGACGAAGTTGTGTAAGAGACGAGGAGACGAAG<br>AAAAAACGGAAGAGAGAAGAGACGATTTGTTGTTACTAGCTTTAACGCCGATGGTACGATCGAAATCTCAAGGAACTACTCGGCGCGTCACTCCCACGCCA<br>CCTCCTGTTGACGTGGAGAAACCGTTACCAAACGGAGATCTCTACATGGGAACGTTTTCCGGCGGGTTTCCCAACGGATCCGGTAAGTATTTGTGGAAGGA<br>TGGGTGTATGTACGAAGGCGAGTGGAAACGTGGTAAAGCGAGTGGTAAAGGCAAGTTTTCGTGGCCGAGTGGTGCGACTTATGAAGGAGAGTTCAAATCTG<br>GGAGAATGGAAGGATCTGGGACTTTTGTTGGTGTTGATGGTGATACTTATCGTGGCTCTTGGGTTGCTGATCGGAAACAAGGTCATGGTCAGAAGAGATAT<br>GCCAACGGAGATTACTATGAAGGTACATGGCGGCGGAATCTTCAGGATGGGAGAGGGAGATATGTTTGGATGAATGGGAATCAGTATACAGGAGAGTGGAG<br>AAATGGTGTGATATGTGGTAAAGGTGTGCTTGCTTGGCCTAATGGGAATAGATATGAAGGTCAATGGGAAAATGGTGTTCCTAAAGGAAGTGGTGTGTTTA<br>CTTGGGCTGATGGAAGTTCGTGGATTGGTTCTTGGAATGAGAGTAGTAATCTCATGAGGAATTTCTTTGATGGGATTGAGAAGAATGAGTTGATTGTTGCG<br>ACTAGGAAGAGATCTTCGGTTGATAGTGGCGCTGGAAGTTTGACTGGGGAGAAGATTTTCCCTAGGATATGTATTTGGGAGTCTGATGGAGAAGCTGGGGA<br>TATTACTTGTGATATTGTTGATAATGTGGAAGCTTCTGTGATATACAGAGATAGGATTTCTATTGATAAAGATGGGTTTCGTCAGTTTAGGAAGAATCCTT<br>GTTGTTTCAGCGGTGAGGCTAAGAAACCTGGAGAGACGATATCTAAAGGGCATAAGAAATATGATTTGATGCTCAACCTGCAGCATGGAATTAGGTAACTT<br>CTTAGAAGCTTGAGGATCCCAATGTTTGGAGAGTTTCTGGATCTGTGTTTTTCTAACCAAGTTGTGAATTTTTAGGTACTCTATTCATTTTTGTAAATATG<br>TACGTACACATAATGCTTTTATCTTCCATTACGTACACTAGATGTTATTGGTGCGCTCATTGAGGTTGGCAACTTGACCGAAGACTACCGTGGACGGAAGC<br>TTCCTTTTAAGATAATGGATAAACAGTAAGTTGTTTATTTTTAATTATTTAGGGTTTAGGGTTTGTTATATCCTGTGTATAATATATATTCTCTGTAGCAA<br>GGAAGTTCTTCAATGTGAAGCACAAAATGACCAAGCTTTCGAATTTCAAAGATTCTTCCAGCGGCTAACTGGTCCTCAAAACATTATCGCACTCGTATCCT<br>GGCGTCTAACCGGGATTCATAAACTGAAGGCTGTGACCAATGGCCCCATCTCCAAAGTCATTGCTAATCCGGATATACCGGAAGTGGAAGAAATCAGAGTC<br>GTCGTCTATTAGTTAATTATTGTTTCTTTTATTCTAAATAACTACAGATTTGTAACCGAATTTAAACATATCCCTACTTTTATGCTAAAGTTTTACCTTCT<br>AATTGCTACAGTTTACTTATTACAAATACAGCCTAATTGTAATGACTTTCCATGGACGTACAATTATGTTTTTGTACGTTCAAATAGGTGTACTAGATGTA<br>TAATGTAAACAAACTTAGTTCCATTCATAAGATAAACAATAAAACAAGGAAATAAAAAAACGTTAAAATAAGAGTAGGAAATACAAAGCATAACAACAAAC<br>CGAGTAGAGCATACAACCATGTTAATAAAGCCGAAGCATATAACAGGAAAACATATGCTTTACAAACAACACGAAATTATAGCCGATGCATAGAATGGGAA<br>ATCAGATGCTGAACGAACGCCTTTCCTTAATAAAATATGCACAACTCCCTAGCATACCAGTAGAAGAAGCAATTGCGGCAGCAGGCTCCGTAGCTGCAACG<br>ACCCAAGCAGTCTGGTGAAACCACGAACCTCCATGAACGATTCCACAAACGAGTTCCTTCAGCATGCAGTCGACTTATGTGATACATAACCAACTCCCCCA<br>CTGTGTTTGCAGCTTCGAATGATGGCACTAAATCCCAGAAAGTCGCGTTCACGGCTTCAACCTCTCGTCCGTTGCCAGTTGCGGTGAGGAACAGACCATGA<br>GAGAAACATGCGTAAGCGTCGATGGGCGAGATAGAACTCAACATGGTTAAACCATGAGCAACCGTAACGAAATTGCAAGCTAAACGAACCCCTTCAAGGTA<br>AATCGCCGGCCTTGCTCCACGTTCAAGCAGGAGGGTGAAATATGTTCGGAAAGGTGTTTCCGGCAACACGAAATCAGGCCACGTCATAAACTCATCCATGT<br>TGATAGCGGTCGGCTCCAGCGTGCTAACGAGACCTCTGAGTGTTTCAATGGGAGTTACGGAAACGGGTAAGTAGTTGACGAGGATGCTGAGTGGGTTAGGT<br>AGCTGAATGGCCAGTAAATTTCCGCCGGTGGGGGTGTGGGGACGTCCATTGCCGAGACTGAGGGATTTATTTGTGAGTTTAGGTTATATATGGCGTAGTAA<br>GAGCAATGAAAAGTGTGGAGAAGATGACTGTCGCCACCATGGTGAGCAAGGGCGAGGAGCTGTTCACCGGGGTGGTGCCCATCCTGGTCGAGCTGGACGGC<br>GACGTAAACGGCCACAAGTTCAGCGTGTCCGGCGAGGGCGAGGGCGATGCCACCTACGGCAAGCTGACCCTGAAGTTCATCTGCACCACCGGCAAGCTGCC<br>CGTGCCCTGGCCCACCCTCGTGACCACCCTGACCTACGGCGTGCAGTGCTTCAGCCGCTACCCCGACCACATGAAGCAGCACGACTTCTTCAAGTCCGCCA<br>TGCCCGAAGGCTACGTCCAGGAGCGCACCATCTTCTTCAAGGACGACGGCAACTACAAGACCCGCGCCGAGGTGAAGTTCGAGGGCGACACCCTGGTGAAC<br>CGCATCGAGCTGAAGGGCATCGACTTCAAGGAGGACGGCAACATCCTGGGGCACAAGCTGGAGTACAACTACAACAGCCACAACGTCTATATCATGGCCGA<br>CAAGCAGAAGAACGGCATCAAGGTGAACTTCAAGATCCGCCACAACATCGAGGACGGCAGCGTGCAGCTCGCCGACCACTACCAGCAGAACACCCCCATCG<br>GCGACGGCCCCGTGCTGCTGCCCGACAACCACTACCTGAGCACCCAGTCCGCCCTGAGCAAAGACCCCAACGAGAAGCGCGATCACATGGTCCTGCTGGAG<br>TTCGTGACCGCCGCCGGGATCACTCTCGGCATGGACGAGCTGTACAAGTAAAGCGGCCGCGACTCTAGATCATAATCAGCCATACCACATTTGTAGAGGTT<br>TTACTTGCTTTAAAAAACCTCCCACACCTCCCCCTGAACCTGAAACATAAAATGAATGCAATTGTTGTTGTTAACTTGTTTATTGCAGCTTATAATGGTTA<br>CAAATAAAGCAATAGCATCACAAATTTCACAAATAAAGCATTTTTTTCACTGCATTCTAGTTGTGGTTTGTCCAAACTCATCAATGTATCTTAAGGCGTAA<br>ATTGTAAGCGTTAATATTTTGTTAAAATTCGCGTTAAATTTTTGTTAAATCAGCTCATTTTTTAACCAATAGGCCGAAATCGGCAAAATCCCTTATAAATC<br>AAAAGAATAGACCGAGATAGGGTTGAGTGTTGTTCCAGTTTGGAACAAGAGTCCACTATTAAAGAACGTGGACTCCAACGTCAAAGGGCGAAAAACCGTCT<br>ATCAGGGCGATGGCCCACTACGTGAACCATCACCCTAATCAAGTTTTTTGGGGTCGAGGTGCCGTAAAGCACTAAATCGGAACCCTAAAGGGAGCCCCCGA<br>TTTAGAGCTTGACGGGGAAAGCCGGCGAACGTGGCGAGAAAGGAAGGGAAGAAAGCGAAAGGAGCGGGCGCTAGGGCGCTGGCAAGTGTAGCGGTCACGCT<br>GCGCGTAACCACCACACCCGCCGCGCTTAATGCGCCGCTACAGGGCGCGTCAGGTGGCACTTTTCGGGGAAATGTGCGCGGAACCCCTATTTGTTTATTTT |

| | |
|---|---|
| | TCTAAATACATTCAAATATGTATCCGCTCATGAGACAATAACCCTGATAAATGCTTCAATAATATTGAAAAAGGAAGAGTCCTGAGGCGGAAAGAACCAGC<br>TGTGGAATGTGTGTCAGTTAGGGTGTGGAAAGTCCCCAGGCTCCCCAGCAGGCAGAAGTATGCAAAGCATGCATCTCAATTAGTCAGCAACCAGGTGTGGA<br>AAGTCCCCAGGCTCCCCAGCAGGCAGAAGTATGCAAAGCATGCATCTCAATTAGTCAGCAACCATAGTCCCGCCCCTAACTCCGCCCATCCCGCCCCTAAC<br>TCCGCCCAGTTCCGCCCATTCTCCGCCCCATGGCTGACTAATTTTTTTTATTTATGCAGAGGCCGAGGCCGCCTCGGCCTCTGAGCTATTCCAGAAGTAGT<br>GAGGAGGCTTTTTTGGAGGCCTAGGCTTTTGCAAAGATCGATCAAGAGACAGGATGAGGATCGTTTCGCATGATTGAACAAGATGGATTGCACGCAGGTTC<br>TCCGGCCGCTTGGGTGGAGAGGCTATTCGGCTATGACTGGGCACAACAGACAATCGGCTGCTCTGATGCCGCCGTGTTCCGGCTGTCAGCGCAGGGGCGCC<br>CGGTTCTTTTTGTCAAGACCGACCTGTCCGGTGCCCTGAATGAACTGCAAGACGAGGCAGCGCGGCTATCGTGGCTGGCCACGACGGGCGTTCCTTGCGCA<br>GCTGTGCTCGACGTTGTCACTGAAGCGGGAAGGGACTGGCTGCTATTGGGCGAAGTGCCGGGGCAGGATCTCCTGTCATCTCACCTTGCTCCTGCCGAGAA<br>AGTATCCATCATGGCTGATGCAATGCGGCGGCTGCATACGCTTGATCCGGCTACCTGCCCATTCGACCACCAAGCGAAACATCGCATCGAGCGAGCACGTA<br>CTCGGATGGAAGCCGGTCTTGTCGATCAGGATGATCTGGACGAAGAGCATCAGGGGCTCGCGCCAGCCGAACTGTTCGCCAGGCTCAAGGCGAGCATGCCC<br>GACGGCGAGGATCTCGTCGTGACCCATGGCGATGCCTGCTTGCCGAATATCATGGTGGAAAATGGCCGCTTTTCTGGATTCATCGACTGTGGCCGGCTGGG<br>TGTGGCGGACCGCTATCAGGACATAGCGTTGGCTACCCGTGATATTGCTGAAGAGCTTGGCGGCGAATGGGCTGACCGCTTCCTCGTGCTTTACGGTATCG<br>CCGCTCCCGATTCGCAGCGCATCGCCTTCTATCGCCTTCTTGACGAGTTCTTCTGAGCGGGACTCTGGGGTTCGAAATGACCGACCAAGCGACGCCCAACC<br>TGCCATCACGAGATTTCGATTCCACCGCCGCCTTCTATGAAAGGTTGGGCTTCGGAATCGTTTTCCGGGACGCCGGCTGGATGATCCTCCAGCGCGGGGAT<br>CTCATGCTGGAGTTCTTCGCCCACCCTAGGGGGAGGCTAACTGAAACACGGAAGGAGACAATACCGGAAGGAACCCGCGCTATGACGGCAATAAAAAGACA<br>GAATAAAACGCACGGTGTTGGGTCGTTTGTTCATAAACGCGGGGTTCGGTCCCAGGGCTGGCACTCTGTCGATACCCCACCGAGACCCCATTGGGGCCAAT<br>ACGCCCGCGTTTCTTCCTTTTCCCCACCCCACCCCCCAAGTTCGGGTGAAGGCCCAGGGCTCGCAGCCAACGTCGGGGCGGCAGGCCCTGCCATAGCCTCA<br>GGTTACTCATATATACTTTAGATTGATTTAAAACTTCATTTTTAATTTAAAAGGATCTAGGTGAAGATCCTTTTTGATAATCTCATGACCAAAATCCCTTA<br>ACGTGAGTTTTCGTTCCACTGAGCGTCAGACCCCGTAGAAAAGATCAAAGGATCTTCTTGAGATCCTTTTTTTCTGCGCGTAATCTGCTGCTTGCAAACAA<br>AAAAACCACCGCTACCAGCGGTGGTTTGTTTGCCGGATCAAGAGCTACCAACTCTTTTTCCGAAGGTAACTGGCTTCAGCAGAGCGCAGATACCAAATACT<br>GTCCTTCTAGTGTAGCCGTAGTTAGGCCACCACTTCAAGAACTCTGTAGCACCGCCTACATACCTCGCTCTGCTAATCCTGTTACCAGTGGCTGCTGCCAG<br>TGGCGATAAGTCGTGTCTTACCGGGTTGGACTCAAGACGATAGTTACCGGATAAGGCGCAGCGGTCGGGCTGAACGGGGGGTTCGTGCACACAGCCCAGCT<br>TGGAGCGAACGACCTACACCGAACTGAGATACCTACAGCGTGAGCTATGAGAAAGCGCCACGCTTCCCGAAGGGAGAAAGGCGGACAGGTATCCGGTAAGC<br>GGCAGGGTCGGAACAGGAGAGCGCACGAGGGAGCTTCCAGGGGGAAACGCCTGGTATCTTTATAGTCCTGTCGGGTTTCGCCACCTCTGACTTGAGCGTCG<br>ATTTTTGTGATGCTCGTCAGGGGGGCGGAGCCTATGGAAAAACGCCAGCAACGCGGCCTTTTTACGGTTCCTGGCCTTTTGCTGGCCTTTTGCTCACATGT<br>TCTTTCCTGCGTTATCCCCTGATTCTGTGGATAACCGTATTACCGCCATGCAT |
| pBlockC | TAGTTATTAATAGTAATCAATTACGGGGTCATTAGTTCATAGCCCATATATGGAGTTCCGCGTTACATAACTTACGGTAAATGGCCCGCCTGGCTGACCGC<br>CCAACGACCCCCGCCCATTGACGTCAATAATGACGTATGTTCCCATAGTAACGCCAATAGGGACTTTCCATTGACGTCAATGGGTGGAGTATTTACGGTAA<br>ACTGCCCACTTGGCAGTACATCAAGTGTATCATATGCCAAGTACGCCCCCTATTGACGTCAATGACGGTAAATGGCCCGCCTGGCATTATGCCCAGTACAT<br>GACCTTATGGGACTTTCCTACTTGGCAGTACATCTACGTATTAGTCATCGCTATTACCATGGTGATGCGGTTTTGGCAGTACATCAATGGGCGTGGATAGC<br>GGTTTGACTCACGGGGATTTCCAAGTCTCCACCCCATTGACGTCAATGGGAGTTTGTTTTGGCACCAAAATCAACGGGACTTTCCAAAATGTCGTAACAAC<br>TCCGCCCCATTGACGCAAATGGGCGGTAGGCGTGTACGGTGGGAGGTCTATATAAGCAGAGCTGGTTTAGTGAACCGTCAGATCCGCTAGCGCTACCGGAC<br>TCAGATCTCGAGCTCAAGCTTCGAATTCTGCAGTCGACGGTACCGCGGGCCCGGATTATACCCACTGAAGGGTATTTCCGTCAATCCCAATTCTAACAATG<br>AATTCAGGAGTATAAAAACGTAAATTCAAGCGTGCCAATTATAAACCGTCGATCATAATCTAATCCAACGGCAGTAACATCGATCCGCGTGATTGTTTATT<br>ATTGGATAAGAATCACTCAACCGTCTCTACACAGTATATATAATAACCAAAGAGCGTCCTCTTACGCTTATCTTAATTTCCCTCGCATTGAGAATTTTCAA<br>CTTTTTCTATCTCTCTTCCCAAATCACAAATGGCGGGTCGCGGCAAAACTCTCGGATCTGGCGTTGCTAAGAAATCAACATCGAGAAGCAGCAAAGCCGGT<br>CTCCAATTCCCCGTTGGTCGTATCGCTCGTTTTCTAAAGAACGGCAAGTACGCAACACGTGTTGGTGCCGGAGCTCCGGTTTACTTAGCCGCCGTTCTCGA<br>ATACCTCGCCGCTGAGGTAATTATCCCCTTCTCTCCCTATATCTCTTTACTCTTTCGATCTTCAATTTCGTAAAACCCTAATTTCTAAATTGGATCTGTTG<br>TGTTGTAGGTATTGGAATTGGCTGGAAACGCAGCTAGGGATAACAAGAAGACTAGGATTGTGCCACGTCACATTCAGCTCGCGGTGAGAAACGATGAGGAG<br>CTGAGTAAACTGCTTGGAGATGTGACGATTGCTAATGGAGGTGTGATGCCTAACATTCACAGTCTTCTTCTTCCCAAGAAAGCTGGTGCTTCAAAACCTTC<br>CGCTGATGAAGATTAGATTAGGGATTTGTGTTGTGGTTGTTTAGCTAATTAATGTGTAGCTTAGTCTTTCATTAGATTAGATCTGAATTAGTTTTCATTAA<br>TGGTGTTGTGTAGTCTCTCTTTTGCTTCAAAAACAAGTATTAAAATCTTATTATTTTGAATTGAATCCACAATCAATACACATTGAAGTCCTAACAAACTA<br>CTTCTTCCCAGTGATATTTGAAACCAAATCACTAAGAAACTTAGCTGATTTGGTAATAGGAGAATTCATAGCCATCAAGTTATACAGAACAAGCTCAACTT<br>CTTCGATTGATGGTCGAGAATTGAATTGTGAAACAACTTTCAAAGTACCATTACCTTCTTCTTCTTCAACGAGAACATTCCATCTTTCTCCACTCACAACA<br>CCGTCTCTTACACACTTAAAGATCACTCCTTTTCTCTTACTAAACACTCCTCTCACTTCTAAGCCAGTGAACGCATAAATCACTTCAAAGGAACCAATAAA<br>GCCATCTCCAAAGCTTTTCTCTTCTTCAAACAACCATCTCGGATTGATCATTACTACAGGTTTTGTAGAAAAAGCTTCACTCGCGGTCTTAACATCCTCCA<br>GCTGCGATTTCTCCGGAGCCATGAAGATCGCCACGTCAGCAGATCTCAGAGCTGTTTTAAGAGAGTCTGAGACCGGAAGTCATCTTCTCCACACTTTTCAT<br>TGCTCTTACTACGCCATATATAACCTAAACTCACAAATAAATCCCTCAGTCTCGGCAATGGACGTCCCCACACCCCCACCGGCGGAAATTTACCTCGCCAT<br>TCAGCTACCTAACCCACTCAGCATCCTCGTCAACTACTTACCCGTTTCCGTAACTCCCATTGAAACACTCAGAGGTCTCGTTAGCACGCTGGAGCCGACCG<br>CTATCAACATGGATGAGTTTATGACGTGGCCTGATTTCGTGTTGCCGGAAACACCTTTCCGAACATATTTCACCCTCCTGCTTGAACGTGGAGCAAGGCCG<br>GCGATTTACCTTGAAGGGGTTCGTTTAGCTTGCAATTTCATTACGGTTGCTCATGGTTTAACCATGTTGAGTTCTATCTCGCCCATCGACGCTTACGCATG<br>TTTCTCTCATGGTCTGTTCCTCACCGCAACTGGCAACGGACGAGAGGTTGAAGCCGTGAACGCGACTTTCTGGGATTTAGTGCCATCATTCGAAGCTGCAA<br>ACACAGTGGGGGAGTTGGTTATGTATCACATAAGTCGACTGCATGCTGAAGGAACTCGTTTGTGGAATCGTTCATGGAGGTTCGTGGTTTCACCAGACTGC<br>TTGGGTCGTTGCAGCTACGGAGCCTGCTGCCGCAATTGCTTCTTCTACTGGTATGCTAGGGAGTTGTGCATATTTTATTAAGGAAAGGCGTTCGTTCAGCA<br>TCTGATTTCCCATTCTATGCATCGGCTATAATTTCGCGTTGTTTGTAAAGCATATGTTTTCCTGTTATATGCTTCGGCTTTATTAACATGGTTGTATGCTC<br>TACTCGGTTTGTTGTTATGCTTTGTATTTCCTACTCTTATTTTAACGTTTTTTTATTTCCTTGTTTTATTGTTTATCTTATGAATGGAACTAAGTTTGTTT<br>ACATTATACATATAGTACACCTATTTGAACGTACCAAAACATAATTGTACGTCCATGGAAAGTCATTACAATTAGGCTGTATTTGTAATAAGTAAACTGTA<br>GCAATTAGAAGGTAAAACTTTAGCATAAAAGTAGGGATATGTTTAAATTCGGTTACAAATCTGTAGTTATTTAGAATAAAAGAAACAATAATTAACTAATA<br>GACCACGACTCTGATTTCTTCCACTTCCGGTATATCCGGATTAGCAATGACTTTGGAGATGGGGCCATTGGTCACAGCCTTCAGTTTATGAATCCCGGTTA<br>GACGCCAGGATACGAGTGCGATAATGTTTTGAGGACCAGTTAGCCGCTGGAAGAATCTTTGAAATTCGAAAGCTTGGGCATTTTGTGCTTCACATTGAAGA<br>ACTTCCTTGCTACAGAGAATATATATTATACACAGGATATAACAAACCTAAACCCTAAATAATTAAAAATAAACAACTTATTGTTTATCCATTATCTTAAA<br>AGGAAGCTTCCGTCCACGGTAGTCTTCGGTCAAGTTGCCAACCTCAATGAGCACACCAATAACATCTAGTGTACGTAATGGAAGATAAAAGCATTATGTGT<br>ACGTACATATTTACAAAAATGGTCGCCACCATGGTGAGCAAGGGCGAGGAGCTGTTCACCGGGGTGGTGCCCATCCTGGTCGAGCTGGACGGCGACGTAAA<br>CGGCCACAAGTTCAGCGTGTCCGGCGAGGGCGAGGGCGATGCCACCTACGGCAAGCTGACCCTGAAGTTCATCTGCACCACCGGCAAGCTGCCCGTGCCCT<br>GGCCCACCCTCGTGACCACCCTGACCTACGGCGTGCAGTGCTTCAGCCGCTACCCCGACCACATGAAGCAGCACGACTTCTTCAAGTCCGCCATGCCCGAA<br>GGCTACGTCCAGGAGCGCACCATCTTCTTCAAGGACGACGGCAACTACAAGACCCGCGCCGAGGTGAAGTTCGAGGGCGACACCCTGGTGAACCGCATCGA<br>GCTGAAGGGCATCGACTTCAAGGAGGACGGCAACATCCTGGGGCACAAGCTGGAGTACAACTACAACAGCCACAACGTCTATATCATGGCCGACAAGCAGA<br>AGAACGGCATCAAGGTGAACTTCAAGATCCGCCACAACATCGAGGACGGCAGCGTGCAGCTCGCCGACCACTACCAGCAGAACACCCCCATCGGCGACGGC<br>CCCGTGCTGCTGCCCGACAACCACTACCTGAGCACCCAGTCCGCCCTGAGCAAAGACCCCAACGAGAAGCGCGATCACATGGTCCTGCTGGAGTTCGTGAC<br>CGCCGCCGGGATCACTCTCGGCATGGACGAGCTGTACAAGTAAAGCGGCCGCGACTCTAGATCATAATCAGCCATACCACATTTGTAGAGGTTTTACTTGC<br>TTTAAAAAACCTCCCACACCTCCCCCTGAACCTGAAACATAAAATGAATGCAATTGTTGTTGTTAACTTGTTTATTGCAGCTTATAATGGTTACAAATAAA<br>GCAATAGCATCACAAATTTCACAAATAAAGCATTTTTTTCACTGCATTCTAGTTGTGGTTTGTCCAAACTCATCAATGTATCTTAAGGCGTAAATTGTAAG<br>CGTTAATATTTTGTTAAAATTCGCGTTAAATTTTTGTTAAATCAGCTCATTTTTTAACCAATAGGCCGAAATCGGCAAAATCCCTTATAAATCAAAAGAAT<br>AGACCGAGATAGGGTTGAGTGTTGTTCCAGTTTGGAACAAGAGTCCACTATTAAAGAACGTGGACTCCAACGTCAAAGGGCGAAAAACCGTCTATCAGGGC<br>GATGGCCCACTACGTGAACCATCACCCTAATCAAGTTTTTTGGGGTCGAGGTGCCGTAAAGCACTAAATCGGAACCCTAAAGGGAGCCCCCGATTTAGAGC<br>TTGACGGGGAAAGCCGGCGAACGTGGCGAGAAAGGAAGGGAAGAAAGCGAAAGGAGCGGGCGCTAGGGCGCTGGCAAGTGTAGCGGTCACGCTGCGCGTAA<br>CCACCACACCCGCCGCGCTTAATGCGCCGCTACAGGGCGCGTCAGGTGGCACTTTTCGGGGAAATGTGCGCGGAACCCCTATTTGTTTATTTTTCTAAATA<br>CATTCAAATATGTATCCGCTCATGAGACAATAACCCTGATAAATGCTTCAATAATATTGAAAAAGGAAGAGTCCTGAGGCGGAAAGAACCAGCTGTGGAAT |

GTGTGTCAGTTAGGGTGTGGAAAGTCCCCAGGCTCCCCAGCAGGCAGAAGTATGCAAAGCATGCATCTCAATTAGTCAGCAACCAGGTGTGGAAAGTCCCC
AGGCTCCCCAGCAGGCAGAAGTATGCAAAGCATGCATCTCAATTAGTCAGCAACCATAGTCCCGCCCCTAACTCCGCCCATCCCGCCCCTAACTCCGCCCA
GTTCCGCCCATTCTCCGCCCCATGGCTGACTAATTTTTTTTATTTATGCAGAGGCCGAGGCCGCCTCGGCCTCTGAGCTATTCCAGAAGTAGTGAGGAGGC
TTTTTTGGAGGCCTAGGCTTTTGCAAAGATCGATCAAGAGACAGGATGAGGATCGTTTCGCATGATTGAACAAGATGGATTGCACGCAGGTTCTCCGGCCG
CTTGGGTGGAGAGGCTATTCGGCTATGACTGGGCACAACAGACAATCGGCTGCTCTGATGCCGCCGTGTTCCGGCTGTCAGCGCAGGGGCGCCCGGTTCTT
TTTGTCAAGACCGACCTGTCCGGTGCCCTGAATGAACTGCAAGACGAGGCAGCGCGGCTATCGTGGCTGGCCACGACGGGCGTTCCTTGCGCAGCTGTGCT
CGACGTTGTCACTGAAGCGGGAAGGGACTGGCTGCTATTGGGCGAAGTGCCGGGGCAGGATCTCCTGTCATCTCACCTTGCTCCTGCCGAGAAAGTATCCA
TCATGGCTGATGCAATGCGGCGGCTGCATACGCTTGATCCGGCTACCTGCCCATTCGACCACCAAGCGAAACATCGCATCGAGCGAGCACGTACTCGGATG
GAAGCCGGTCTTGTCGATCAGGATGATCTGGACGAAGAGCATCAGGGGCTCGCGCCAGCCGAACTGTTCGCCAGGCTCAAGGCGAGCATGCCCGACGGCGA
GGATCTCGTCGTGACCCATGGCGATGCCTGCTTGCCGAATATCATGGTGGAAAATGGCCGCTTTTCTGGATTCATCGACTGTGGCCGGCTGGGTGTGGCGG
ACCGCTATCAGGACATAGCGTTGGCTACCCGTGATATTGCTGAAGAGCTTGGCGGCGAATGGGCTGACCGCTTCCTCGTGCTTTACGGTATCGCCGCTCCC
GATTCGCAGCGCATCGCCTTCTATCGCCTTCTTGACGAGTTCTTCTGAGCGGGACTCTGGGGTTCGAAATGACCGACCAAGCGACGCCCAACCTGCCATCA
CGAGATTTCGATTCCACCGCCGCCTTCTATGAAAGGTTGGGCTTCGGAATCGTTTTCCGGGACGCCGGCTGGATGATCCTCCAGCGCGGGGATCTCATGCT
GGAGTTCTTCGCCCACCCTAGGGGGAGGCTAACTGAAACACGGAAGGAGACAATACCGGAAGGAACCCGCGCTATGACGGCAATAAAAAGACAGAATAAAA
CGCACGGTGTTGGGTCGTTTGTTCATAAACGCGGGGTTCGGTCCCAGGGCTGGCACTCTGTCGATACCCCACCGAGACCCCATTGGGGCCAATACGCCCGC
GTTTCTTCCTTTTCCCCACCCCACCCCCCAAGTTCGGGTGAAGGCCCAGGGCTCGCAGCCAACGTCGGGGCGGCAGGCCCTGCCATAGCCTCAGGTTACTC
ATATATACTTTAGATTGATTTAAAACTTCATTTTTAATTTAAAAGGATCTAGGTGAAGATCCTTTTTGATAATCTCATGACCAAAATCCCTTAACGTGAGT
TTTCGTTCCACTGAGCGTCAGACCCCGTAGAAAAGATCAAAGGATCTTCTTGAGATCCTTTTTTTCTGCGCGTAATCTGCTGCTTGCAAACAAAAAAACCA
CCGCTACCAGCGGTGGTTTGTTTGCCGGATCAAGAGCTACCAACTCTTTTTCCGAAGGTAACTGGCTTCAGCAGAGCGCAGATACCAAATACTGTCCTTCT
AGTGTAGCCGTAGTTAGGCCACCACTTCAAGAACTCTGTAGCACCGCCTACATACCTCGCTCTGCTAATCCTGTTACCAGTGGCTGCTGCCAGTGGCGATA
AGTCGTGTCTTACCGGGTTGGACTCAAGACGATAGTTACCGGATAAGGCGCAGCGGTCGGGCTGAACGGGGGGTTCGTGCACACAGCCCAGCTTGGAGCGA
ACGACCTACACCGAACTGAGATACCTACAGCGTGAGCTATGAGAAAGCGCCACGCTTCCCGAAGGGAGAAAGGCGGACAGGTATCCGGTAAGCGGCAGGGT
CGGAACAGGAGAGCGCACGAGGGAGCTTCCAGGGGGAAACGCCTGGTATCTTTATAGTCCTGTCGGGTTTCGCCACCTCTGACTTGAGCGTCGATTTTTGT
GATGCTCGTCAGGGGGGCGGAGCCTATGGAAAAACGCCAGCAACGCGGCCTTTTTACGGTTCCTGGCCTTTTGCTGGCCTTTTGCTCACATGTTCTTTCCT
GCGTTATCCCCTGATTCTGTGGATAACCGTATTACCGCCATGCAT

p3IR7kb12

GCTAGGGTGAAGAGCACATGTGTTTTTCCATAGGCTCCGCCCCCCTGACGAGCATCACAAAAATCGACGCTCAAGTCAGAGGTGGCGAAACCCGACAGGAC
TATAAAGATACCAGGCGTTTCCCCCTGGAAGCTCCCTCGTGCGCTCTCCTGTTCCGACCCTGCCGCTTACCGGATACCTGTCCGCCTTTCTCCCTTCGGGA
AGCGTGGCGCTTTCTCATAGCTCACGCTGTAGGTATCTCAGTTCGGTGTAGGTCGTTCGCTCCAAGCTGGGCTGTGTGCACGAACCCCCCGTTCAGCCCGA
CCGCTGCGCCTTATCCGGTAACTATCGTCTTGAGTCCAACCCGGTAAGACACGACTTATCGCCACTGGCAGCAGCCACTGGTAACAGGATTAGCAGAGCGA
GGTATGTAGGCGGTGCTACAGAGTTCTTGAAGTGGTGGCCTAACTACGGCTACACTAGAAGAACAGTATTTGGTATCTGCGCTCTGCTGAAGCCAGTTACC
TTCGGAAAAAGAGTTGGTAGCTCTTGATCCGGCAAACAAACCACCGCTGGTAGCGGTGGTTTTTTTGTTTGCAAGCAGCAGATTACGCGCAGAAAAAAAGG
ATCTCAAGAAGATCCTTTGATCTTTTCTACGGGGTCTGACGCTCAGTGGAACGAAAACTCACGTTAAGGGATTTTGGTCATGAGATTATCAAAAAGGATCT
TCACCTAGATCCTTTTAAATTAAAAATGAAGTTTTAAATCAATCTAAAGTATATATGAGTAAACTTGGTCTGACAGTTACCAATGCTTAATCAGTGAGGCA
CCTATCTCAGCGATCTGTCTATTTCGTTCATCCATAGTTGCCTGACTCCCCGTCGTGTAGATAACTACGATACGGGAGGGCTTACCATCTGGCCCCAGTGC
TGCAATGATACCGCGAGACCCACGCTCACCGGCTCCAGATTTATCAGCAATAAACCAGCCAGCCGGAAGGGCCGAGCGCAGAAGTGGTCCTGCAACTTTAT
CCGCCTCCATCCAGTCTATTAATTGTTGCCGGGAAGCTAGAGTAAGTAGTTCGCCAGTTAATAGTTTGCGCAACGTTGTTGCCATTGCTACAGGCATCGTG
GTGTCACGCTCGTCGTTTGGTATGGCTTCATTCAGCTCCGGTTCCCAACGATCAAGGCGAGTTACATGATCCCCCATGTTGTGCAAAAAAGCGGTTAGCTC
CTTCGGTCCTCCGATCGTTGTCAGAAGTAAGTTGGCCGCAGTGTTATCACTCATGGTTATGGCAGCACTGCATAATTCTCTTACTGTCATGCCATCCGTAA
GATGCTTTTCTGTGACTGGTGAGTACTCAACCAAGTCATTCTGAGAATAGTGTATGCGGCGACCGAGTTGCTCTTGCCCGGCGTCAATACGGGATAATACC
GCGCCACATAGCAGAACTTTAAAAGTGCTCATCATTGGAAAACGTTCTTCGGGGCGAAAACTCTCAAGGATCTTACCGCTGTTGAGATCCAGTTCGATGTA
ACCCACTCGTGCACCCAACTGATCTTCAGCATCTTTTACTTTCACCAGCGTTTCTGGGTGAGCAAAAACAGGAAGGCAAAATGCCGCAAAAAAGGGAATAA
GGGCGACACGGAAATGTTGAATACTCATACTCTTCCTTTTTCAATATTATTGAAGCATTTATCAGGGTTATTGTCTCATGAGCGGATACATATTTGAATGT
ATTTAGAAAAATAAACAAATAGGGGTTCCGCGCACATTTCCCCGAAAAGTGCCACCTGACGTCTAAGAAACCATTATTATCACCCTTTTCATGATGACCCT
TTTCAACTCTATCCTTTTGTTTTATCCATTTTGTTTCTACCCTCTTTAAATTTTTAATGACCATTTTACCCTTATTTGGAAATTGTTTTAGATTAACAAAA
TGAAATATTAATATTTTTCCGCCTAAAATATTCCAAAATAAATTTCCCGCCAAAAATTTTCAAAAAAAAAATTCCGCCAAAAATTTTCAAAAAACAAATTC
CCGCCAAATTTTGTTTTCAAAAATATTTTTCCCGCCAATTTTTTTTTTGGCGGGAAATAGATTTACAAGGGATTTTTAAAGAGTTTTAAAAGATTTACAAG
AGATTTTAAAGGGGTTTTATAAGATTTACAAGAGATTTTAAAAGGGTGATTTACAAGGGATTTTAAAAGGTTTTTAAAATATTTACAAGAGATTTTAAAAG
GGTTTTAAAAGATTTACAAGAGATTTTAAAGAGATTTAAAAGATTTACAAAGGTCTTTTAAAGATTTTTAAACGGTTTAAAATTATTTTAAAATGATTTTA
AAATATTTACAAGGGATTTTAAAAGGGTTTTACAATATTTACAACAGATTTTAAAGGACTTTAAAATATTTACAAAAGATCTGTAAAGATTCTTAAAGGGT
TTAAAAGTGTTTTGAAAGATTTACAAGAGATTTTAAAACCTTTTAAAATTACTTGTAAATTTTTTAAAACCCTTGTAAATCTTTTAAAATCTCTTTTAAAT
CTTTTAAAACCTTTTTAAAATCTCTTGTAAATCTTTTAAGACCCTTTTAAAATCCCTTGTAAATCTATTTGCCGCAAACAAATATTTACAAGGATTTTTAA
AATTATTTTTTTGGCTCAGAAAAAAATTTGTTACGGAACAAAAAATTTGGCGGGAAACTTTTTTTTTGGCGGGAAAAAAGTTCGGCGGGAAATCTTTTTTGG
CGGGAAAAAAAAAATTTTGGCGCCAATAAATTTGGCGGGAAAAGTTCAGTTTCCAGCTGTCGGTACGCATCCGACAATCTCTCTATTCGCTATTCCCAGATC
TCCATGACCCACTTCACTCCCTGCGCGTCCGGTACAATCGCCACTTCCATAACACCAAGAGTCAATACAAAAAAAAAGCCGACTCCATTATTATCCCTCTC
TCTACAAAAAAAACAGTAGAGAGAGATATAGAGAGCATCAATGGCGTCGTCGTCAAGACTTTGCGATTCGTGCAAATCGACGGCAGCAACTCTATTCTGCC
GCGCAGACGCCGCGTTTCTCTGCGGCGACTGCGACGGTAAGATCCACACAGCTAACAAACTCGCATCACGTCACGAACGAGTTTGGCTATGTGAAGTATGC
GAACAAGCACCAGCACACGTCACGTGCAAAGCTGACGCCGCTGCGTTATGCGTCACGTGCGACCGTGATATCCACTCAGCTAATCCACTCTCTCGCCGCCA
CGAGCGCGTACCAATCACTCCTTTCTACGACGCTGTAGGTCCAGCAAAATCCGCTTCCTCCTCCGTCAATTTCGTAGATGAAGATGGCGGTGACGTCACTG
CTTCGTGGCTTTTAGCTAAAGAAGGAATCGAAATCACTAATTTGTTCTCCGATCTTGATTATCCGAAGATTGAGGTAACGTCGGAGGAGAATAGCTCCGGT
AACGACGGAGTTGTTCCTGTGCAGAATAAGTTGTTTCTTAACGAAGATTACTTCAATTTCGATCTCTCTGCTTCCAAAATTTCTCAACAAGGATTCAATTT
CATCAACCAAACTGTACGAATCTCTCTCTATCTTCTTAAGCTTCCATGGATCTGGAAAATTTTCAGATCTAGATCTGAAAGTTTGGATCTGATTTGAGTTT
GTGAAATGTTGTTAGGTTTCGACGAGAACGATAGATGTACCGTTGGTGCCTGAAAGTGGAGGAGTGACGGCGGAGATGACGAACACGGAGACGCCAGCTGT
GCAGTTATCACCGGCGGAGAGGGAAGCAAGGGTTTTGAGGTATAGAGAGAAGAGGAAGAATCGGAAATTTGAGAAGACGATTAGGTATGCGTCGCGTAAAG
CTTACGCTGAGATGAGGCCGAGAATCAAAGGACGTTTTGCTAAGAGAACAGATTCGAGAGAGAATGATGGTGGAGACGTCGGAGTTTATGGCGGATTCGGT
GTTGTTCCGAGTTTCTGATATTTTCCGGTGAAAGAAACATGGTAGTAGTATATCGGTTAATAGCAGATTAAGGATAATTAAAACTAAAATGTTTCCTGATT
TAATCAGGGGCTAAGTTATTGTTAGAAGATTGTTGTTTTTTGAATTAATTCTCACAATTGTATCCTAAATTATATAGTTTACTTAATGTTGAACTAATAAA
TTTTGATTTACAATCTCTGTAAGAAATCGATATGCAAAAATAAATGAGTTGAGCTATTAAGATTAACATTCATATAGAACACAAACTCATACAATCAGCTG
TGGTGGGAAAAAAAGCATTATAATCCTTATATTACTAGTGAGTAGTGACTGATACAAAAGATTTTTTATACGTTAAAAGTTTAAAAATTCAAAAGTACAGA
AAACGTTATATTCGCTTGTTACCTTTTTTAACTTTATCCACGTCATTACCCCTGAGAGTAGCCCAACGAAAAATGGAAAATTTAACCCAAAAAACGGAAAA
TATTAGAGAAAGCGTTAATGCCTTTTCTAACAACTAAACGACGACGTATCAGTTCAATCCTCTCTCTATTATTATTCCTTAAAACTTAACTCTAAAGCTTT
CGAGTATTCTTCAATCTCTTCTTCTCCGATCGATCTGAAGTAAGAGAGCTTCTTAAGGAAGAAGAAGATGGGTTGTGCTCAATCAAAGATCGAGAACGAAG
AAGCAGTTACTCGTTGCAAAGAACGAAAACAATTGATGAAAGACGCCGTCACTGCTCGTAACGCTTTCGCCGCCGCTCACTCAGCTTACGCTATGGCTCTT
AAAAACACCGGAGCTGCTCTTTCCGATTACTCTCACGGCGAGTTTTTAGTCTCTAATCACTCGTCTTCCTCCGCAGCTGCAGCAATCGCTTCTACTTCTTC
TCTTCCCACTGCTATATCTCCTCCTCTTCCTTCTTCCACCGCTCCGGTTTCTAATTCAACCGCTTCTTCTTCCTCCGCTGCGGTTCCTCAGCCGATTCCTG
ATACTCTTCCTCCTCCTCCTCCTCCACCACCGCTTCCTCTTCAACGTGCTGCTACTATGCCGGAGATGAACGGTAGATCCGGTGGTGGTCATGCTGGTAGT
GGACTCAACGGAATTGAAGAAGATGGAGCCCTAGATAACGATGATGATGACGATGATGATGATGATGACTCTGAAATGGAGAATCGTGATCGTTTGATTAG
GAAATCGAGAAGCCGTGGAGGTAGTACTAGAGGAAATAGGACGACGATTGAAGATCATCATCTTCAGGAGGAGAAAGCTCCGCCACCTCCCCCTTTGGCGA
ATTCACGGCCAATTCCGCCGCCACGTCAGCATCAGCATCAACATCAGCAACAGCAACAACAACCTTTCTACGATTACTTCTTCCCTAATGTTGAGAATATG

| | |
|---|---|
| | CCTGGAACTACTTTAGAAGATACTCCTCCACAACCACAACCACAACCAACAAGGCCTGTGCCTCCTCAACCACATTCACCAGTCGTTACTGAGGATGACGA |
| | AGATGAGGAGGAGGAAGAGGAGGAAGAGGAGGAGGAAGAGGAGACGGTGATTGAACGGAAACCACTGGTGGAGGAAAGACCGAAGAGAGTAGAGGAAGTGA |
| | CGATTGAATTGGAAAAAGTTACTAATTTGAGAGGGATGAAGAAGAGTAAAGGGATAGGGATTCCCGGAGAGAGGAGAGGAATGCGAATGCCGGTGACTGCG |
| | ACGCATTTGGCGAATGTATTCATTGAGCTTGATGATAATTTCTTGAAAGCTTCTGAAAGTGCTCATGATGTTTCTAAGATGCTTGAAGCTACTAGGCTCCA |
| | TTACCATTCTAATTTTGCAGATAACCGAGGTAATTTTGATTCGATAAATGTCGGACGTGCACCGACATCTTTATCCTTTTTCTTTTGATGCTTCATTCTCT |
| | CTTTTCTTTTTTTTCCTCTGTAGCGTTTCTAGGAATCTCTCTCCAACGTTTAGGTTTCTTCTCAGATTTAGAAAGTTTCTTCTCTATCTGAAGAATTTTCA |
| | TTTCGATTTCGCTTCTTCCTTCACCCTAATCCGTAATCGAATGCCATTTCGAAGATTGTGTTGAAGAGTAGAGAGGAAGAAGAAGACTAAGAAGCTAGGGT |
| | TTCTCTGATGAGTAGATGATGCGTGAACCGCTTGTTAGCGAAGAAGAAGAAGAAGCTACTGAAGTTTTACTAGTGGAGAAGACGAAGTTGTGTAAGAG |
| | ACGAGGAGACGAAGAAAAAACGGAAGAGAGAAGAGACGATTTGTTGTTACTAGCTTTAACGCCGATGGTACGATCGAAATCTCAAGGAACTACTCGGCGCG |
| | TCACTCCCACGCCACCTCCTGTTGACGTGGAGAAACCGTTACCAAACGGAGATCTCTACATGGGAACGTTTTCCGGCGGGTTTCCCAACGGATCCGGTAAG |
| | TATTTGTGGAAGGATGGGTGTATGTACGAAGGCGAGTGGAAACGTGGTAAAGCGAGTGGTAAAGGCAAGTTTTCGTGGCCGAGTGGTGCGACTTATGAAGG |
| | AGAGTTCAAATCTGGGAGAATGGAAGGATCTGGGACTTTTGTTGGTGTTGATGGTGATACTTATCGTGGCTCTTGGGTTGCTGATCGGAAACAAGGTCATG |
| | GTCAGAAGAGATATGCCAACGGAGATTACTATGAAGGTACATGGCGGCGGAATCTTCAGGATGGGAGAGGGAGATATGTTTGGATGAATGGGAATCAGTAT |
| | ACAGGAGAGTGGAGAAATGGTGTGATATGTGGTAAAGGTGTGCTTGCTTGGCCTAATGGGAATAGATATGAAGGTCAATGGGAAAATGGTGTTCCTAAAGG |
| | AAGTGGTGTGTTTACTTGGGCTGATGGAAGTTCGTGGATTGGTTCTTGGAATGAGAGTAGTAATCTCATGAGGAATTTCTTTGATGGGATTGAGAAGAATG |
| | AGTTGATTGTTGCGACTAGGAAGAGATCTTCGGTTGATAGTGGCGCTGGAAGTTTGACTGGGGAGAAGATTTTCCCTAGGATATGTATTTGGGAGTCTGAT |
| | GGAGAAGCTGGGGATATTACTTGTGATATTGTTGATAATGTGGAAGCTTCTGTGATATACAGAGATAGGATTTCTATTGATAAAGATGGGTTTCGTCAGTT |
| | TAGGAAGAATCCTTGTTGTTTCAGCGGTGAGGCTAAGAAACCTGGAGAGACGATATCTAAAGGGCATAAGAAATATGATTTGATGCTCAACCTGCAGCATG |
| | GAATTAGGTAACTTCTTAGAAGCTTGAGGATCCCAATGTTTGGAGAGTTTCTGGATCTGTGTTTTTCTAACCAAGTTGTGAATTTTTAGGTACTCTATTCA |
| | TTTTTGTAAATATGTACGTACACATAATGCTTTTATCTTCCATTACGTACACTAGATGTTATTGGTGCGCTCATTGAGGTTGGCAACTTGACCGAAGACTA |
| | CCGTGGACGGAAGCTTCCTTTTAAGATAATGGATAAACAGTAAGTTGTTTATTTTTAATTATTTAGGGTTTAGGGTTTGTTATATCCTGTGTATAATATAT |
| | ATTCTCTGTAGCAAGGAAGTTCTTCAATGTGAAGCACAAAATGACCAAGCTTTCGAATTTCAAAGATTCTTCCAGCGGCTAACTGGTCCTCAAAACATTAT |
| | CGCACTCGTATCCTGGCGTCTAACCGGGATTCATAAACTGAAGGCTGTGACCAATGGCCCCATCTCCAAAGTCATTGCTAATCCGGATATACCGGAAGTGG |
| | AAGAAATCAGAGTCGTCGTCTATTAGTTAATTATTGTTTCTTTTATTCTAAATAACTACAGATTTGTAACCGAATTTAAACATATCCCTACTTTTATGCTA |
| | AAGTTTTACCTTCTAATTGCTACAGTTTACTTATTACAAATACAGCCTAATTGTAATGACTTTCCATGGACGTACAATTATGTTTTTGTACGTTCAAATAG |
| | GTGTACTAGATGTATAATGTAAACAAACTTAGTTCCATTCATAAGATAAACAATAAAACAAGGAAATAAAAAAACGTTAAAATAAGAGTAGGAAATACAAA |
| | GCATAACAACAAACCGAGTAGAGCATACAACCATGTTAATAAAGCCGAAGCATATAACAGGAAAACATATGCTTTACAAACAACACGAAATTATAGCCGAT |
| | GCATAGAATGGGAAATCAGATGCTGAACGAACGCCTTTCCTTAATAAAATATGCACAACTCCCTAGCATACCAGTAGAAGAAGCAATTGCGGCAGCAGGCT |
| | CCGTAGCTGCAACGACCCAAGCAGTCTGGTGAAACCACGAACCTCCATGAACGATTCCACAAACGAGTTCCTTCAGCATGCAGTCGACTTATGTGATACAT |
| | AACCAACTCCCCCACTGTGTTTGCAGCTTCGAATGATGGCACTAAATCCCAGAAAGTCGCGTTCACGGCTTCAACCTCTCGTCCGTTGCCAGTTGCGGTGA |
| | GGAACAGACCATGAGAGAAACATGCGTAAGCGTCGATGGGCGAGATAGAACTCAACATGGTTAAACCATGAGCAACCGTAACGAAATTGCAAGCTAAACGA |
| | ACCCCTTCAAGGTAAATCGCCGGCCTTGCTCCACGTTCAAGCAGGAGGGTGAAATATGTTCGGAAAGGTGTTTCCGGCAACACGAAATCAGGCCACGTCAT |
| | AAACTCATCCATGTTGATAGCGGTCGGCTCCAGCGTGCTAACGAGACCTCTGAGTGTTTCAATGGGAGTTACGGAAACGGGTAAGTAGTTGACGAGGATGC |
| | TGAGTGGGTTAGGTAGCTGAATGGCCAGTAAATTTCCGCCGGTGGGGGTGTGGGGACGTCCATTGCCGAGACTGAGGGATTTATTTGTGAGTTTAGGTTAT |
| | ATATGGCGTAGTAAGAGCAATGAAAAGTGTGGAGAAGATGACTTGTCGGCAGTGACCCGACAGATTATACCCACTGAAGGGTATTTCCGTCAATCCCAATT |
| | CTAACAATGAATTCAGGAGTATAAAAACGTAAATTCAAGCGTGCCAATTATAAACCGTCGATCATAATCTAATCCAACGGCAGTAACATCGATCCGCGTGA |
| | TTGTTTATTATTGGATAAGAATCACTCAACCGTCTCTACACAGTATATATAATAACCAAAGAGCGTCCTCTTACGCTTATCTTAATTTCCCTCGCATTGAG |
| | AATTTTCAACTTTTTCTATCTCTCTTCCCAAATCACAAATGGCGGGTCGCGGCAAAACTCTCGGATCTGGCGTTGCTAAGAAATCAACATCGAGAAGCAGC |
| | AAAGCCGGTCTCCAATTCCCCGTTGGTCGTATCGCTCGTTTTCTAAAGAACGGCAAGTACGCAACACGTGTTGGTGCCGGAGCTCCGGTTTACTTAGCCGC |
| | CGTTCTCGAATACCTCGCCGCTGAGGTAATTATCCCCTTCTCTCCCTATATCTCTTTACTCTTTCGATCTTCAATTTCGTAAAACCCTAATTTCTAAATTG |
| | GATCTGTTGTGTTGTAGGTATTGGAATTGGCTGGAAACGCAGCTAGGGATAACAAGAAGACTAGGATTGTGCCACGTCACATTCAGCTCGCGGTGAGAAAC |
| | GATGAGGAGCTGAGTAAACTGCTTGGAGATGTGACGATTGCTAATGGAGGTGTGATGCCTAACATTCACAGTCTTCTTCTTCCCAAGAAAGCTGGTGCTTC |
| | AAAACCTTCCGCTGATGAAGATTAGATTAGGGATTTGTGTTGTGGTTGTTTAGCTAATTAATGTGTAGCTTAGTCTTTCATTAGATTAGATCTGAATTAGT |
| | TTTCATTAATGGTGTTGTGTAGTCTCTCTTTTGCTTCAAAAACAAGTATTAAAATCTTATTATTTTGAATTGAATCCACAATCAATACACATTGAAGTCCT |
| | AACAAACTACTTCTTCCCAGTGATATTTGAAACCAAATCACTAAGAAACTTAGCTGATTTGGTAATAGGAGAATTCATAGCCATCAAGTTATACAGAACAA |
| | GCTCAACTTCTTCGATTGATGGTCGAGAATTGAATTGTGAAACAACTTTCAAAGTACCATTACCTTCTTCTTCTTCAACGAGAACATTCCATCTTTCTCCA |
| | CTCACAACACCGTCTCTTACACACTTAAAGATCACTCCTTTTCTCTTACTAAACACTCCTCTCACTTCTAAGCCAGTGAACGCATAAATCACTTCAAAGGA |
| | ACCAATAAAGCCATCTCCAAAGCTTTTCTCTTCTTCAAACAACCATCTCGGATTGATCATTACTACAGGTTTTGTAGAAAAAGCTTCACTCGCGGTCTTAA |
| | CATCCTCCAGCTGCGATTTCTCCGGAGCCATGAAGATCGCCACGTCAGCAGATCTCAGAGCTGTTTTAAGAGAGTCTGAGACCGGAAGTCATCTTCTCCAC |
| | ACTTTTCATTGCTCTTACTACGCCATATATAACCTAAACTCACAAATAAATCCCTCAGTCTCGGCAATGGACGTCCCCACACCCCCACCGGCGGAAATTTA |
| | CCTCGCCATTCAGCTACCTAACCCACTCAGCATCCTCGTCAACTACTTACCCGTTTCCGTAACTCCCATTGAAACACTCAGAGGTCTCGTTAGCACGCTGG |
| | AGCCGACCGCTATCAACATGGATGAGTTTATGACGTGGCCTGATTTCGTGTTGCCGGAAACACCTTTCCGAACATATTTCACCCTCCTGCTTGAACGTGGA |
| | GCAAGGCCGGCGATTTACCTTGAAGGGGTTCGTTTAGCTTGCAATTTCATTACGGTTGCTCATGGTTTAACCATGTTGAGTTCTATCTCGCCCATCGACGC |
| | TTACGCATGTTTCTCTCATGGTCTGTTCCTCACCGCAACTGGCAACGGACGAGAGGTTGAAGCCGTGAACGCGACTTTCTGGGATTTAGTGCCATCATTCG |
| | AAGCTGCAAACACAGTGGGGGAGTTGGTTATGTATCACATAAGTCGACTGCATGCTGAAGGAACTCGTTTGTGGAATCGTTCATGGAGGTTCGTGGTTTCA |
| | CCAGACTGCTTGGGTCGTTGCAGCTACGGAGCCTGCTGCCGCAATTGCTTCTTCTACTGGTATGCTAGGGAGTTGTGCATATTTTATTAAGGAAAGGCGTT |
| | CGTTCAGCATCTGATTTCCCATTCTATGCATCGGCTATAATTTCGCGTTGTTTGTAAAGCATATGTTTTCCTGTTATATGCTTCGGCTTTATTAACATGGT |
| | TGTATGCTCTACTCGGTTTGTTGTTATGCTTTGTATTTCCTACTCTTATTTTAACGTTTTTTTATTTCCTTGTTTTATTGTTTATCTTATGAATGGAACTA |
| | AGTTTGTTTACATTATACATATAGTACACCTATTTGAACGTACCAAAACATAATTGTACGTCCATGGAAAGTCATTACAATTAGGCTGTATTTGTAATAAG |
| | TAAACTGTAGCAATTAGAAGGTAAAACTTTAGCATAAAAGTAGGGATATGTTTAAATTCGGTTACAAATCTGTAGTTATTTAGAATAAAAGAAACAATAAT |
| | TAACTAATAGACCACGACTCTGATTTCTTCCACTTCCGGTATATCCGGATTAGCAATGACTTTGGAGATGGGGCCATTGGTCACAGCCTTCAGTTTATGAA |
| | TCCCGGTTAGACGCCAGGATACGAGTGCGATAATGTTTTGAGGACCAGTTAGCCGCTGGAAGAATCTTTGAAATTCGAAAGCTTGGGCATTTTGTGCTTCA |
| | CATTGAAGAACTTCCTTGCTACAGAGAATATATATTATACACAGGATATAACAAACCTAAACCCTAAATAATTAAAAATAAACAACTTATTGTTTATCCAT |
| | TATCTTAAAAGGAAGCTTCCGTCCACGGTAGTCTTCGGTCAAGTTGCCAACCTCAATGAGCACACCAATAACATCTAGTGTACGTAATGGAAGATAAAAGC |
| | ATTATGTGTACGTACATATTTACAAAAATGAGC |
| pNSkb12 | GCTAGGGTGAAGAGCACATGTGTTTTTCCATAGGCTCCGCCCCCCTGACGAGCATCACAAAAATCGACGCTCAAGTCAGAGGTGGCGAAACCCGACAGGAC |
| | TATAAAGATACCAGGCGTTTCCCCCTGGAAGCTCCCTCGTGCGCTCTCCTGTTCCGACCCTGCCGCTTACCGGATACCTGTCCGCCTTTCTCCCTTCGGGA |
| | AGCGTGGCGCTTTCTCATAGCTCACGCTGTAGGTATCTCAGTTCGGTGTAGGTCGTTCGCTCCAAGCTGGGCTGTGTGCACGAACCCCCCGTTCAGCCCGA |
| | CCGCTGCGCCTTATCCGGTAACTATCGTCTTGAGTCCAACCCGGTAAGACACGACTTATCGCCACTGGCAGCAGCCACTGGTAACAGGATTAGCAGAGCGA |
| | GGTATGTAGGCGGTGCTACAGAGTTCTTGAAGTGGTGGCCTAACTACGGCTACACTAGAAGAACAGTATTTGGTATCTGCGCTCTGCTGAAGCCAGTTACC |
| | TTCGGAAAAAGAGTTGGTAGCTCTTGATCCGGCAAACAAACCACCGCTGGTAGCGGTGGTTTTTTTGTTTGCAAGCAGCAGATTACGCGCAGAAAAAAAGG |
| | ATCTCAAGAAGATCCTTTGATCTTTTCTACGGGGTCTGACGCTCAGTGGAACGAAAACTCACGTTAAGGGATTTTGGTCATGAGATTATCAAAAAGGATCT |
| | TCACCTAGATCCTTTTAAATTAAAAATGAAGTTTTAAATCAATCTAAAGTATATATGAGTAAACTTGGTCTGACAGTTACCAATGCTTAATCAGTGAGGCA |
| | CCTATCTCAGCGATCTGTCTATTTCGTTCATCCATAGTTGCCTGACTCCCCGTCGTGTAGATAACTACGATACGGGAGGGCTTACCATCTGGCCCCAGTGC |
| | TGCAATGATACCGCGAGACCCACGCTCACCGGCTCCAGATTTATCAGCAATAAACCAGCCAGCCGGAAGGGCCGAGCGCAGAAGTGGTCCTGCAACTTTAT |
| | CCGCCTCCATCCAGTCTATTAATTGTTGCCGGGAAGCTAGAGTAAGTAGTTCGCCAGTTAATAGTTTGCGCAACGTTGTTGCCATTGCTACAGGCATCGTG |
| | GTGTCACGCTCGTCGTTTGGTATGGCTTCATTCAGCTCCGGTTCCCAACGATCAAGGCGAGTTACATGATCCCCCATGTTGTGCAAAAAAGCGGTTAGCTC |
| | CTTCGGTCCTCCGATCGTTGTCAGAAGTAAGTTGGCCGCAGTGTTATCACTCATGGTTATGGCAGCACTGCATAATTCTCTTACTGTCATGCCATCCGTAA |

GATGCTTTTCTGTGACTGGTGAGTACTCAACCAAGTCATTCTGAGAATAGTGTATGCGGCGACCGAGTTGCTCTTGCCCGGCGTCAATACGGGATAATACC
GCGCCACATAGCAGAACTTTAAAAGTGCTCATCATTGGAAAACGTTCTTCGGGGCGAAAACTCTCAAGGATCTTACCGCTGTTGAGATCCAGTTCGATGTA
ACCCACTCGTGCACCCAACTGATCTTCAGCATCTTTTACTTTCACCAGCGTTTCTGGGTGAGCAAAAACAGGAAGGCAAAATGCCGCAAAAAAGGGAATAA
GGGCGACACGGAAATGTTGAATACTCATACTCTTCCTTTTTCAATATTATTGAAGCATTTATCAGGGTTATTGTCTCATGAGCGGATACATATTTGAATGT
ATTTAGAAAAATAAACAAATAGGGGTTCCGCGCACATTTCCCCGAAAAGTGCCACCTGACGTCTAAGAAACCATTATTATCACCCTTTTCATGATGACCCT
TTTCAACTCTATCCTTTTGTTTTATCCATTTTGTTTCTACCCTCTTTAAATTTTTAATGACCATTTTACCCTTATTTGGAAATTGTTTTAGATTAACAAAA
TGAAATATTAATATTTTTCCGCCTAAAATATTCCAAAATAAATTTCCCGCCAAAAATTTTCAAAAAAAAAATTCCGCCAAAAATTTTCAAAAAACAAATTC
CCGCCAAATTTTGTTTTCAAAAATATTTTTCCCGCCAATTTTTTTTTTGGCGGGAAATAGATTTACAAGGGATTTTTAAAGAGTTTTAAAAGATTTACAAG
AGATTTTAAAGGGGTTTTATAAGATTTACAAGAGATTTTAAAAGGGTGATTTACAAGGGATTTTAAAAGGTTTTTAAAATATTTACAAGAGATTTTAAAAG
GGTTTTAAAAGATTTACAAGAGATTTTAAAGAGATTTAAAAGATTTACAAAGGTCTTTTAAAGATTTTTAAACGGTTTAAAATTATTTTAAAATGATTTTA
AAATATTTACAAGGGATTTTAAAAGGGTTTTACAATATTTACAACAGATTTTAAAGGACTTTAAAATATTTACAAAAGATCTGTAAAGATTCTTAAAGGGT
TTAAAAGTGTTTTGAAAGATTTACAAGAGATTTTAAAACCTTTTAAAATTACTTGTAAATTTTTTAAAACCCTTGTAAATCTTTTAAAATCTCTTTTAAAT
CTTTTAAAACCTTTTTAAAATCTCTTGTAAATCTTTTAAGACCCTTTTAAAATCCCTTGTAAATCTATTTGCCGCAAACAAATATTTACAAGGATTTTTAA
AATTATTTTTTTGGCTCAGAAAAAAATTTGTTACGGAACAAAAAATTTGGCGGGAAACTTTTTTTTTGGCGGGAAAAAAGTTCGGCGGGAAATCTTTTTTGG
CGGGAAAAAAAAAATTTTGGCGCCAATAAATTTGGCGGGAAAAGTTCAGTTTCCAGCATCTCTCTATTCGCTATTCCCAGATCTCCATGACCCACTTCACTC
CCTGCGCGTCCGGTACAATCGCCACTTCCATAACACCAAGAGTCAATACAAAAAAAAAAGCCGACTCCATTATTATCCCTCTCTCTACAAAAAAAAACAGTAG
AGAGAGATATAGAGAGCATCAATGGCGTCGTCGTCAAGACTTTGCGATTCGTGCAAATCGACGGCAGCAACTCTATTCTGCCGCGCAGACGCCGCGTTTCT
CTGCGGCGACTGCGACGGTAAGATCCACACAGCTAACAAACTCGCATCACGTCACGAACGAGTTTGGCTATGTGAAGTATGCGAACAAGCACCAGCACACG
TCACGTGCAAAGCTGACGCCGCTGCGTTATGCGTCACGTGCGACCGTGATATCCACTCAGCTAATCCACTCTCTCGCCGCCACGAGCGCGTACCAATCACT
CCTTTCTACGACGCTGTAGGTCCAGCAAAATCCGCTTCCTCCTCCGTCAATTTCGTAGATGAAGATGGCGGTGACGTCACTGCTTCGTGGCTTTTAGCTAA
AGAAGGAATCGAAATCACTAATTTGTTCTCCGATCTTGATTATCCGAAGATTGAGGTAACGTCGGAGGAGAATAGCTCCGGTAACGACGGAGTTGTTCCTG
TGCAGAATAAGTTGTTTCTTAACGAAGATTACTTCAATTTCGATCTCTCTGCTTCCAAAATTTCTCAACAAGGATTCAATTTCATCAACCAAACTGTACGA
ATCTCTCTCTATCTTCTTAAGCTTCCATGGATCTGGAAAATTTTCAGATCTAGATCTGAAAGTTTGGATCTGATTTGAGTTTGTGAAATGTTGTTAGGTTT
CGACGAGAACGATAGATGTACCGTTGGTGCCTGAAAGTGGAGGAGTGACGGCGGAGATGACGAACACGGAGACGCCAGCTGTGCAGTTATCACCGGCGGAG
AGGGAAGCAAGGGTTTTGAGGTATAGAGAGAAGAGGAAGAATCGGAAATTTGAGAAGACGATTAGGTATGCGTCGCGTAAAGCTTACGCTGAGATGAGGCC
GAGAATCAAAGGACGTTTTGCTAAGAGAACAGATTCGAGAGAGAATGATGGTGGAGACGTCGGAGTTTATGGCGGATTCGGTGTTGTTCCGAGTTTCTGAT
ATTTTCCGGTGAAAGAAACATGGTAGTAGTATATCGGTTAATAGCAGATTAAGGATAATTAAAACTAAAATGTTTCCTGATTTAATCAGGGGCTAAGTTAT
TGTTAGAAGATTGTTGTTTTTTGAATTAATTCTCACAATTGTATCCTAAATTATATAGTTTACTTAATGTTGAACTAATAAATTTTGATTTACAATCTCTG
TAAGAAATCGATATGCAAAAATAAATGAGTTGAGCTATTAAGATTAACATTCATATAGAACACAAACTCATACAATCAGCTGTGGTGGGAAAAAAAGCATT
ATAATCCTTATATTACTAGTGAGTAGTGACTGATACAAAAGATTTTTTATACGTTAAAAGTTTAAAAATTCAAAAGTACAGAAAACGTTATATTCGCTTGT
TACCTTTTTTAACTTTATCCACGTCATTACCCCTGAGAGTAGCCCAACGAAAAATGGAAAATTTAACCCAAAAAACGGAAAATATTAGAGAAAGCGTTAAT
GCCTTTTCTAACAACTAAACGACGACGTATCAGTTCAATCCTCTCTCTATTATTATTCCTTAAAACTTAACTCTAAAGCTTTCGAGTATTCTTCAATCTCT
TCTTCTCCGATCGATCTGAAGTAAGAGAGCTTCTTAAGGAAGAAGAAGATGGGTTGTGCTCAATCAAAGATCGAGAACGAAGAAGCAGTTACTCGTTGCAA
AGAACGAAAACAATTGATGAAAGACGCCGTCACTGCTCGTAACGCTTTCGCCGCCGCTCACTCAGCTTACGCTATGGCTCTTAAAAACACCGGAGCTGCTC
TTTCCGATTACTCTCACGGCGAGTTTTTAGTCTCTAATCACTCGTCTTCCTCCGCAGCTGCAGCAATCGCTTCTACTTCTTCTCTTCCCACTGCTATATCT
CCTCCTCTTCCTTCTTCCACCGCTCCGGTTTCTAATTCAACCGCTTCTTCTTCCTCCGCTGCGGTTCCTCAGCCGATTCCTGATACTCTTCCTCCTCCTCC
TCCTCCACCACCGCTTCCTCTTCAACGTGCTGCTACTATGCCGGAGATGAACGGTAGATCCGGTGGTGGTCATGCTGGTAGTGGACTCAACGGAATTGAAG
AAGATGGAGCCCTAGATAACGATGATGATGACGATGATGATGATGATGACTCTGAAATGGAGAATCGTGATCGTTTGATTAGGAAATCGAGAAGCCGTGGA
GGTAGTACTAGAGGAAATAGGACGACGATTGAAGATCATCATCTTCAGGAGGAGAAAGCTCCGCCACCTCCCCCTTTGGCGAATTCACGGCCAATTCCGCC
GCCACGTCAGCATCAGCATCAACATCAGCAACAGCAACAACAACCTTTCTACGATTACTTCTTCCCTAATGTTGAGAATATGCCTGGAACTACTTTAGAAG
ATACTCCTCCACAACCACAACCACAACCAACAAGGCCTGTGCCTCCTCAACCACATTCACCAGTCGTTACTGAGGATGACGAAGATGAGGAGGAGGAAGAG
GAGGAAGAGGAGGAGGAAGAGGAGACGGTGATTGAACGGAAACCACTGGTGGAGGAAAGACCGAAGAGAGTAGAGGAAGTGACGATTGAATTGGAAAAAGT
TACTAATTTGAGAGGGATGAAGAAGAGTAAAGGGATAGGGATTCCCGGAGAGAGGAGAGGAATGCGAATGCCGGTGACTGCGACGCATTTGGCGAATGTAT
TCATTGAGCTTGATGATAATTTCTTGAAAGCTTCTGAAAGTGCTCATGATGTTTCTAAGATGCTTGAAGCTACTAGGCTCCATTACCATTCTAATTTTGCA
GATAACCGAGGTAATTTTGATTCGATAAATCTTTATCCTTTTTCTTTTGATGCTTCATTCTCTCTTTTCTTTTTTTTTCCTCTGTAGCGTTTCTAGGAATCT
CTCTCCAACGTTTAGGTTTCTTCTCAGATTTAGAAAGTTTCTTCTCTATCTGAAGAATTTTCATTTCGATTTCGCTTCTTCCTTCACCCTAATCCGTAATC
GAATGCCATTTCGAAGATTGTGTTGAAGAGTAGAGAGGAAGAAGAAGACTAAGAAGCTAGGGTTTCTCTGATGAGTAGATGATGCGTGAACCGCTTGTTAG
CGAAGAAGAAGAAGAAGAAGCTACTGAAGTTTTACTAGTGGAGAAGACGAAGTTGTGTAAGAGACGAGGAGACGAAGAAAAAACGGAAGAGAGAAGAGACG
ATTTGTTGTTACTAGCTTTAACGCCGATGGTACGATCGAAATCTCAAGGAACTACTCGGCGCGTCACTCCCACGCCACCTCCTGTTGACGTGGAGAAACCG
TTACCAAACGGAGATCTCTACATGGGAACGTTTTCCGGCGGGTTTCCCAACGGATCCGGTAAGTATTTGTGGAAGGATGGGTGTATGTACGAAGGCGAGTG
GAAACGTGGTAAAGCGAGTGGTAAAGGCAAGTTTTCGTGGCCGAGTGGTGCGACTTATGAAGGAGAGTTCAAATCTGGGAGAATGGAAGGATCTGGGACTT
TTGTTGGTGTTGATGGTGATACTTATCGTGGCTCTTGGGTTGCTGATCGGAAACAAGGTCATGGTCAGAAGAGATATGCCAACGGAGATTACTATGAAGGT
ACATGGCGGCGGAATCTTCAGGATGGGAGAGGGAGATATGTTTGGATGAATGGGAATCAGTATACAGGAGAGTGGAGAAATGGTGTGATATGTGGTAAAGG
TGTGCTTGCTTGGCCTAATGGGAATAGATATGAAGGTCAATGGGAAAATGGTGTTCCTAAAGGAAGTGGTGTGTTTACTTGGGCTGATGGAAGTTCGTGGA
TTGGTTCTTGGAATGAGAGTAGTAATCTCATGAGGAATTTCTTTGATGGGATTGAGAAGAATGAGTTGATTGTTGCGACTAGGAAGAGATCTTCGGTTGAT
AGTGGCGCTGGAAGTTTGACTGGGGAGAAGATTTTCCCTAGGATATGTATTTGGGAGTCTGATGGAGAAGCTGGGGATATTACTTGTGATATTGTTGATAA
TGTGGAAGCTTCTGTGATATACAGAGATAGGATTTCTATTGATAAAGATGGGTTTCGTCAGTTTAGGAAGAATCCTTGTTGTTTCAGCGGTGAGGCTAAGA
AACCTGGAGAGACGATATCTAAAGGGCATAAGAAATATGATTTGATGCTCAACCTGCAGCATGGAATTAGGTAACTTCTTAGAAGCTTGAGGATCCCAATG
TTTGGAGAGTTTCTGGATCTGTGTTTTTCTAACCAAGTTGTGAATTTTTAGGTACTCTATTCATTTTTGTAAATATGTACGTACACATAATGCTTTTATCT
TCCATTACGTACACTAGATGTTATTGGTGCGCTCATTGAGGTTGGCAACTTGACCGAAGACTACCGTGGACGGAAGCTTCCTTTTAAGATAATGGATAAAC
AGTAAGTTGTTTATTTTTAATTATTTAGGGTTTAGGGTTTGTTATATCCTGTGTATAATATATATTCTCTGTAGCAAGGAAGTTCTTCAATGTGAAGCACA
AAATGACCAAGCTTTCGAATTTCAAAGATTCTTCCAGCGGCTAACTGGTCCTCAAAACATTATCGCACTCGTATCCTGGCGTCTAACCGGGATTCATAAAC
TGAAGGCTGTGACCAATGGCCCCATCTCCAAAGTCATTGCTAATCCGGATATACCGGAAGTGGAAGAAATCAGAGTCGTCGTCTATTAGTTAATTATTGTT
TCTTTTATTCTAAATAACTACAGATTTGTAACCGAATTTAAACATATCCCTACTTTTATGCTAAAGTTTTACCTTCTAATTGCTACAGTTTACTTATTACA
AATACAGCCTAATTGTAATGACTTTCCATGGACGTACAATTATGTTTTTGTACGTTCAAATAGGTGTACTAGATGTATAATGTAAACAAACTTAGTTCCAT
TCATAAGATAAACAATAAAACAAGGAAATAAAAAAACGTTAAAATAAGAGTAGGAAATACAAAGCATAACAACAAACCGAGTAGAGCATACAACCATGTTA
ATAAAGCCGAAGCATATAACAGGAAAACATATGCTTTACAAACAACACGAAATTATAGCCGATGCATAGAATGGGAAATCAGATGCTGAACGAACGCCTTT
CCTTAATAAAATATGCACAACTCCCTAGCATACCAGTAGAAGAAGCAATTGCGGCAGCAGGCTCCGTAGCTGCAACGACCCAAGCAGTCTGGTGAAACCAC
GAACCTCCATGAACGATTCCACAAACGAGTTCCTTCAGCATGCAGTCGACTTATGTGATACATAACCAACTCCCCCACTGTGTTTGCAGCTTCGAATGATG
GCACTAAATCCCAGAAAGTCGCGTTCACGGCTTCAACCTCTCGTCCGTTGCCAGTTGCGGTGAGGAACAGACCATGAGAGAAACATGCGTAAGCGTCGATG
GGCGAGATAGAACTCAACATGGTTAAACCATGAGCAACCGTAACGAAATTGCAAGCTAAACGAACCCCTTCAAGGTAAATCGCCGGCCTTGCTCCACGTTC
AAGCAGGAGGGTGAAATATGTTCGGAAAGGTGTTTCCGGCAACACGAAATCAGGCCACGTCATAAACTCATCCATGTTGATAGCGGTCGGCTCCAGCGTGC
TAACGAGACCTCTGAGTGTTTCAATGGGAGTTACGGAAACGGGTAAGTAGTTGACGAGGATGCTGAGTGGGTTAGGTAGCTGAATGGCCAGTAAATTTCCG
CCGGTGGGGGTGTGGGGACGTCCATTGCCGAGACTGAGGGATTTATTTGTGAGTTTAGGTTATATATGGCGTAGTAAGAGCAATGAAAAGTGTGGAGAAGA
TGACTGATTATACCCACTGAAGGGTATTTCCGTCAATCCCAATTCTAACAATGAATTCAGGAGTATAAAAACGTAAATTCAAGCGTGCCAATTATAAACCG
TCGATCATAATCTAATCCAACGGCAGTAACATCGATCCGCGTGATTGTTTATTATTGGATAAGAATCACTCAACCGTCTCTACACAGTATATATAATAACC
AAAGAGCGTCCTCTTACGCTTATCTTAATTTCCCTCGCATTGAGAATTTTCAACTTTTTCTATCTCTCTTCCCAAATCACAAATGGCGGGTCGCGGCAAAA
CTCTCGGATCTGGCGTTGCTAAGAAATCAACATCGAGAAGCAGCAAAGCCGGTCTCCAATTCCCCGTTGGTCGTATCGCTCGTTTTCTAAAGAACGGCAAG

TACGCAACACGTGTTGGTGCCGGAGCTCCGGTTTACTTAGCCGCCGTTCTCGAATACCTCGCCGCTGAGGTAATTATCCCCTTCTCTCCCTATATCTCTTT
ACTCTTTCGATCTTCAATTTCGTAAAACCCTAATTTCTAAATTGGATCTGTTGTGTTGTAGGTATTGGAATTGGCTGGAAACGCAGCTAGGGATAACAAGA
AGACTAGGATTGTGCCACGTCACATTCAGCTCGCGGTGAGAAACGATGAGGAGCTGAGTAAACTGCTTGGAGATGTGACGATTGCTAATGGAGGTGTGATG
CCTAACATTCACAGTCTTCTTCTTCCCAAGAAAGCTGGTGCTTCAAAACCTTCCGCTGATGAAGATTAGATTAGGGATTTGTGTTGTGGTTGTTTAGCTAA
TTAATGTGTAGCTTAGTCTTTCATTAGATTAGATCTGAATTAGTTTTCATTAATGGTGTTGTGTAGTCTCTCTTTTGCTTCAAAAACAAGTATTAAAATCT
TATTATTTTGAATTGAATCCACAATCAATACACATTGAAGTCCTAACAAACTACTTCTTCCCAGTGATATTTGAAACCAAATCACTAAGAAACTTAGCTGA
TTTGGTAATAGGAGAATTCATAGCCATCAAGTTATACAGAACAAGCTCAACTTCTTCGATTGATGGTCGAGAATTGAATTGTGAAACAACTTTCAAAGTAC
CATTACCTTCTTCTTCTTCAACGAGAACATTCCATCTTTCTCCACTCACAACACCGTCTCTTACACACTTAAAGATCACTCCTTTTCTCTTACTAAACACT
CCTCTCACTTCTAAGCCAGTGAACGCATAAATCACTTCAAAGGAACCAATAAAGCCATCTCCAAAGCTTTTCTCTTCTTCAAACAACCATCTCGGATTGAT
CATTACTACAGGTTTTGTAGAAAAAGCTTCACTCGCGGTCTTAACATCCTCCAGCTGCGATTTCTCCGGAGCCATGAAGATCGCCACGTCAGCAGATCTCA
GAGCTGTTTTAAGAGAGTCTGAGACCGGAAGTCATCTTCTCCACACTTTTCATTGCTCTTACTACGCCATATATAACCTAAACTCACAAATAAATCCCTCA
GTCTCGGCAATGGACGTCCCCACACCCCCACCGGCGGAAATTTACCTCGCCATTCAGCTACCTAACCCACTCAGCATCCTCGTCAACTACTTACCCGTTTC
CGTAACTCCCATTGAAACACTCAGAGGTCTCGTTAGCACGCTGGAGCCGACCGCTATCAACATGGATGAGTTTATGACGTGGCCTGATTTCGTGTTGCCGG
AAACACCTTTCCGAACATATTTCACCCTCCTGCTTGAACGTGGAGCAAGGCCGGCGATTTACCTTGAAGGGGTTCGTTTAGCTTGCAATTTCATTACGGTT
GCTCATGGTTTAACCATGTTGAGTTCTATCTCGCCCATCGACGCTTACGCATGTTTCTCTCATGGTCTGTTCCTCACCGCAACTGGCAACGGACGAGAGGT
TGAAGCCGTGAACGCGACTTTCTGGGATTTAGTGCCATCATTCGAAGCTGCAAACACAGTGGGGGAGTTGGTTATGTATCACATAAGTCGACTGCATGCTG
AAGGAACTCGTTTGTGGAATCGTTCATGGAGGTTCGTGGTTTCACCAGACTGCTTGGGTCGTTGCAGCTACGGAGCCTGCTGCCGCAATTGCTTCTTCTAC
TGGTATGCTAGGGAGTTGTGCATATTTTATTAAGGAAAGGCGTTCGTTCAGCATCTGATTTCCCATTCTATGCATCGGCTATAATTTCGCGTTGTTTGTAA
AGCATATGTTTTCCTGTTATATGCTTCGGCTTTATTAACATGGTTGTATGCTCTACTCGGTTTGTTGTTATGCTTTGTATTTCCTACTCTTATTTTAACGT
TTTTTTATTTCCTTGTTTTATTGTTTATCTTATGAATGGAACTAAGTTTGTTTACATTATACATATAGTACACCTATTTGAACGTACCAAAACATAATTGT
ACGTCCATGGAAAGTCATTACAATTAGGCTGTATTTGTAATAAGTAAACTGTAGCAATTAGAAGGTAAAACTTTAGCATAAAAGTAGGGATATGTTTAAAT
TCGGTTACAAATCTGTAGTTATTTAGAATAAAAGAAACAATAATTAACTAATAGACCACGACTCTGATTTCTTCCACTTCCGGTATATCCGGATTAGCAAT
GACTTTGGAGATGGGGCCATTGGTCACAGCCTTCAGTTTATGAATCCCGGTTAGACGCCAGGATACGAGTGCGATAATGTTTTGAGGACCAGTTAGCCGCT
GGAAGAATCTTTGAAATTCGAAAGCTTGGGCATTTTGTGCTTCACATTGAAGAACTTCCTTGCTACAGAGAATATATATTATACACAGGATATAACAAACC
TAAACCCTAAATAATTAAAAATAAACAACTTATTGTTTATCCATTATCTTAAAAGGAAGCTTCCGTCCACGGTAGTCTTCGGTCAAGTTGCCAACCTCAAT
GAGCACACCAATAACATCTAGTGTACGTAATGGAAGATAAAAGCATTATGTGTACGTACATATTTACAAAAATGAGC

**Movie S1. MpARF2 nanoclusters inside the nucleus at native expression**

Time-lapse confocal microscopy showing *MpArf2-mNeonGreen* assemblies in the nucleus of a rhizoid precursor cell, with native expression level. The cell is located in the gemma of a six-week-old *Marchantia polymorpha* plant. The image data were corrected to compensate for photobleaching during image acquisition.

**Movie S2. Coalescence of MpARF2 condensates**

Time-lapse confocal microscopy of *MpARF2-mNeonGreen* condensates coalescing on a coverslip surface at a protein concentration of 600 nM.

**Movie S3. Single-molecule assay of a DNA-bound nanocluster**

Microfluidic assay showing 3IR7kb12 DNA (green) as template and Atto633-labeled *MpARF2* protein (magenta) at 5 nM concentration. Bound DNA–protein complexes move with the DNA, while protein adsorbed to the microfluidic chip surface remains stationary. A slight temporal offset between channels arises from alternating acquisition of green and magenta signals. At the end of the movie, flow is stopped, causing the DNA strand to coil.
